# Human pancreatic islet 3D chromatin architecture provides insights into the genetics of type 2 diabetes

**DOI:** 10.1101/400291

**Authors:** Irene Miguel-Escalada, Silvia Bonàs-Guarch, Inês Cebola, Ponsa-Cobas Joan, Julen Mendieta-Esteban, Delphine M.Y. Rolando, Biola M. Javierre, Goutham Atla, Irene Farabella, Claire C. Morgan, Javier García-Hurtado, Anthony Beucher, Ignasi Morán, Lorenzo Pasquali, Mireia Ramos, Emil V.R. Appel, Allan Linneberg, Anette P. Gjesing, Daniel R. Witte, Oluf Pedersen, Niels Grarup, Philippe Ravassard, David Torrents, Josep Maria Mercader, Lorenzo Piemonti, Thierry Berney, Eelco J.P. Koning de, Julie Kerr-Conte, François Pattou, Iryna O. Fedko, Inga Prokopenko, Torben Hansen, Marc A. Marti-Renom, Peter Fraser, Jorge Ferrer

## Abstract

Genetic studies promise to provide insight into the molecular mechanisms underlying type 2 diabetes (T2D). Variants associated with T2D are often located in tissue-specific enhancer regions (enhancer clusters, stretch enhancers or super-enhancers). So far, such domains have been defined through clustering of enhancers in linear genome maps rather than in 3D-space. Furthermore, their target genes are generally unknown. We have now created promoter capture Hi-C maps in human pancreatic islets. This linked diabetes-associated enhancers with their target genes, often located hundreds of kilobases away. It further revealed sets of islet enhancers, super-enhancers and active promoters that form 3D higher-order hubs, some of which show coordinated glucose-dependent activity. Hub genetic variants impact the heritability of insulin secretion, and help identify individuals in whom genetic variation of islet function is important for T2D. Human islet 3D chromatin architecture thus provides a framework for interpretation of T2D GWAS signals.

## Introduction

Protein-coding mutations account for most known causes of Mendelian disease, yet fail to explain the inherited basis of most polygenic disorders. Recently, hundreds of thousands of transcriptional enhancers have been mapped to the human genome, holding promise to decipher disease-causing noncoding genetic variants ^1-3^. A standing challenge is to connect distal enhancers with the target genes that mediate disease-relevant cellular phenotypes.

Type 2 diabetes (T2D) affects more than 400 million people worldwide ^4^, and is a classic example of a polygenic disease in which noncoding sequence variation plays a central predisposing role ^5^. Several lines of evidence implicate noncoding sequence variation in the pathogenesis of T2D, such as the lack of plausible protein-coding causal variants at most associated loci ^6^, and the enrichment of associated variants in pancreatic islet enhancers or eQTLs ^7-11^. T2D susceptibility variants are particularly enriched in active islet enhancer clusters–variably defined as super-enhancers, COREs, enhancer clusters, or stretch enhancers ^7,8,12,13^. Enhancer clusters in other cell types are also enriched in risk variants for common diseases, and have been shown to harbor somatic oncogenic mutations ^7,8,14-19^. Despite the importance of enhancer clusters, several pivotal questions remain unaddressed. First, enhancers can loop over long distances to gain proximity with their target genes. This warrants a need to link enhancer clusters to their target genes based on their spatial proximity in disease-relevant cells. Second, enhancer clusters have so far been defined with unidimensional epigenome maps, which do not necessarily reflect the capacity of enhancers to cluster in three-dimensional (3D) nuclear space. A definition of enhancer clusters based on 3D proximity could thus provide more relevant definitions of genomic spaces that underlie human disease.

Here, we describe a chromatin interactome map of human pancreatic islets. Because most enhancer-based long-range chromatin interactions are not captured by untargeted Hi-C methods, we used promoter capture Hi-C (pcHi-C) ^20^, an approach that enables high-resolution mapping of interactions between promoters and their regulatory elements. Using genome and epigenome editing we validated predicted gene targets of T2D-associated islet enhancers. We further describe hundreds of tissue-specific enhancer hubs formed by groups of enhancers that cluster with key islet gene promoters in a restricted 3D space. Finally, we demonstrate that islet hub enhancers have the potential to stratify individuals in whom T2D susceptibility is predominantly driven by pancreatic islet regulatory variants.

## Results

### The promoter interactome of human islets

To create a genome-wide map of long-range interactions between gene promoters and distant regulatory elements in human pancreatic islets, we prepared Hi-C libraries from four human islet samples. We performed hybridization capture of 31,253 promoter-containing HindIII fragment baits and their ligated DNA fragments, which were then sequenced and processed with the CHiCAGO algorithm to define promoter-interacting DNA fragments ^20,21^ (Figure 1a,b, **Supplementary 1a-c**). This resulted in 175,784 high-confidence interactions (CHiCAGO score > 5) between annotated promoters and distal genomic regions. We independently validated pcHi-C landscapes at two loci by 4C-seq analysis in the EndoC-βH1 human β cell line ^22^ (Supplementary Figure 1j,k).

**Figure 1.**
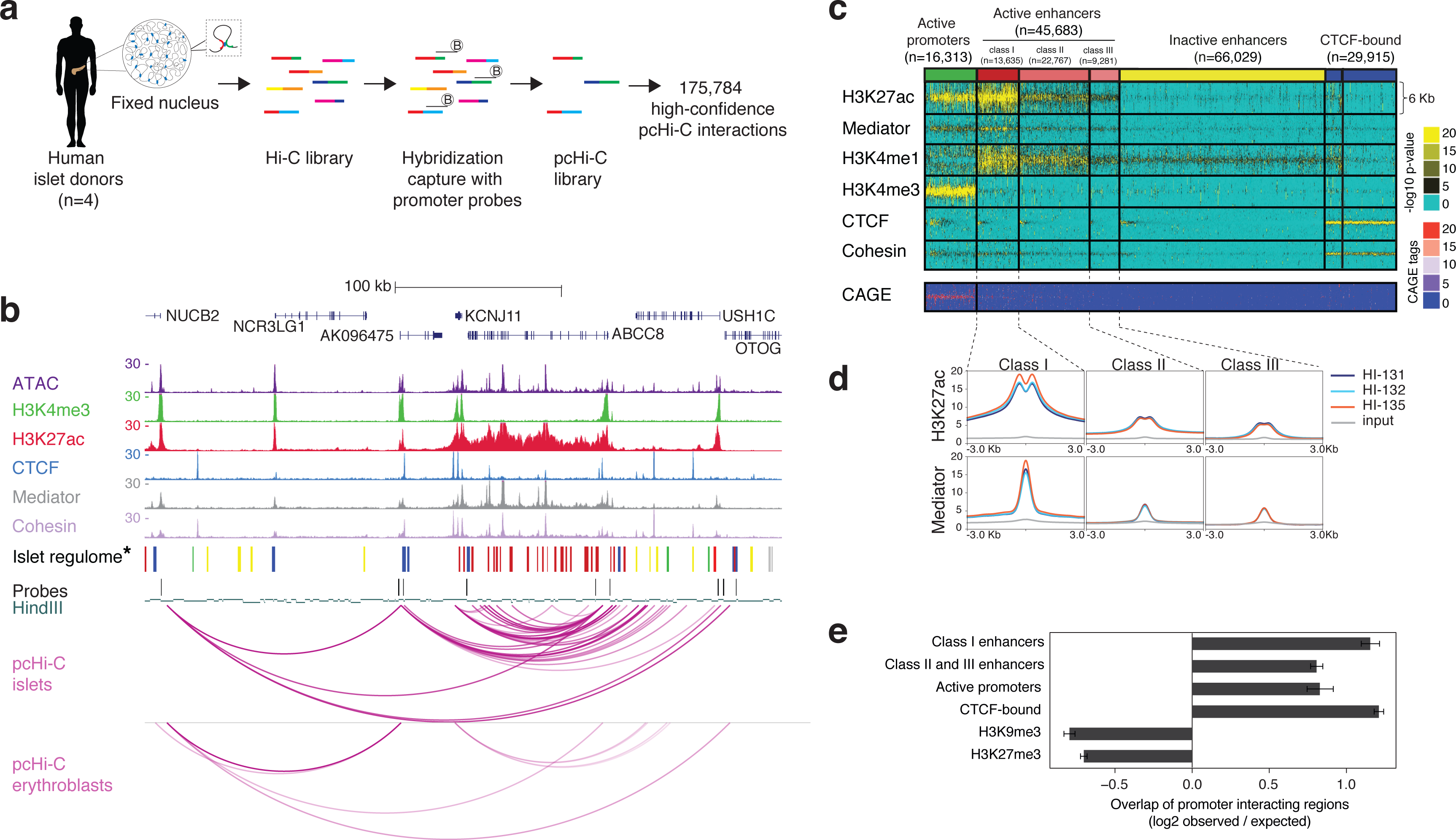
The promoter interactome of human pancreatic islets. (a) Overview of promoter-capture Hi-C (pcHi-C) in human islets. (b) Integrative map of the *KCNJ11-ABCC8* locus, which encode two subunits of the islet K_ATP_ channel, showing human islet ATAC-seq and ChIP-seq profiles, HindIII fragments used as baits, and arcs representing high-confidence pcHi-C interactions in human islets and erythroblasts. **\***Color code for islet regulome is as shown in panel c. (c) High-resolution annotations of islet open chromatin regions. ATAC-seq data from 13 islet samples was used to define consistent open chromatin regions, which were then classified with k-medians clustering based on additional epigenomic features. Mediator and H3K27ac binding patterns allowed sub-classification of active enhancer classes. Post-hoc analysis of islet CAGE tags confirms that transcription start sites are highly enriched in promoters and weakly in class I enhancers. These annotations are hereafter referred to as the islet regulome, and represented with indicated color codes. The genomic locations are provided in **Data S1.1**. (d) Average H3K27ac and Mediator signal distribution centered on open chromatin regions for each active enhancer subtype in three human islet (HI) samples and input DNA. (e) Overlap of promoter-interacting regions with epigenomic features, expressed as a ratio over the median of a null distribution calculated with overlaps obtained with 100 sets of fragments matched for distance relative to bait fragments. Bars represent mean log2 ratios and error bars represent 95% confidence intervals. (f) Ratio of tissue-invariant to islet-selective interactions overlapping with major open chromatin classes, normalized by the total number of tissue-invariant and islet-selective interactions. All categories showed significant differences with interactions in the remaining genome (Fisher’s P < 0.01).

In parallel, we created new human islet ChIP-seq and ATAC-seq datasets, and refined human islet epigenome annotations to subclassify active enhancers according to Mediator, cohesin, and H3K27ac occupancy patterns (Figure 1b-d, **and Extended Data 1.1**), which were then integrated them with 3D chromatin maps. This showed that, expectedly, promoter-interacting genomic regions were markedly enriched in active enhancers, promoters, and CTCF-bound regions (Figure 1b,e, Supplementary Figure 1d-f). Islet interacting regions that were also observed in distant cell types were enriched in CTCF-binding sites and active promoters, whereas islet-selective interacting regions were enriched in active enhancers (particularly enhancers with strong Mediator occupancy, which we term class I enhancers) and islet-specific promoters (Supplementary Figure 1g-i). This genome-scale map of the human pancreatic islet promoter interactome is accessible for visualization along with pcHi-C maps of other human tissues (www.chicp.org) ^23^, or as virtual 4C representations for all genes along with other islet regulatory annotations (http://isletregulome.org/regulomebeta/) ^24^.

### Identification of target genes for pancreatic islet enhancers

Long-range chromatin interactions are largely constrained within topologically associating domains (TADs), which typically span hundreds of kilobases and are often invariant across tissues ^25,26^ (Supplementary Figure 2a-e). TADs, however, define broad genomic spaces that do not inform on the specific interactions that take place in each tissue between individual *cis*-regulatory elements and their target genes. Human islet pcHi-C maps identified high-confidence pcHi-C interactions (CHiCAGO score > 5) between gene promoters and 18,031 different islet enhancers, comprising ∼40% of all annotated islet enhancers (Figure 2a).

**Figure 2.**
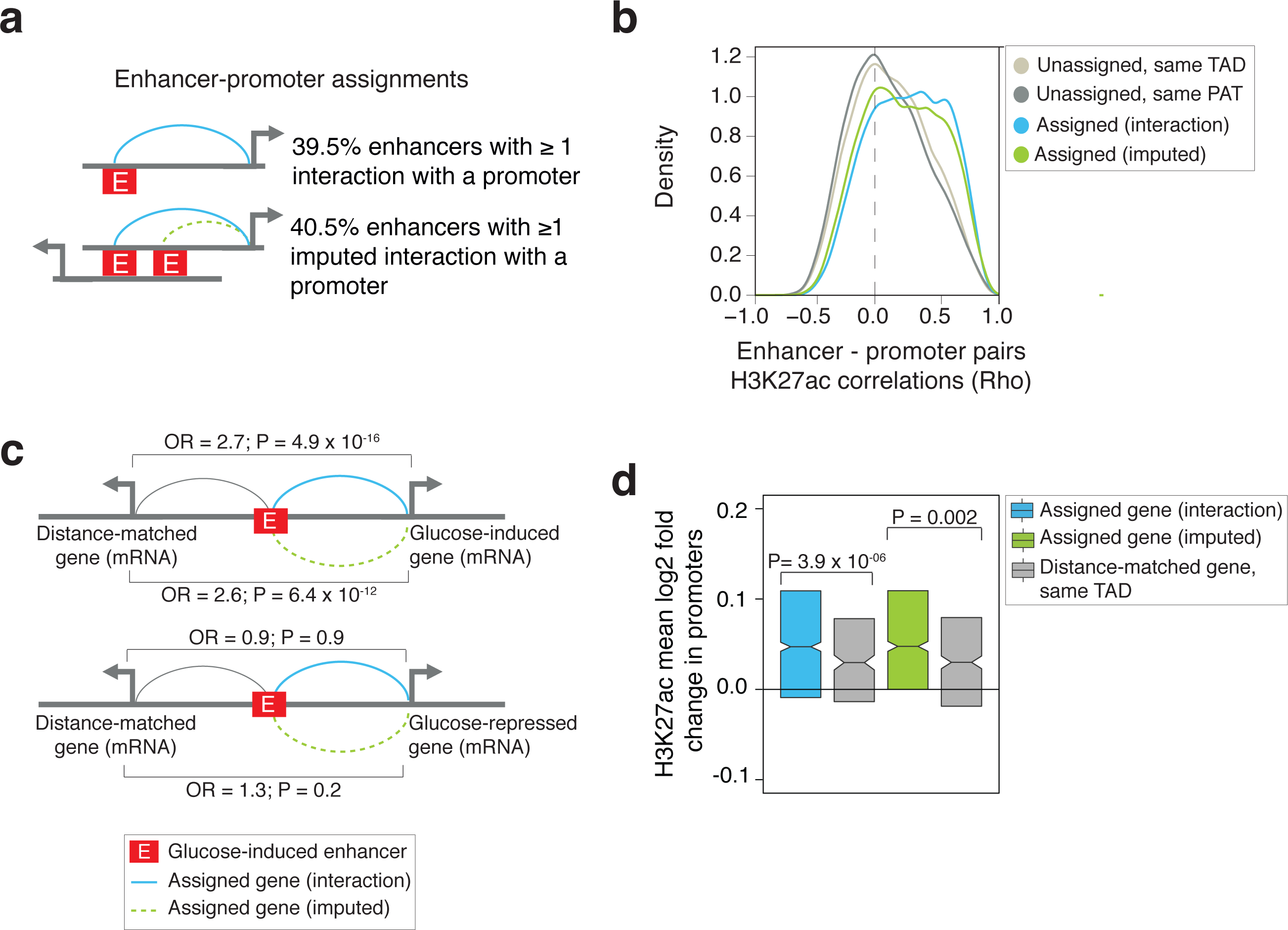
Identification of target genes of pancreatic islet enhancers. (a) pcHi-C maps allowed the assignment of target genes to 39.5% of all 45,683 active enhancers through high-confidence interactions. PAT features allowed imputing the assignment of promoters to another 40% of all active enhancers (see also Supplementary Figure 2k, and evidence that imputations are enriched in lower confidence interactions in Supplementary Figure 2l). (b) Functional correlation of assigned enhancer-gene pairs. Spearman’s Rho values for normalized H3K27ac signal in enhancer-promoter pairs across 14 human islet samples and 51 Roadmap Epigenomics tissues. Rho values are shown for enhancer-gene pairs assigned through high-confidence interactions or imputations, and control enhancer-gene pairs in which the enhancer overlapped a PAT in linear maps but was not assigned to the PAT promoter, or other unassigned gene-enhancer pairs from the same TAD. (c) Genes assigned to glucose-induced enhancers show concordant glucose-induced expression. Top: glucose-induced enhancers showed enriched high-confidence or imputed assignments to glucose-induced genes, compared with distance-matched genes from the same TAD. Bottom: glucose-induced enhancers showed no enrichment for assignments to genes that were inhibited by high glucose concentrations. OR = odds ratio. P values were calculated with Chi-square tests. (d) Genes assigned to glucose-induced enhancers through high-confidence interactions or imputations showed increased glucose-induced changes in promoter H3K27ac, compared with other genes from the same TAD as the enhancer. Box plots represent IQRs, and notches are 95% confidence intervals of median. P values were calculated with Wilcoxon’s signed ranked test. See also **Data S1.2**.

High-confidence interactions between enhancers and their target genes can be missed due the high stringency of detection thresholds, the strong bias of Hi-C methods against detection of proximal interactions, or the likely dependence of some interactions on specific physiological conditions. We thus used pcHi-C data to impute additional enhancer-promoter assignments. To this end, we considered promoter-associated three-dimensional spaces (PATs). A PAT is defined as the space that contains all pcHi-C interactions that stem from a promoter bait (Supplementary Figure 2f-i). We observed that PATs that have a high-confidence interaction with an enhancer exhibit features that distinguish them from other PATs. For example, they tend to show other enhancer-promoter interactions. We thus used PAT features to impute plausible target promoter(s) of an additional 18,633 islet enhancers that did not show high-confidence interactions (Figure 2a; see also Supplementary Figure 2k and **Methods** for a detailed description of the imputation pipeline). The distribution of CHiCAGO scores of imputed promoter-enhancer pairs showed a significant shift toward higher values compared with the same promoters paired with unassigned enhancers from the same PATs or TADs (Kruskall-Wallis P < 10^−323^; Supplementary Figure 2l). In total, we assigned 36,664 human islet active enhancers (80% of all enhancers) to at least one candidate gene (Figure 2a, **Extended Data 1.2**).

To validate assignments, we calculated normalized H3K27ac signals in assigned enhancer-promoter pairs across human tissues and human islet samples, and found distinctly higher correlation values than for pairs of enhancers and distance-matched promoters from the same TAD or PAT (Figure 2b). Importantly, this was true for both high-confidence interacting and imputed assignments (Figure 2b). Furthermore, islet-selective expression was enriched in enhancer-assigned genes but not in unassigned genes from the same TAD, as expected (Supplementary Figure 2m).

To further test enhancer-promoter assignments, we used a dynamic perturbation model. We exposed human islets from 7 donors to moderately low (4 mM) or high (11 mM) glucose for 72 hours, which correspond to quasi-physiological glucose concentrations. This led to significant glucose-dependent changes in H3K27ac levels (at adjusted P < 0.05) in 3,850 enhancers, most of which showed increased activity at high glucose concentrations (Supplementary Figure 2n). This indicates that changes in glucose concentrations elicit global quantitative changes in enhancer activity in pancreatic islets. We predicted that if glucose-regulated enhancers cause glucose-regulated expression of their target genes, this should be reflected in our enhancer-promoter assignments. Indeed, glucose-induced enhancers were preferentially assigned to genes that showed glucose-induced mRNA levels, compared with distance-matched actively transcribed control genes from the same TAD (odds ratio 2.7 and 2.6, Fisher’s P = 4.9 × 10^−16^ and 6.4 × 10^−12^, for high-confidence or imputed assignments, respectively) (Figure 2c). By contrast, glucose-induced enhancers were not preferentially assigned to glucose-inhibited genes (Figure 2c). Likewise, genes assigned to glucose-induced enhancers showed significantly greater glucose-induction of promoter H3K27ac than distance-matched promoters in the same TAD (Figure 2d). Taken together, these findings indicate that pcHi-C identifies functional target genes of transcriptional enhancers in human pancreatic islets.

### Identification of transcriptional targets of T2D-relevant enhancers

A fundamental challenge to translate GWAS data into biological knowledge is that the target genes of disease-associated regulatory variants are often unknown. To link noncoding variants to their target genes, we compiled T2D and/or fasting glycemia (FG) associated variants from 109 loci, most of which had been fine-mapped to a credible variant set (Supplementary Figure 3a, **Extended Data 1.3** and **Methods**). We identified 62 loci that contain candidate causal T2D and/or FG-associated variants that overlap islet enhancers. For 54 (87%) of these loci we assigned one or more candidate target genes (**Figure 3a, Supplementary Table 1** and **Methods**). Some of these target genes were expected based on their linear proximity to the associated variants (e.g. *ADCY5, TCF7L2, PROX1, FOXA2*), but for 72% of loci we identified more distant candidate genes. Examples of unexpected distal target genes, sometimes in addition to previously nominated proximal genes, include *SOX4* (in the *CDKAL1* locus), *OPTN* (*CDC123/CAMK1D*), *TRPM5* (*MIR4686*), *PDE8B* (*ZBED3*), *SLC36A4* (*MTNR1B*), *POLR3A and RPS24* (*ZMIZ1*), and *PHF21A* (*CRY2*) (Figure 3a,b, **Supplementary Table 1,** see http://isletregulome.org/regulomebeta/ or www.chicp.org).

**Figure 3.**
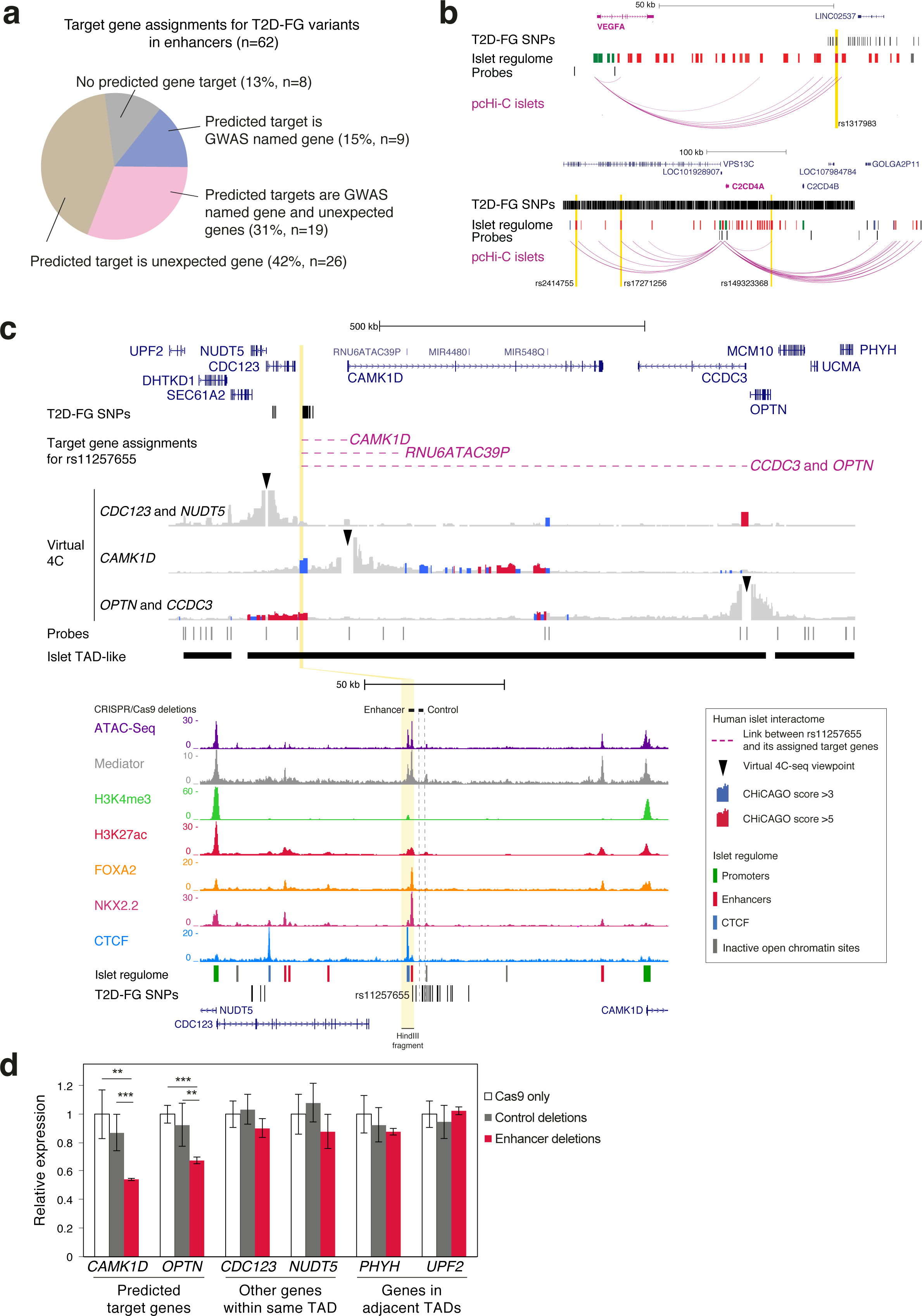
Identification of gene targets of T2D-relevant enhancers. (a) Identification of candidate target genes for 87% of 62 T2D-FG associated loci that contained genetic variants in islet enhancers (related to Supplementary Table 1). Targets were assigned through high-confidence interactions or imputations. (b) Examples of assignments of T2D-FG SNPs to candidate genes based on pcHi-C interactions. Vertical yellow stripes highlight associated variants located in enhancers and assigned to VEGFA (top) or C2CD4A (bottom) through high-confidence interactions (pink arcs). (c) Islet pcHi-C analysis defines gene targets of T2D-associated variants at CDC123/CAMK1D. Virtual 4C representations of islet pcHi-C interactions are shown for the three indicated viewpoints. Only one variant in the T2D risk haplotype (rs11257655) maps to an islet enhancer (zoomed inset). Dashed pink lines depict links between this enhancer and assigned target genes. Deleted regions are highlighted in yellow. (d) CAMK1D and OPTN are functionally dependent on the rs11257655-containing enhancer. RNA analysis of EndoC-βH3 cells was carried out after deletion of the rs11257655-containing enhancer, an adjacent control region with a T2D-associated variant but apparent regulatory function (rs33932777), or a control region in an unrelated locus (see **Supplementary Table 9**). Data from all control deletions was merged. Deletions were tested with two guide RNA pairs. Delivery of the empty vector (Cas9 only) was used as an additional reference. Data are presented as mean ± s.d. Two independent experiments were performed in triplicates. Data shown has been normalized by RPLP0 and is shown relative to the mean levels of the Cas9 controls. ** P <0.01, *** P <0.001, two-tailed Student’s t-test.

We used genome editing to validate target genes of diabetes-relevant enhancers. We performed these experiments in EndoC-βH3 cells, a glucose-responsive human β cell line^27^ that recapitulates an enhancer profile that closely resembles that of human islets (unpublished).

We first examined variants located between *CDC123 and CAMK1D*. Amongst the credible set of T2D-associated variants from fine-mapping studies, only one SNP is located in an islet enhancer, and it was found to show allele-specific enhancer activity in EndoC-βH3 cells (Figure 3c, Supplementary Figure 3b,c). This enhancer showed high-confidence pcHi-C interactions with an unexpected distant gene, *OPTN*, and moderately confident interactions (CHiCAGO = 4.42) with *CAMK1D* (Figure 3c, Supplementary Figure 3b). Accordingly, deletion or epigenetic silencing of this enhancer (but not of an adjacent region) led to selectively decreased β cell expression of both *OPTN* and *CAMK1D* (Figure 3d, Supplementary Figure 3d,e), whereas targeted activation of the enhancer stimulated their expression (Supplementary Figure 3f). These results confirm functional relationships predicted by pcHi-C maps, and point to *OPTN* and *CAMK1D* as candidate mediators of this T2D-associated genetic signal.

We then focused on rs7903146, a plausible causal SNP in the *TCF7L2* locus. This is the strongest known genetic signal for T2D, that is also known to influence islet-cell traits in non-diabetic individuals^5,28-30^. rs7903146 lies in a class I enhancer with unusually high Mediator occupancy (Supplementary Figure 3g). The SNP causes allele-specific accessibility and episomal enhancer activity in islet cells^13^, although it is unknown if this enhancer regulates *TCF7L2* because deletion of the enhancer in human colon cancer cells affects *ACSL5* rather than *TCF7L2*^31^. We found that the rs7903146-bearing enhancer has imputed and moderate confidence interactions with *TCF7L2*, but not with other genes (Supplementary Figure 3g). Consistently, deletion and targeted activation of the enhancer caused selective changes in *TCF7L2* mRNA (Supplementary Figure 3h,i). Therefore, the enhancer that harbors rs7903146 regulates *TCF7L2* in human β cells. Regardless of the possible metabolic role of this locus in other cell types^32^, this finding indicates that *TCF7L2* is a likely mediator of the genetic association between rs7903146 and islet-related traits.

Taken together, these examples illustrate the utility of human pcHi-C maps to connect regulatory elements that harbor T2D-associated variants with functional target genes that can mediate disease susceptibility mechanisms.

### Islet-specific transcription is linked to enhancer hubs

Earlier studies demonstrated that risk variants for common diseases such as T2D are highly enriched in enhancer clusters that regulate key cell identity genes (enhancer clusters, stretch enhancers, or super-enhancers) ^7,8,12,13^. We hypothesized that such domains could be more accurately defined by considering how enhancers cluster in 3D space. To identify 3D domains that regulate islet-cell identity genes, we considered multiple features of PATs (e.g. number and tissue-specificity of pcHi-C interactions, number of enhancer-based interactions, distance to TAD borders), and used multiple logistic regression analysis to assess which of these are most informative to predict islet-specific gene expression (see Supplementary Figure 4a and **Methods** for further details). This showed that the presence of ≥3 class I enhancers (enhancers with highest H3K27ac and Mediator occupancy, Figure 1c) in a PAT was a strong predictor (Supplementary Figure 4a). We thus identified all 2,633 PATs with ≥3 class I enhancers (*enhancer-rich* PATs) (Supplementary Figure 4b). Many active enhancers (∼40%) had high-confidence interactions with ≥1 promoter (Supplementary Figure 4c), and we thus merged enhancer-rich PATs with other PATs that were connected through common enhancer-mediated high-confidence interactions, yielding 1,318 islet *enhancer hubs* (Figure 4a, Supplementary Figure 4d and **Extended Data 1.4**). Enhancer hubs, therefore, are 3D chromatin domains that contain one or more interconnected enhancer-rich PATs. They contain a median of 18 enhancers and two active promoters connected through two shared enhancer interactions (Supplementary Figure 4e). They are often tissue-specific interaction domains, because hub promoters had 2.8-fold more islet-selective interactions than non-hub promoters (Wilcoxon’s P = 2.8 × 10^−36^) (**Supplementary Figure 4f,** examples in Figure 1b, 5a, Supplementary Figures 1j,k and 5a). Importantly, the genes that form part of enhancer hubs are enriched in islet-selective transcripts (Figure 4b) and in functional annotations that are central to islet cell identity, differentiation, and diabetes (Figure 4c, **Supplementary Table 2**).

**Figure 4.**
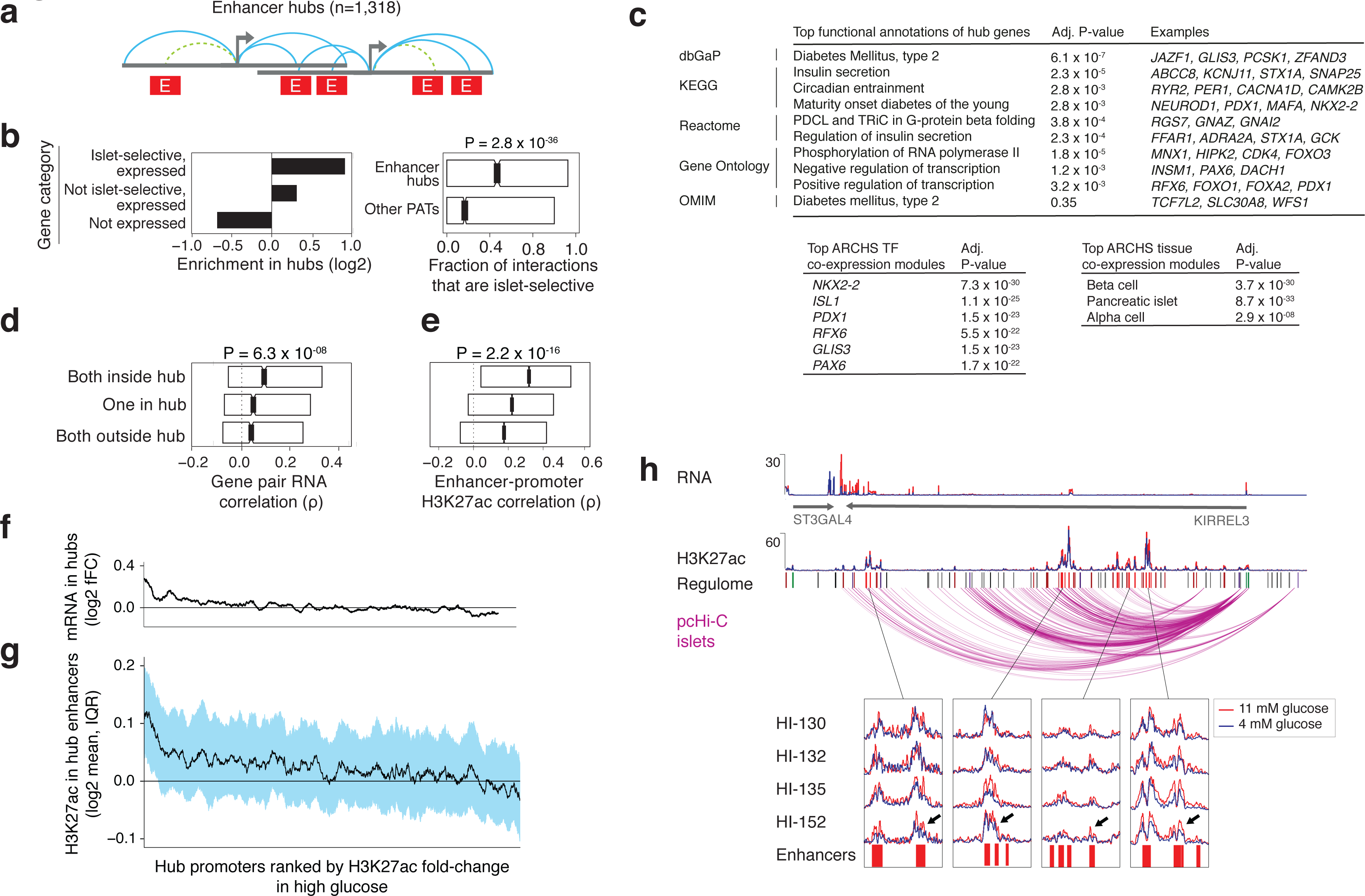
Tissue-specific enhancer hubs regulate key islet genes. (a) Schematic of enhancer hubs. Hubs are composed of one or more enhancer-rich PATs (≥ 3 class I enhancers) interconnected in cis through at least 1 common interacting enhancer. Turquoise and dashed green lines depict high-confidence and imputed assignments, respectively. Descriptive features of hubs are summarized in Supplementary Figure 4e. (b) Islet hubs are enriched in islet-selective expressed genes. Ratios were calculated relative to all annotated genes. (c) Genes connected with enhancer hubs are enriched in annotations important for islet differentiation, function and diabetes. See complete lists in **Supplementary Table 2**. (d) Gene pairs from the same hub show higher RNA correlations across human islet samples and 15 tissues than pairs from the same TAD in which only one gene or neither gene is in a hub. P values were derived with a Kruskall-Wallis analysis of variance. (e) Enhancer-promoter pairs from the same hub show high H3K27ac correlations across 14 human islet samples and 51 Epigenome RoadMap tissues, compared with pairs from the same TAD in which only one element or neither are in a hub. P values were derived with a Kruskall-Wallis test. (f-g) Culture of islets at 4 vs. 11 mM glucose shows concerted changes in H3K27ac in hub enhancers connected with glucose-dependent genes. Hub promoters were ranked by their median fold-change in H3K27ac at high glucose, so that glucose-induced promoters are on the left of the × axis. (g) shows the median glucose-dependent fold-change of H3K27ac in enhancers from hubs connecting with each promoter, and the hub IQR is shown in blue. (f) shows the median mRNA values for genes associated with each promoter. In both graphs median fold changes are shown as a running average (window = 50). (h) Coordinated glucose-induced H3K27ac in enhancers of a hub connected to the *KIRREL3* gene. The top tracks show RNA and H2K27ac binding in a representative sample. The track named regulome is color coded as in Figure 1b, where class I enhancers are bright red and promoters are green. The bottom inset shows representative regions across the hub showing coordinated glucose-induced changes in most enhancers in four human islet samples. Black arrows are used to highlight some enhancers in which the normalized H2K27ac signal is higher at 11 mM glucose (red) vs. 4 mM (blue). See also Supplementary Figure 1, **Supplementary Table 1, Data S1.4**

Consistent with the high internal connectivity of hubs, gene pairs from the same hub showed increased RNA expression correlation values across tissues and islet samples, as compared to control gene pairs from the same TAD (P = 6.3 × 10^−8^) (Figure 4d). Moreover, hub enhancers and their target promoters showed higher H3K27ac correlations than when they were paired with non-hub promoters (P = 2.2 × 10^−16^) (Figure 4e). These findings are consistent with enhancer hubs as functional regulatory domains.

We next tested the functional connectivity between hub enhancers and promoters in a hub that contains *GLIS3,* a gene that is mutated in neonatal diabetes but also harbors T1D and T2D susceptibility variants ^33-35^. We identified an intronic class I enhancer containing four T1D-T2D risk variants that alter episomal enhancer activity (Supplementary Figure 4g,h). This enhancer showed indirect interactions with a nearby islet-specific *GLIS3* promoter through common hub enhancers, and direct interactions with *RFX3,* another islet transcription factor gene ^36^ (Supplementary Figure 4g). In line with such interconnectivity, deletions of the enhancer led to decreased mRNA levels of both hub genes, *GLIS3* and *RFX3* (Supplementary Figure 4i). These studies, therefore, identify functional target genes of an enhancer that harbors diabetes-associated variants, and points to functional connectivity of two enhancer hub promoters.

To further explore the behavior of hubs as functional domains, we again examined islets exposed to moderately low vs. high glucose concentrations, using above-mentioned datasets. Glucose-induced enhancers and mRNAs were highly enriched in hubs, compared with non-hub genomic regions (Fisher’s P = 1.1 × 10^−7^ and 2.2 × 10^−16^, respectively). Of 297 promoters that showed increased H3K27ac at high glucose, 94 were contained in hubs, and 65% of these also showed a significant glucose-dependent mRNA increase (**Supplementary Table 3**). If hubs are indeed regulatory domains, we expect that hub enhancers connected to glucose-induced genes should show concordant glucose-dependent changes. We found that hubs of glucose-induced promoters showed a widespread parallel increase in H3K27ac levels of their enhancers (**Figure 4f-h**). Thus, varying glucose concentrations can elicit chromatin changes in human islets at the level of broad regulatory domains. Taken together, our findings indicate that enhancer hubs are often regulated by varying glucose concentrations, and they have properties of functional regulatory units.

### Enhancer hubs contain super-enhancers and enhancer clusters

We compared islet enhancer hubs with previously defined islet enhancer domains, such as linear enhancer clusters and super-enhancers (Supplementary Figure 4j). We found that hubs overlapped 70% of enhancer clusters ^8^, and 87% of super-enhancers defined with a standard algorithm ^12^ (Supplementary Figure 4k-m). Hubs, however, differ in that they are connected with their target genes (**Extended Data 1.4**). Furthermore, hubs capture spatial clusters of Mediator-bound enhancers that do not cluster in the linear genome and do not fulfill definitions of super-enhancers and enhancer clusters (Supplementary Figure 4n-p) ^8,12^. In fact, many hubs contained several inter-connected enhancer clusters or super-enhancers (Supplementary Figure 4q-s). This is illustrated by the *ISL1* locus, which has several enhancer clusters and super-enhancers distributed across an entire TAD, whereas pcHi-C points to a single hub that connects dozens of enhancers with *ISL1* and lncRNA *HI-LNC57* (Figure 5a). Thus, enhancer hubs are 3D domains that often include one or more enhancer clusters or super-enhancers and their target gene(s).

**Figure 5.**
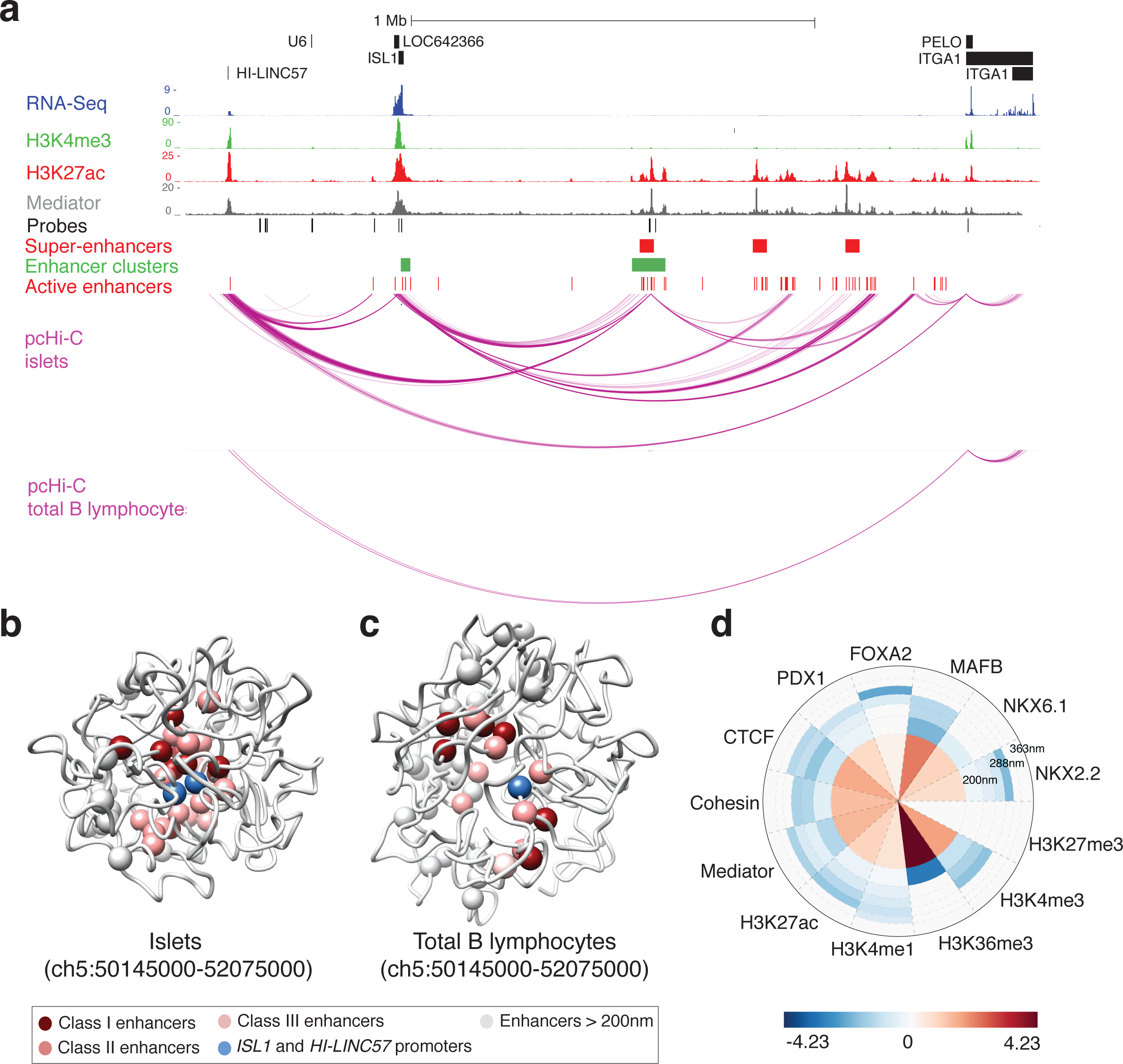
Tissue-specific topology of the *ISL1* enhancer hub. (a) The *ISL1* locus forms a tissue-specific enhancer hub. Human islet epigenome maps and high-confidence pcHi-C interactions in islets and total B lymphocytes are shown to illustrate that multiple active enhancers, super-enhancers and enhancer clusters distributed across a TAD share islet-selective 3D interactions with common genomic regions including *ISL1* and *HI-LNC57*. (b-c) 3D chromatin conformation models of the ISL1 enhancer hub created with all interaction fragments from this locus in pcHi-C libraries of human islets (b) and total B lymphocytes (c). Images represent the top scoring model from the ensemble of structures that best satisfies the spatial restraints, topological constraints, confinement, and supercoiling of the chromatin fibre. Islet enhancers and promoters are represented as white spheres. Class I, II and III enhancers within 200nm of *ISL1* promoter (in blue) are colored in dark to light red. Note that the intergenic lncRNA HI-LNC57 (represented in blue) is in close proximity with *ISL1* promoter. These models show that active islet regulatory elements interact in a common restricted space in islet nuclei. Additional views of models are shown in Supplementary Figure 5b,c and **Supplementary Videos 1 and 2**. (d) Density wheel depicting histone modification mark and transcription factor ChIP-seq signal in human islets mapped on the 3D model of *ISL1* enhancer hub in human islets. This radial representation is centered on *ISL1* promoter.

### Tissue-specific architecture of the *ISL1* enhancer hub

To gain further insight into the topology of enhancer hubs and their relationship with TADs, we simulated 3D models of the *ISL1* locus. We considered islet pcHi-C interaction frequencies as a proxy for spatial proximities between loci, built interaction matrices at a resolution of 5 kb, and converted the frequency of interactions between genomic segments into spatial restraints ^37,38^. We then used molecular dynamic optimization to generate an ensemble of 500 models that best satisfied the imposed restraints. This revealed a topology in which islet enhancers and target genes co-localize in a constrained space (Figure 5b,c, Supplementary Figure 5b-g, **Supplementary Videos 1-2**). By contrast, models built from B lymphocyte pcHi-C libraries showed decreased aggregation of these regions (Figure 5b-d, Supplementary Figure 5b-g). Accordingly, active chromatin features and architectural proteins aggregated near *ISL1* in islets but not in total B lymphocytes (Figure 5d, Supplementary Figure 5d). These models, which capture the average topology in a population of cells, serve to highlight that whereas TADs are defined as genomic intervals in unidimensional maps, hubs represent a 3D subspace of TADs that harbors interactions between active genomic regions distributed across the linear TAD space. Taken together, these results leverage knowledge of 3D chromatin structure to define islet-specific enhancer domains, many of which can be directly linked to key genes underlying cell identity and diabetes.

### Prominent contribution of islet hubs to the heritability of T2D and islet traits

Previous evidence that T2D and FG susceptibility variants are enriched in islet enhancer clusters ^7,8,13,30,39^ prompted us to examine the relationship between diabetes-associated variants and our newly defined enhancer hubs. We found that SNPs from the 109 T2D-FG associated loci (**Supplementary Table 4**) were enriched in islet pcHi-C high-confidence interaction regions (Figure 6a). Furthermore, they were selectively enriched in class I enhancers that form part of islet enhancer hubs, rather than in those located outside hubs, or in other active enhancer subtypes (Figure 6b, Supplementary Figure 6a-f). These observations indicated that enhancer hub class I enhancers, rather than other enhancers, define a critically important genomic space for T2D genetic susceptibility.

This finding led us to estimate the contribution of sequence variation across enhancer hubs to the heritability of T2D. Several GWA studies suggest that a major portion of the heritability of common diseases is driven by a large number of variants that individually have not achieved genome-wide significance, yet exert a large aggregate effect ^34,40-42^. Consistent with this notion, common variants that have so far not shown genome-wide significance for T2D association, but are located in pcHi-C interacting regions or hub class I enhancers, showed lower association p-values than expected distributions (Figure 6c,d).

**Figure 6:**
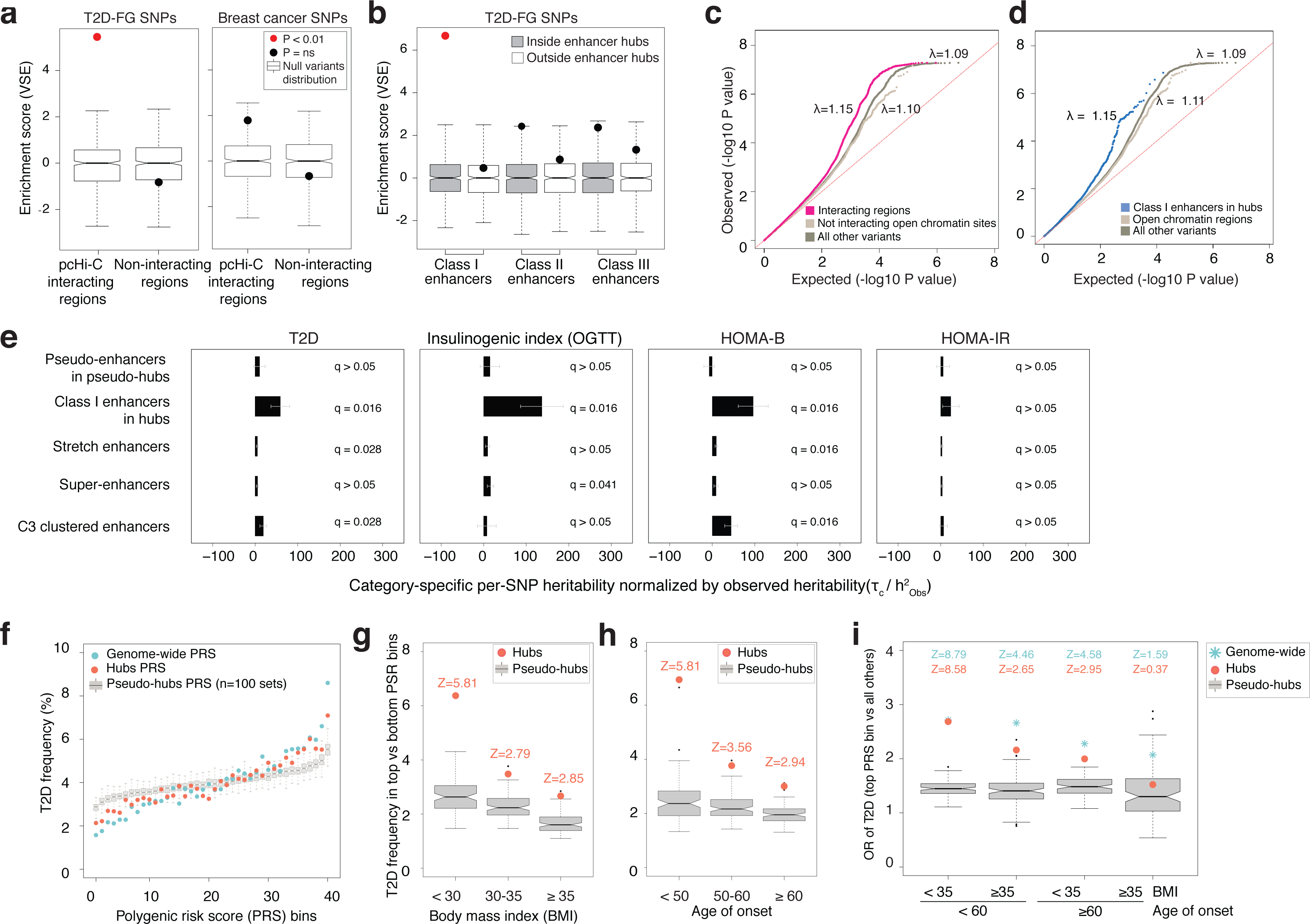
Islet enhancer hubs carry an increased burden of T2D-associated variants. (a) Variant Set Enrichment (VSE) of T2D and FG-associated variants (dot in left graph, see **Supplementary Table 3**) and breast cancer-associated variants (dot in right graph) in islet high-confidence pcHi-C interacting fragments. Box plots show null distributions based on 500 permutations of matched random haplotype blocks. Red dots indicate significant enrichment relative to the null distribution (Bonferroni–corrected P < 0.01). (b) T2D and FG GWAS-significant variants are specifically enriched in class I islet enhancers within enhancer hubs, but not in other islet enhancer subtypes (see also Supplementary Figure 6c). (c) Genomic inflation of T2D association P-values for non-GWAS significant (p > 5×10−8) variants located in islet interacting regions. Quantile-quantile (Q-Q) plots are shown for variants in pcHi-C interacting regions, as described in panel A (magenta), non-interacting open chromatin regions (light beige), and all other variants reported in the summary statistics (brown). The λ coefficients correspond to the median χ^2^ test statistic divided by the median expected χ^2^ test statistic under the null hypothesis. (d) Genomic inflation of T2D association P-values for non-GWAS significant variants in class I islet enhancers from enhancer hubs (blue), compared to variants in open chromatin regions outside hubs (light beige) and to all other variants (brown). (e) Heritability estimates of variants in five islet enhancer domain subtypes for T2D, insulinogenic index (OGTT), homeostasis model assessment of ß-cell function (HOMA-B) and insulin resistance (HOMA-IR). Bars show category specific per-SNP heritability coefficients (τc), which estimate the expected increase in phenotypic variance accounted by a single SNP from a given functional category, divided by the LD score heritability (h^2^) values observed for each trait. We estimated τc coefficients by performing independent stratified LD score regression, controlling for 53 functional annotation categories included in the baseline model. All normalized τc coefficients were multiplied by 107 and shown with a SEM. (f) Impact of islet hub polygenic risk scores (PRS) on T2D frequency. T2D frequency (y-axis) was calculated in 40 bins, each one representing 2.5% of individuals in the UK Biobank validation dataset. PRS values were calculated with the entire set of common genetic variants (light blue dots), the fraction of islet non-hub variants (dark blue dots) and islet hub variants (orange dots), or variants from 100 sets of pseudo-hubs (boxplots with IQR). (g) Polygenic risk stratified by BMI. The T2D risk ratio was calculated as the frequency of T2D cases in the highest 2.5% PRS bin divided by the frequency of T2D cases in the lowest 2.5% risk bin. Boxplots show the risk ratio for PRS from 100 sets of pseudo-hubs, the dark blue dot corresponds to the islet non-hub PRS and the orange dot shows that of islet hub PRS. A Z-score of the risk ratio with islet hub and non-hub PRS, respectively, was computed to define the number of standard deviations above the average risk ratio of the pseudo-hub distribution. (h) Polygenic risk stratified by age of onset of T2D. Boxplots show the T2D risk ratio in highest vs. lowest risk bins for PRS from 100 sets of pseudo-hubs and the dark blue and orange dot shows that of islet non-hub and hub PRS, respectively. T2D cases were stratified using age of onset information and age at recruitment was used in control samples. (i) Polygenic risk stratified by BMI and age of onset of T2D. Odds ratio (OR) for T2D were calculated by comparing 2.5% individuals with the highest PRS vs. all the other individuals via adjusted logistic regression. Boxplots show OR for 100 sets of pseudo-hubs, orange dots are OR from islet hub PRS, and the dark and light blue stars shows the OR for the islet non-hub and the genome-wide PRS models. Controls were stratified by age at recruitment. Statistical significance for islet hub OR was calculated using a Z-score defined as the number of standards deviations above the average OR of the pseudo-hub distribution. See also Supplementary Figure 6, **Supplementary Table 4** and **Data S1**.

These observations prompted us to quantify the overall contribution of common variants in islet hubs to the heritability of T2D. We used stratified LD score regression ^43^, and found that hub enhancers (as well as other islet enhancer definitions) had a significantly enriched per-SNP T2D heritability coefficient (*q* = 1.64 × 10^−2^) (Figure 6e, Supplementary Figure 7, **Supplementary Table 5**). Although islet dysfunction is central to the pathophysiology of T2D, other tissues (liver, adipose, muscle, gut, CNS, among others) are also critically important. We predicted that common genetic variation in islet hub enhancers should have an even stronger impact on the heritability of pancreatic islet function. Indeed, common variation in hub class I enhancers, which represent 0.26% of genomic SNPs, explained 9.9% of observed genetic heritability for T2D, 21.9% for acute insulin secretory response in intravenous glucose tolerance tests (AIR-IVGTT)^29^, 17.2% for HOMA-B models of β-cell function, and 31.2% for an insulinogenic index based on oral glucose tolerance tests (OGTT) ^44^ (**Supplementary Table 5**). Accordingly, islet hub variants showed even greater heritability enrichment estimates for islet-cell traits than for T2D (**Figure 6e, Supplementary Figure 7a-f, Supplementary Table 5**). In sharp contrast, islet hub variants showed no enrichment for HOMA-IR, an estimate of insulin resistance (Supplementary Figure 7e). Importantly, although enhancer clusters, stretch enhancers, or super-enhancers also showed significant enrichments of heritability estimates for islet-cell traits and T2D, these estimates were consistently larger for hub enhancers (**Figure 6e**, **Supplementary Figure 7a-d**). These results indicate that enhancer hubs are genomic spaces that play a prominent role in the heritability of islet function and T2D.

### Common variants in islet hubs provide tissue-specific risk scores

The improved definitions of genomic regions relevant to islet function prompted us to examine if islet hub variants could be harnessed to identify individuals in whom variation in islet function, rather than other T2D-relevant cellular functions, plays a preponderant role in T2D susceptibility. Recent studies suggest that polygenic risk scores (PRS) that integrate the combined effects of very large number of variants, including many that lack genome-wide significant association, can be used to identify individuals at risk for polygenic diseases ^34,40-42,45^. The predictive power of PRS for T2D is likely to increase as larger GWAS datasets become available, although it is already possible to explore the potential of hub variants to define tissue-specific specific risk profiles.

We first calculated a PRS using all common variants from a recent BMI-adjusted T2D GWAS meta-analysis ^46^, and examined the ability of this genome-wide PRS to predict T2D in the UK Biobank population cohort ^47,48^. The UK Biobank cohort was divided into a testing dataset used to train the PRS model, and a validation dataset of 236,236 individuals. This showed that individuals of the validation dataset with the 2.5% highest PRS had a 7.1-fold higher frequency of T2D than those in the lowest 2.5%, and an incident T2D hazard ratio of 2.5 relative to the entire population (**Figure 6f, Supplementary Figure 7h**). Next, we built PRS models with islet hub enhancer and promoter variants. Although hub regulatory regions represent < 2% of genomic space, 2.5% of individuals with highest hub PRS scores had 4-fold higher frequency of T2D than those in the lowest bin and an incident T2D hazard ratio of 2.1 relative to the entire population (**Figure 6f, Supplementary Figure 7h**). For comparison, we also built PRS from pseudo-enhancer hubs of similar size that were redistributed across all TADs 100 times. PRS from pseudo-hubs only average 2.2-fold T2D difference between extreme PRS bins, and the highest PRS bin showed an average incident T2D hazard ratio of 1.4 (**Figure 6f)**.

We then tested whether enhancer hub variants define risk profiles that are qualitatively different from other genomic regions. Monogenic defects in islet transcription factors typically cause early-onset diabetes in the absence of obesity, suggesting that islet cis-regulatory variants could also have a more prominent impact on T2D risk at an earlier age and at lower BMI. Individuals with extreme hub PRS showed greatest deviations in risk ratios from pseudo-hubs, and tended to show more extreme T2D incident HR values, in individuals with BMI <30, and earlier age of onset of T2D (**Supplementary Figure 7i-j**). We thus examined the risk of T2D stratified by both variables (BMI and age of onset), and calculated the odds ratios for T2D in individuals with the highest PRS values (Figure 6g). This showed an odds ratio of 2.7 for T2D with BMI < 35 and onset < 60 years in individuals with the highest hub PRS (2.5% of individuals) vs. all others (97.5% of individuals). This odds ratio was a major deviation from that observed with pseudo-hub PRS (Z = 8.5). At the other extreme of the phenotypic spectrum (T2D with BMI ≥ 35 and age of onset ≥ 60), individuals with the highest islet hub PRS showed a lower odds ratio (OR = 1.5), which did not differ from that of pseudo-hub genomic regions. Thus, polygenic scores built with islet hub variants provide qualitative risk profile differences relative to other genomic regions. They indicate that risk scores built from a small fraction of the genome that regulates islet cell transcription and impacts the heritability of islet function retain a substantial ability to predict T2D diabetes, and have the potential to assist the stratification of individuals in whom islet regulatory variation plays a role in pathogenesis.

## Discussion

We have created 3D genome annotations that link human pancreatic islet enhancers to gene promoters. This revealed enhancer hubs that exhibit features of regulatory units, and carry a significant burden of genetic variants that influence islet-cell function and T2D risk. The data provides a resource for the prioritization of putative causal genes in T2D pathophysiology, and for the development of tissue-specific genetic risk scores for patient stratification.

Individual genes are often regulated by the concerted action of multiple long-range enhancers. Enhancer domains have accordingly been defined based on how enhancers cluster in the linear genome ^7,8,12,13^, through in-depth functional characterization of specific loci ^49-52^, or based on evolutionary conserved noncoding sequence blocks ^53,54^. We have now grouped enhancers and promoters based on their spatial proximity within islet-cell nuclei. Our study provides unbiased maps of hundreds of pancreatic islet enhancer hubs, many of which contain one or more previously reported linear enhancer clusters that are now seen to form part of larger 3D domains. Hub enhancers can sometimes be distributed across the entire linear space of TADs, yet differ significantly in that they only occupy a restricted portion of the 3D TAD space, and thereby only target a subset of annotated TAD genes. Hub enhancers exhibit concerted histone acetylation changes across tissues and dynamic settings, consistent with coherent regulatory units. Importantly, islet hubs interact with genes that encode for key islet-cell functions, and are thus critically relevant to human diabetes.

The spatial enhancer clusters described here are compatible with earlier observations derived from lower resolution Hi-C maps, which showed broad genomic regions that exhibit unusually high interaction frequencies ^55^. Our findings also align with numerous individual examples of well-characterized enhancer hubs ^50,52^. We have now systematically annotated hundreds of analogous domains in human islets, and define their candidate target genes.

The 3D enhancer domains described here have implications for our understanding of the genetic basis of T2D. Recent studies suggest that the heritability of common diseases is best explained by the aggregate effect of a large number of SNPs that impact on disease-relevant cellular networks, including many that do not reach conventional significance thresholds for association ^34,40,41^. Our 3D chromatin-based annotations define genomic spaces that play a hierarchically prominent role in the heritability of insulin secretory function and T2D. We propose that enhancer hubs provide a useful gene-centric framework to dissect the contribution regulatory variants to T2D, but also for efforts to define causal non-coding mutations in monogenic diabetes.

Moreover, regulatory spaces defined here can contribute to characterize the heterogeneous, multiorganic nature of T2D pathophysiology ^56,57^. Polygenic risk scores hold promise to exploit GWAS findings for the prediction of common diseases ^58^, perhaps combined with non-genetic risk factors. Theoretically, they can also qualify the genetic risk based on specific underlying mechanisms that are likely to vary in different patients with T2D. Our findings illustrate how regulatory domains can be exploited to define polygenic risk estimates that affect diabetes susceptibility through their influence on tissue-specific gene regulation. Further developments of this approach could assist future patient-specific preventive and therapeutic recommendations. Beyond this prospect, pancreatic islet 3D genome annotations reported here provide a resource for functional interpretation of noncoding variants that underlie the molecular mechanisms of T2D susceptibility.

## Acknowledgements

This research was supported by the National Institute for Health Research (NIHR) Imperial Biomedical Research Centre. Work was funded by grants from the Wellcome Trust (WT101033 to J.F. and WT205915 to I.P.), Horizon 2020 (Research and Innovation Programme 667191 to J.F., 633595 to I.P. and 676556 to M.A.M-R; Marie Sklodowska-Curie 658145 to I.M-E., and 43062/ZENCODE to G.A.), European Research Council (609989 to M.A.M-R.). Marató TV3 (201611, to J.F., M.A.M-R.), Ministerio de Economía y Competitividad (SAF2018 to J.F., BFU2013-47736-P to M.A.M-R., IJCI-2015-23352 to I.F.), AGAUR (to M.A.M-R.). UK Medical Research Council (MR/L007150/1 to P.F.), World Cancer Research Fund (WCRF UK to I.P.) and World Cancer Research Fund International (2017/1641), Biobanking and Biomolecular Resources Research Infrastructure (BBMRI-NL, NWO 184.021.007 to I.O.F.). Work in IDIBAPS, CRG and CNAG was supported by the CERCA Programme, Generalitat de Catalunya and Centros de Excelencia Severo Ochoa 2013-2017 (SEV-2012-0208). Human islets were provided through the European islet distribution program for basic research supported by JDRF (3-RSC-2016-160-I-X). We thank Natalia Ruiz-Gomez for technical assistance, Rodrigo Liberal Fernandes, Thomas Thorne (University of Reading), and Alvaro Perdones-Montero (Imperial College London) for helpful discussions regarding Machine Learning approaches, the CRG Genomics Unit, and the Imperial College High Performance Computing Service.

## Author contributions

Conceptualization, I.M-E. and J.F.; Methodology, I.M-E., S.B-G., I.C., J.P-C., J.M-E., D.M.Y.R., B.M.J., G.A. and A.B.; Software: I.M-E., S.B-G., J.P-C., J.M-E., D.M.Y.R., G.A., I.F. and C.C.M.; Formal Analysis: I.M-E., S.B-G., I.C., J.P-C., J.M-E., D.M.Y.R., G.A., I.F. and C.C.M.; Investigation, I.M-E., I.C., B.M.J., J.G-H.; Data Curation, I.M-E., S.B-G., J.P-C., J.M-E., D.M.Y.R., G.A. I.F., I.M., L.P. and M.R.; Writing – Original Draft, I.M-E., I.C., S.B-G., and J.F; Writing – Review & Editing, I.M-E., I.C., S.B-G., D.M.Y.R., B.M.J., I.F., and J.F.; Funding Acquisition, M.A.M-R., P.F. and J.F.; Resources, E.V.R.A., A.L., T.H., I.P., A.P.G., M.E.J., O.P., N.G., P.R., J.M.M., D.T., L.Pi., T.B., E.J.P.d.K., J.K-C., F.P. and T.H.; Supervision, I.M-E., D.M.Y.R., I.F., L.P., M.A.M-R., P.F. and J.F.

---

**Supplementary Figure 1.**
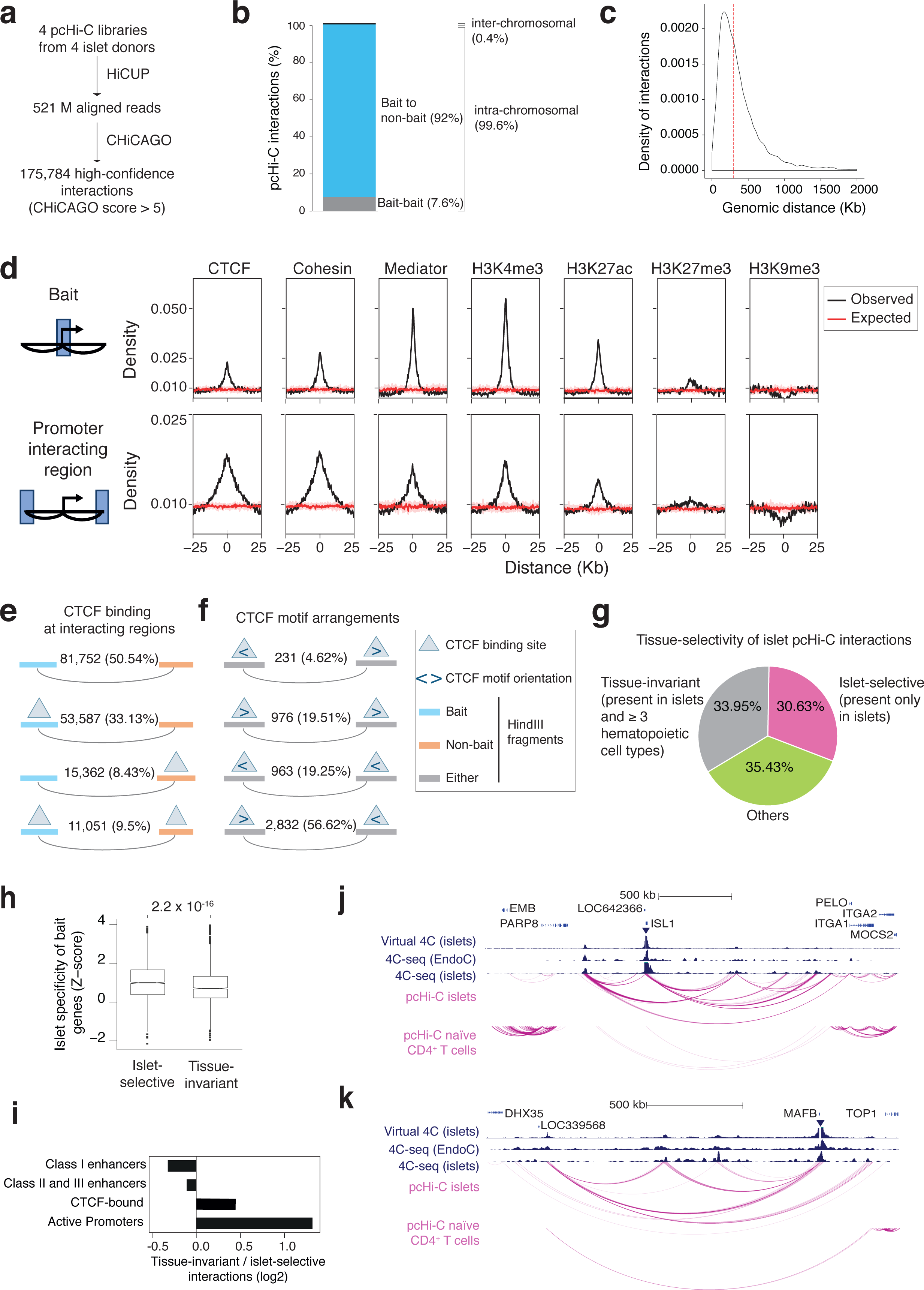
**pcHi-C and chromatin landscape of human islets.** Related to Figure 1. (a) Schematic representation of the pcHi-C analysis workflow. (b) Relative frequency of high-confidence interactions between baits and promoter-interacting regions. (c) Distribution of distances from bait to promoter-interacting region for high-confidence interactions. The dashed line represents the median distance. (d) Observed occupancy profiles for CTCF, cohesin, Mediator, H3K4me3, H3K27ac, H3K27me3, H3K9me3 (black line) in ± 25 Kb regions centered on interacting pcHi-C baits (top), and promoter-interacting regions (bottom). Expected occupancy profiles after randomizing 10 times the positions of indicated signals are represented with a red line, and interquartile ranges are shown as a shade. (e) Relative frequency of CTCF binding sites in baits and non-bait interacting regions. Nearly 50% of interactions are associated with CTCF binding sites in at least one of the interacting regions. Percentages are calculated relative to all intra-chromosomal bait-non-bait interactions. (f) CTCF-binding motif orientation at CTCF-bound interacting regions. Of all four possible CTCF motif orientation configurations, 56.62% of 9,657 interactions are in a convergent orientation, consistent with expectations. (g) Tissue-selectivity of islet pcHi-C interactions. pcHi-C datasets from four primary hematopoietic cell types (erythroblasts, macrophages, naïve CD4+ T cells and total B lymphocytes) were used to classify islet interactions as tissue-invariant, if they were also observed in at least four datasets, or islet-selective, if they were observed in islets and no other datasets. Intermediate selectivity was classified as others. (h) Islet-selectivity of interactions is concordant with islet-selective gene expression. Genes located in baits that form islet-selective interactions show increased gene expression islet-specificity scores than those from tissue-invariant interactions. The islet-specificity Z score was calculated with a gene expression distribution from 18 human tissues. P value was calculated with Wilcoxon’s signed ranked test. (i) Ratio of tissue-invariant to islet-selective interactions overlapping with major open chromatin classes, normalized by the total number of tissue-invariant and islet-selective interactions. All categories showed significant differences with interactions in the remaining genome (Fisher’s P < 0.01). (j,k) pcHi-C recapitulates interactions identified by 4C-seq in human islets and the human β cell line EndoC-βH1 at (j) ISL1 and (k) MAFB. The top track depicts a virtual 4C representation of human islet pcHi-C data in these two promoters. High-confidence interactions from human islets and naïve CD4+ T cells are shown below. The inverted triangle depicts the viewpoint used in the 4C-seq experiments and in the virtual 4C representation.

**Supplementary Figure 2.**
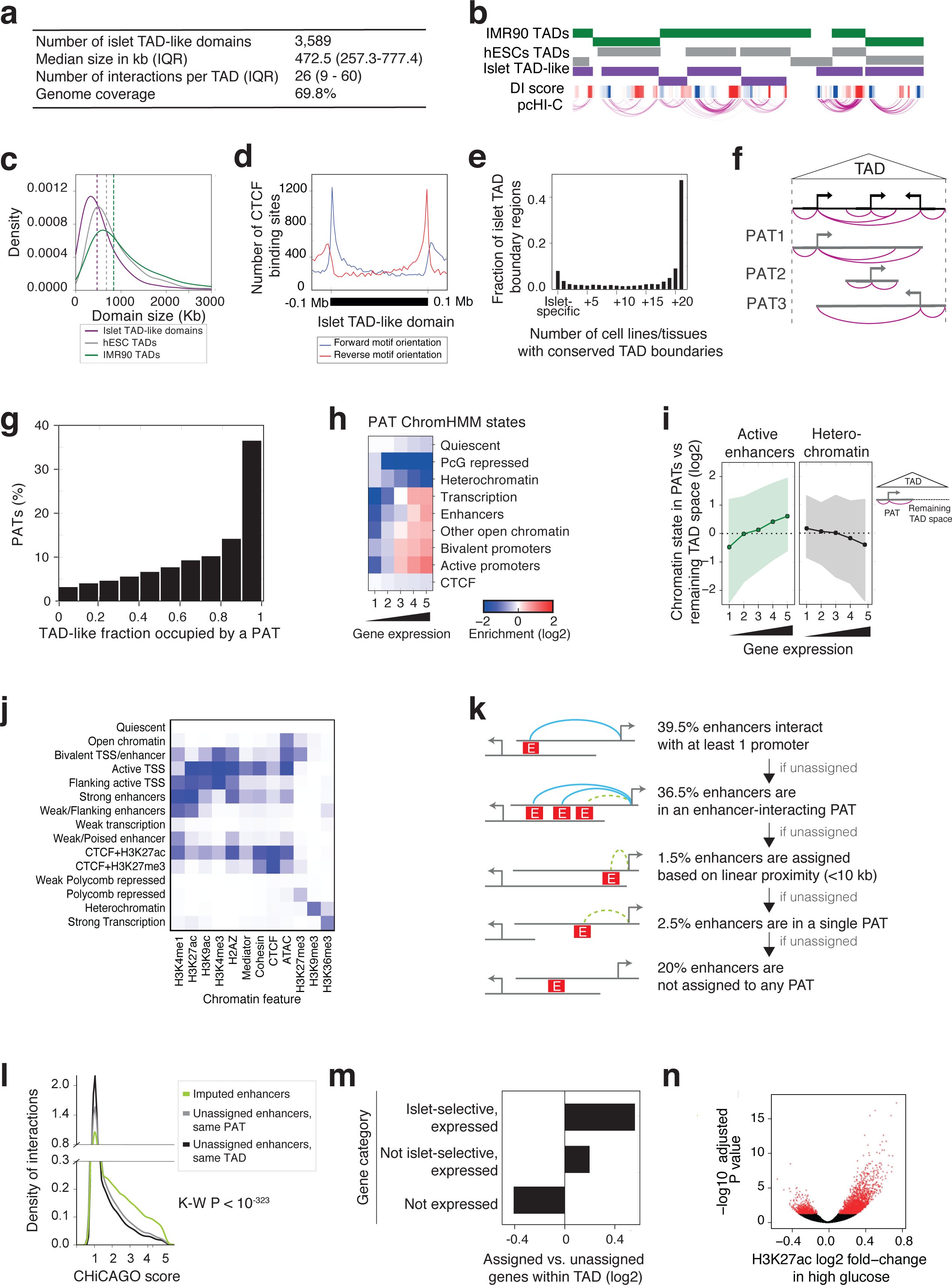
Definition of TAD-like domains, PATs, and enhancer-gene assignments. Related to Figure 2. (a) Descriptive features of islet TAD-like domains defined with directionality scores. (b) Representative example of human islet TAD-like compartments in chromosome 11:1132582-4719948. Negative and positive directionality index (DI) scores are represented in blue and red, respectively. ESC and IMR90 TADs generated with Hi-C are shown for reference. (c) Relative size of TAD-like domains in human islets and Hi-C TADs defined in ESC and IMR90 cells. (d) TAD-like domains display known features of TADs defined in other tissues, such as enrichment of CTCF binding sites in TAD borders and convergent CTCF motif orientation. (e) Tissue-selectivity of islet TAD-like boundary regions was calculated by comparison with TADs defined by Hi-C in 21 tissues. (f) Schematic of promoter-associated territories (PATs), defined as the genomic space that spans high-confidence interactions originating from one bait. (g) Fraction of islet TAD-like linear genomic space occupied by each PAT. (h) ChromHMM state enrichments in PATs were consistent with the expression level of their associated genes. The heatmap shows ChromHMM state median log2 fold-enrichments in PATs over their genomic distributions, in 5 bins based on bait gene expression levels in human islets. See also Supplementary Figure 2j. (i) Active enhancer or H3K9me3-enriched ChromHMM states in PATs are enriched over the remaining TAD-like space in accordance with the expression of PAT genes in islets. Only PATs that were at least 25% smaller than their TAD were used (n=7,085). Median enrichments (circles) and IQR (shade) are shown. (j) Heatmap showing the emission probabilities of the 15 ChromHMM states for all islet chromatin features used to create the model. (k) Diagram illustrating the sequential steps used to impute the assignment of islet enhancers to candidate target genes. Further details of this procedure are described in **Methods**. (l) Density of interactions at different CHiCAGO scores for imputed enhancer-promoter pairs vs. unimputed enhancer-promoter pairings from the same PATs or TADs (Kruskal-Wallis P<10^−323^). Genes assigned through high-confidence interactions or imputation were enriched in islet-specific genes, as compared with unassigned control genes from the same islet TAD-like structure (Chi-square P = 6 × 10^−08^). (n) Glucose causes induced H3K27 acetylation in a large number of islet enhancers. The dots represent mean log2 fold change of H3K27 acetylation in H3K27ac-enriched regions in islets exposed to 4 mM vs. 11 mM glucose. Red dots are values showing Benjamini-Hochberg adjusted P ≤ 0.05.

**Supplementary Figure 3.**
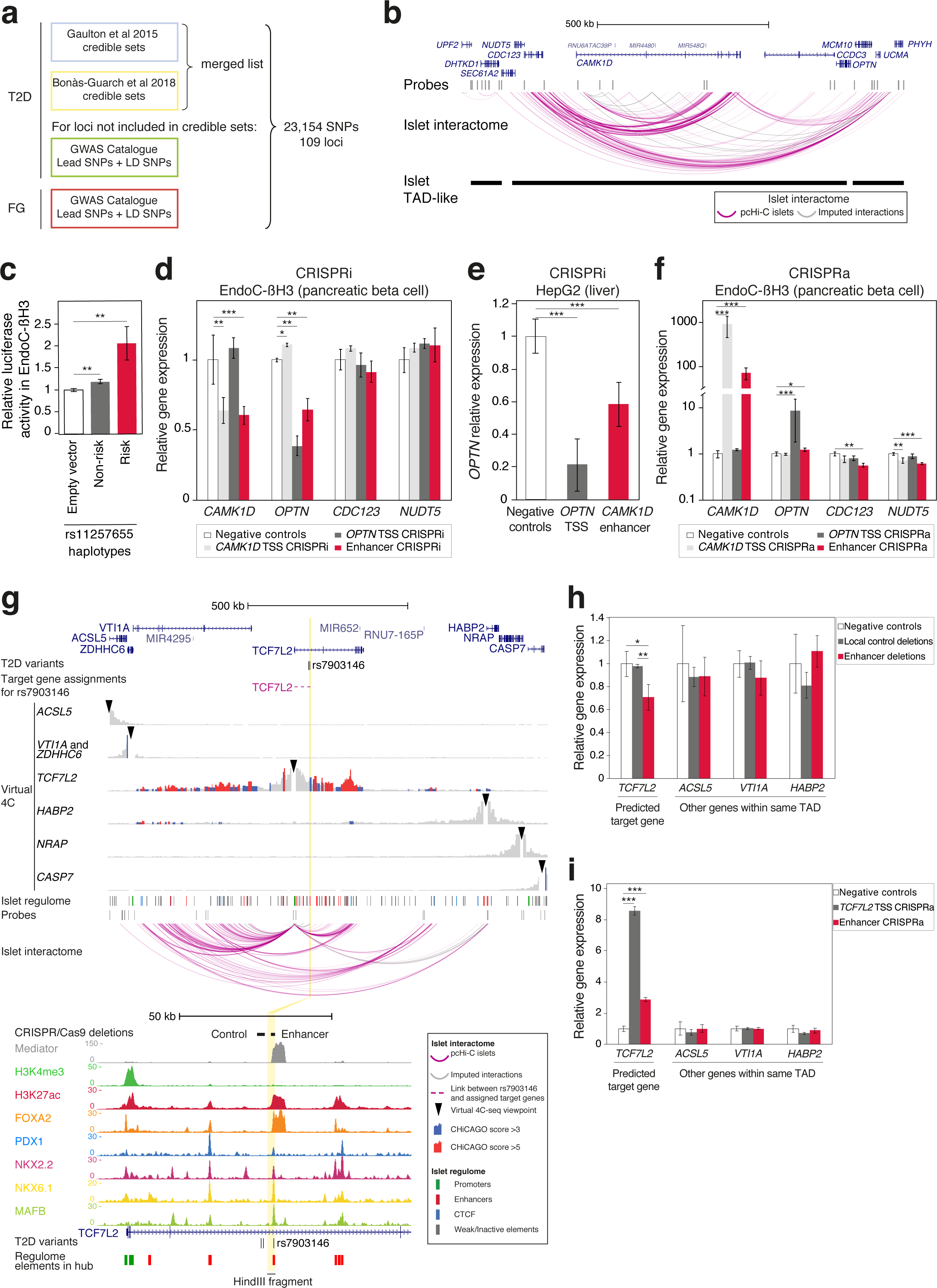
**CRISPR-Cas9 mediated perturbations in β-cells**. Related to Figure 3. (a) Schematic diagram of the selection of T2D and FG-associated variants used in the assignments of 3D targets analysis (see also **Methods**, **Extended Data 1** and **Supplementary Table 1**). (b) Map of the *CDC123/CAMK1D* locus (an alternative representation of the same region depicted in Figure 3C,D) showing pcHi-C high-confidence interactions (pink) and imputed assignments (grey). (c) Luciferase assay in the human β cell line EndoC-βH3 shows allele-dependent activity for the rs11257655-enhancer. Data are presented as mean ± s.d. Two independent experiments were performed in quadruplicates. ** P <0.01, two-tailed Student’s t-test (d,e) Analysis of *OPTN* and *CAMK1D* mRNA after CRISPR inhibition (CRISPRi) of the rs11257655-enhancer in EndoC-βH3 cells (d) and (e) HepG2 cells. Bars show average values of three guide RNAs targeting either the rs11257655 enhancer, or the OPTN transcriptional start site as control. Data are presented as mean ± s.d. Two independent experiments were performed in triplicates. Data shown was normalized by *RPLP0* and is shown relative to the mean levels of the negative controls. ** P <0.01, *** P <0.001, two-tailed Student’s t-test. (f) Analysis of *OPTN* and *CAMK1D* mRNA after CRISPR activation (CRISPRa) in EndoC-βH3 cells. Analysis as in panel D. (g) Analysis of the T2D-associated locus *TCF7L2*. Virtual 4C maps centered on all genes in this locus show that the T2D-associated regulatory variant rs7903146 connects with *TCF7L2* through moderate-confidence interactions and an imputed assignment, but shows no evidence for interactions with other genes. The HindIII fragment that contains the enhancer is highlighted in yellow. The bottom panel shows that this enhancer shows an unusually strong occupancy by Mediator, in addition to islet-enriched transcription factors. (h) Analysis of active islet genes in this locus, *TCF7L2, VT11A, HABP2, ACSL5* in EndoC-βH3 cells after deletion of either the rs7903146-enhancer or a control region in the same locus (see Supplementary Figure 3g). Deletions were tested with 2 different guide RNA pairs. Data are presented as mean ± s.d. Two independent experiments were performed in triplicates. Data was normalized by RPLP0 and is shown relative to the mean levels of the negative control deletions. *** P <0.001, two-tailed Student’s t-test. (i) Analysis of *TCF7L2, VT11A, HABP2* mRNA in EndoC-βH3 cells after CRISPR activation (CRISPRa) of this enhancer. The analysis was carried out as in panel H. *** P <0.001, two-tailed Student’s t-test.

**Supplementary Figure 4.**
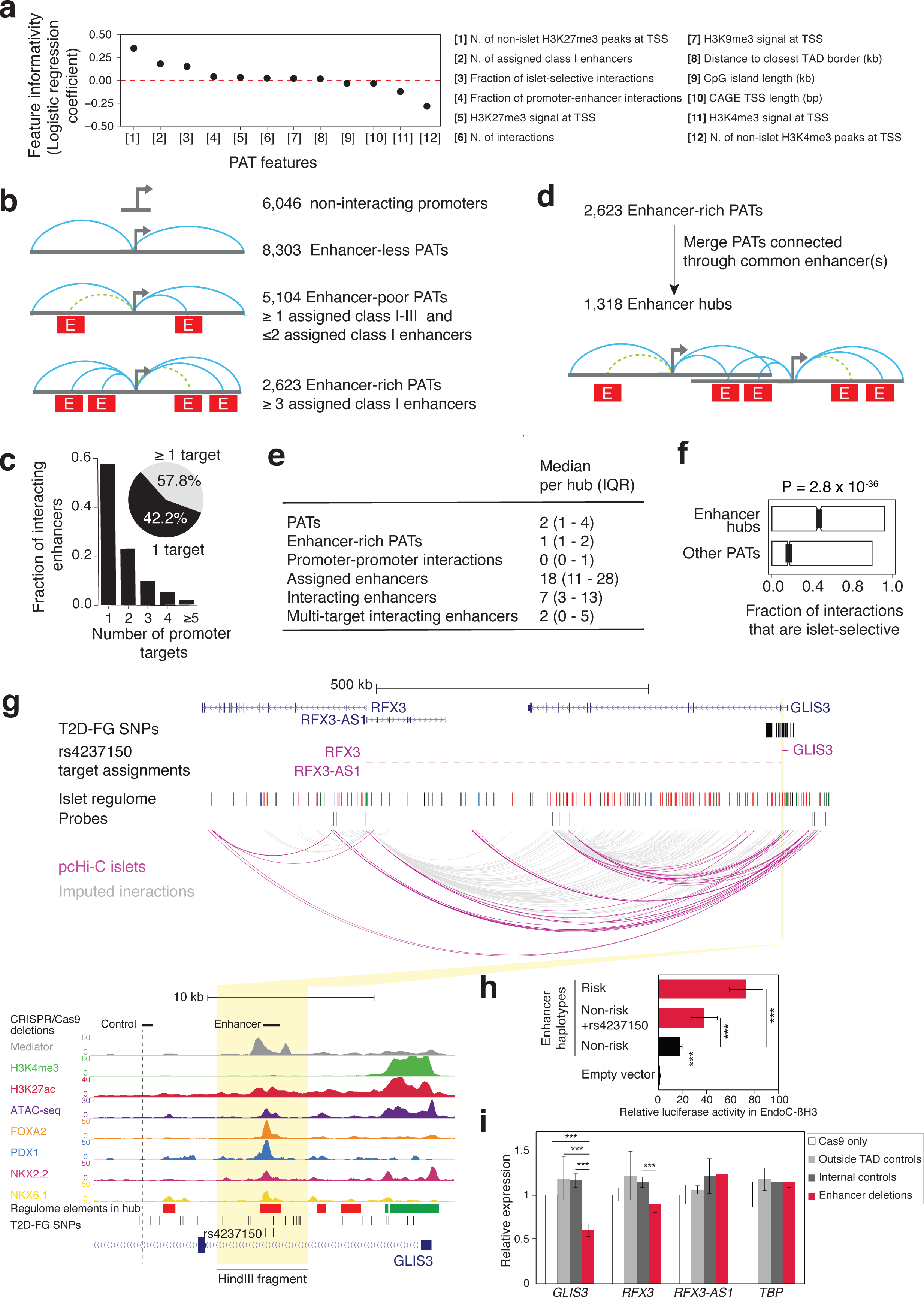
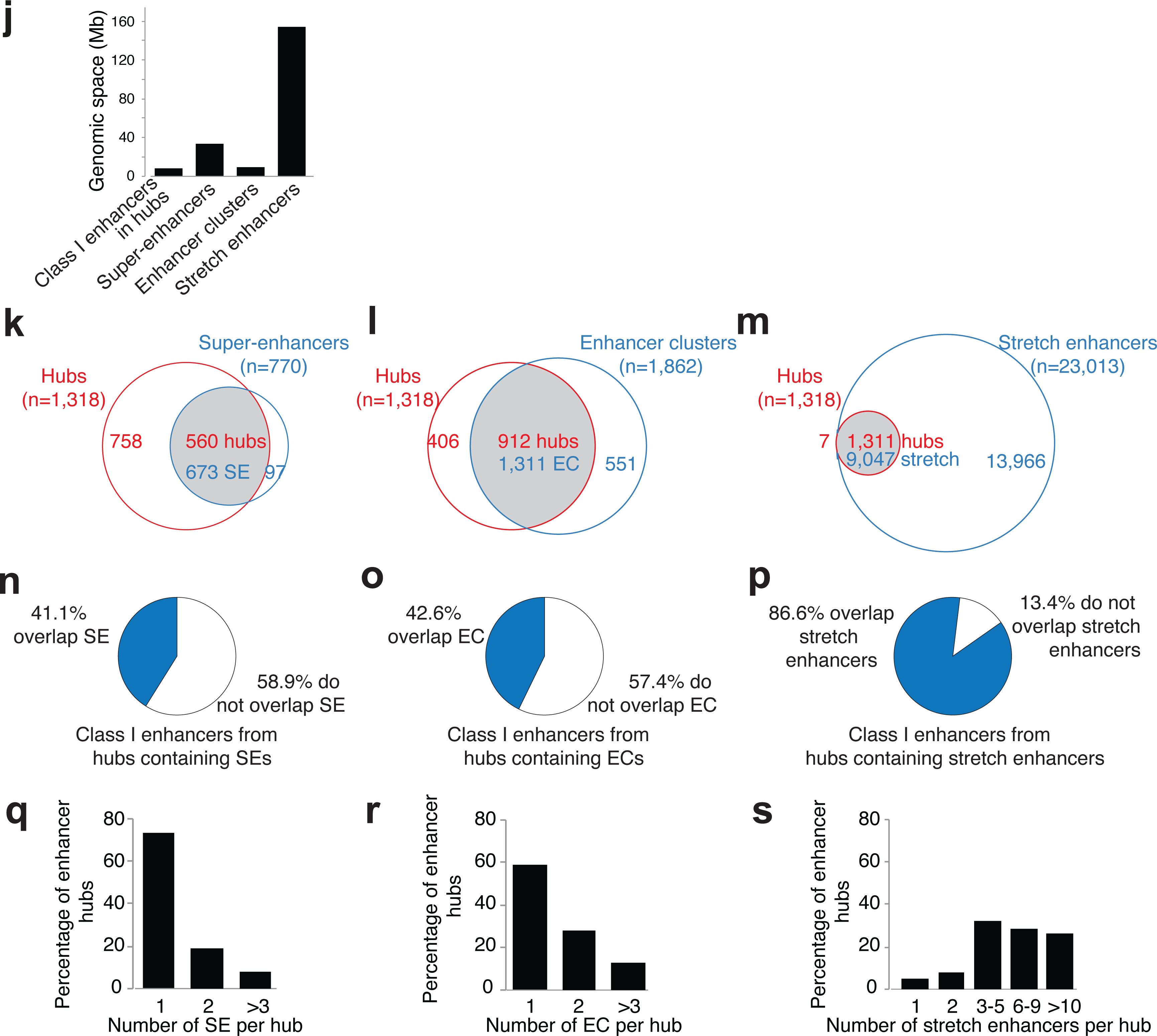
Islet-specific transcription is driven by tissue-specific enhancer hubs. Related to Figure 4. (a) Multiple logistic regression analysis was used to identify PAT features that differentiate islet-expressed genes that show islet-selective vs. non islet-selective expression across tissues. Islet-selective expression was examined as a surrogate endpoint because it is a property of many (though not all) genes that are important for islet cell identity. The PAT feature with the highest logistic regression coefficient was the number of non-islet tissues with promoter H3K27me3-enrichment. This feature was included as a control as it was regarded as nearly synonymous with cell-specific islet expression. The next highest coefficient was the number assigned class I enhancers in the PAT. Further analysis showed that ≥3 assigned class I enhancers in a PAT optimized the prediction of islet-selective expression. (b) Classification of PATs based on assigned enhancers revealed 2,623 enhancer-rich PATs (≥3 assigned class I enhancers). Enhancers are represented as red boxes (E). Turquoise and dashed green lines represent high-confidence interactions and imputed assignments, respectively. (c) Enhancers frequently interact with more than one gene. The graph shows the fraction of enhancers that show high-confidence (CHiCAGO > 5) interactions with 1-5 promoter “baits” in the same TAD. The inset illustrates that 58% of interacting enhancers interact with two or more promoters. (d) Enhancer hubs were defined as enhancer-rich PATs that were merged with other PATs connected through at least one common enhancer-associated high-confidence interaction. (e) Descriptive characteristics of enhancer hubs in human islets. (f) The fraction of all islet high-confidence interactions that are islet-selective was greater in enhancer hubs than in other interacting PATs. Boxes are IQR and P values are from Wilcoxon’s signed rank test. (g) pcHi-C map of an enhancer hub containing the diabetes-associated locus *GLIS3*. The zoomed inset contains the regulatory landscape surrounding the risk haplotype in pancreatic islets, with active chromatin features and binding of pancreatic transcription factors to the enhancer that contains rs4237150 and 3 additional T2D-associated SNPs (rs10116772, rs10814915 and rs6476839). The HindIII fragment that contains the T2D-associated enhancer is highlighted in yellow. This enhancer interacts directly with *RFX3* an indirectly with *GLIS3*; despite that the islet *GLIS3* promoter did not contain a bait in the pcHi-C library, is was connected through high-confidence interactions with other enhancers that in turn interact with the SNP-containing enhancer. Only the GLIS3 gene model containing the active islet promoter is shown for simplicity. (h) Luciferase assay in the human β cell line EndoC-βH3 shows haplotype-dependent activity for the rs4237150-enhancer. Data are presented as mean ± s.d. Two independent experiments were performed in quadruplicates. *** P <0.001, two-tailed Student’s t-test. (i) Analysis of hub genes in EndoC-βH3 cells after deletion of the T2D-associated enhancer, a control region that contains T2D-associated variants with no apparent regulatory function (rs3892354 and rs1574285), or a control region in a locus in a different TAD. Deletions were tested with 2 different guide RNA pairs. Empty vector (Cas9 only) was used as reference in all experiments. Data are presented as mean ± s.d. Two independent experiments were performed in triplicates. Data shown has been normalized by *RPLP0* and is shown relative to the mean levels of the negative control (Cas9 only). *** P <0.001, two-tailed Student’s t-test. (j) Linear genomic space occupied by class I enhancers in three-dimensional enhancer hubs compared with the space occupied by super-enhancers, highly-bound enhancers from linear enhancer clusters, and stretch enhancers. (k-m) Venn diagrams depicting how often enhancers from islet enhancer hubs overlap with other human islet enhancer domains: (k) super-enhancers (SEs) calculated with the ROSE algorithm, (l) highly-bound (top two occupancy quartiles) enhancer clusters (ECs), and (m) stretch enhancers. (n-p) Islet enhancer hubs often contain enhancers that do not form part of SEs or ECs. Charts show the fraction of hub class I enhancers that overlapped SEs, ECs or stretch-enhancers. Note that the genomic space occupied by stretch enhancers is an order of magnitude greater than hubs (panel I). (q-s) Islet enhancer hubs very frequently contain multiple (q) SEs, (r) ECs or (s) stretch enhancers.

**Supplementary Figure 5.**
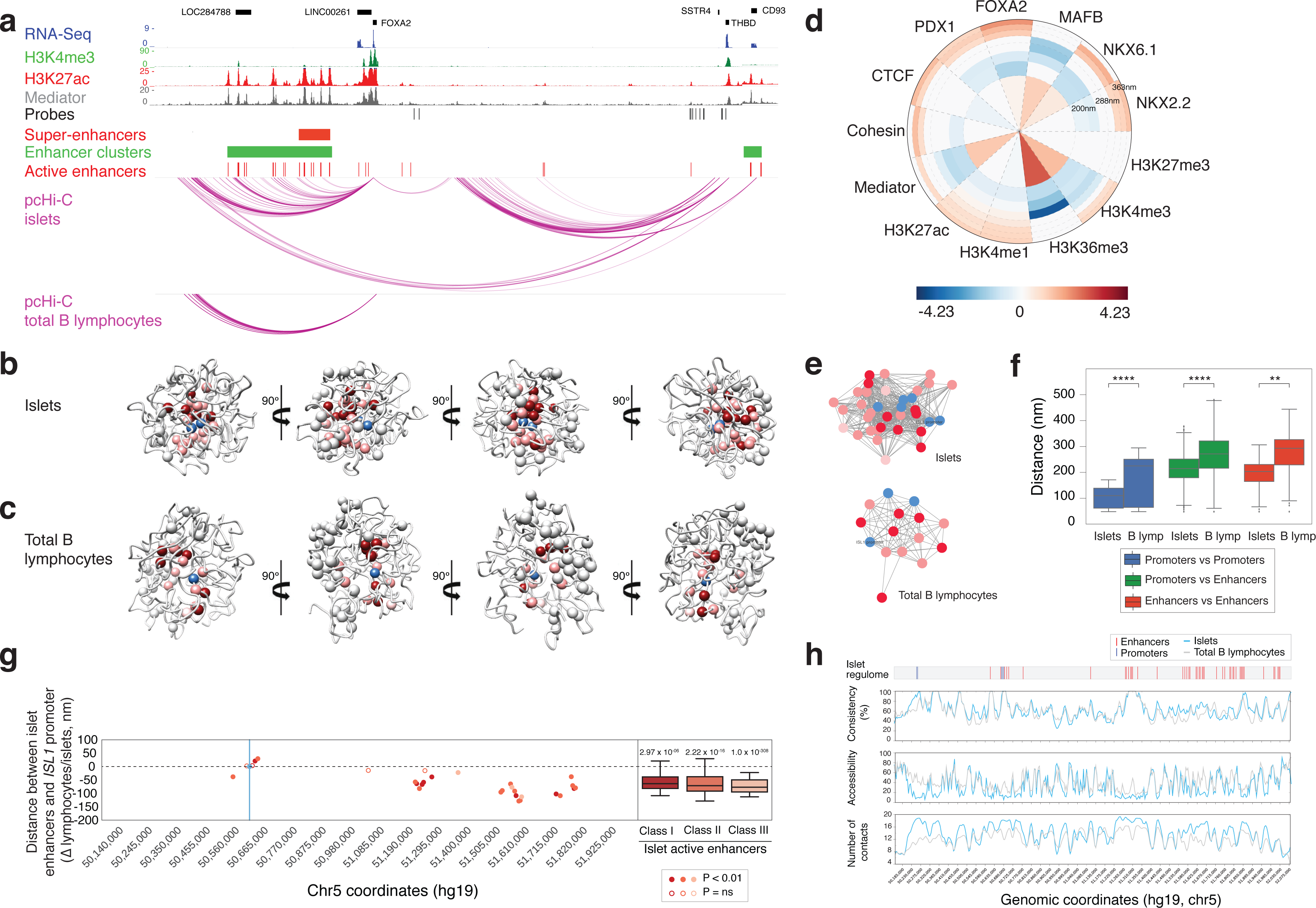
**3D models of enhancer hubs depict tissue-specific regulatory units.** Related to Figure 5. (a) The *FOXA2* locus forms a tissue-specific enhancer hub. Human islet epigenome maps and high-confidence pcHi-C interactions in islets and total B lymphocytes are shown to illustrate that active enhancers, super-enhancers and enhancer clusters interact to form a single tissue-specific three-dimensional structure. (b-c) 360 degree views of top-scoring 3D models of *ISL1* enhancer hub in human islets (b) and total B lymphocytes (c). See also **Supplementary videos 1 and 2**. (d) Density wheel depicting histone modification marks and transcription factor ChIP-seq signal in human islets mapped on the 3D model of *ISL1* enhancer hub in total B lymphocytes (centered on *ISL1* promoter). (e) Network of the first community in the *ISL1* locus revealed through MCODE clustering of the promoter-enhancer mean ensemble distance interaction network in human islets (top) and total B lymphocytes (bottom). Nodes represent promoters (blue) and enhancers (dark to light red corresponding to enhancer classes I to III). Edges between nodes are weighted by the mean distances values in 3D structure ensembles of the *ISL1* enhancer hub in the two tissues. (f) 3D distance distribution between particles in *ISL1* enhancer hub. Distances between enhancers and promoters in the hub are significantly smaller in islets than in total B lymphocytes (see also Figure 5B,C). Statistical significance was computed using two-sample Kolmogorov-Smirnov test (*: p < 5 × 10^−2^, **: p < 10^−2^, ***: p < 10^−3^, ****: p < 10^−4^, and ns: non-significant). (g) Distance measurements between active enhancers and *ISL1* promoter in *ISL1* community in islets expressed as the differential distance between total B lymphocites and islets 3D models. Positive values indicate closer distances in lymphocytes and negative values indicate closer distances in islets. Filled circles represent statistically significant differences (P < 0.01 two-sample Kolmogorov-Smirnov test). The majority of the enhancers (86%) in *ISL1* enhancer hub are significantly change distances in 3D in islets compared to total B lymphocytes. (h) Chromosome view tracks of structure ensemble consistency, local accessibility for a virtual object with radius of 50 nm and local contact density number within spherical volumes with radii 100nm of *ISL1* locus in human islets (turquoise) and total B lymphocytes (grey). Promoter and enhancer locations are highlighted in blue and red, respectively.

**Supplementary Figure 6.**
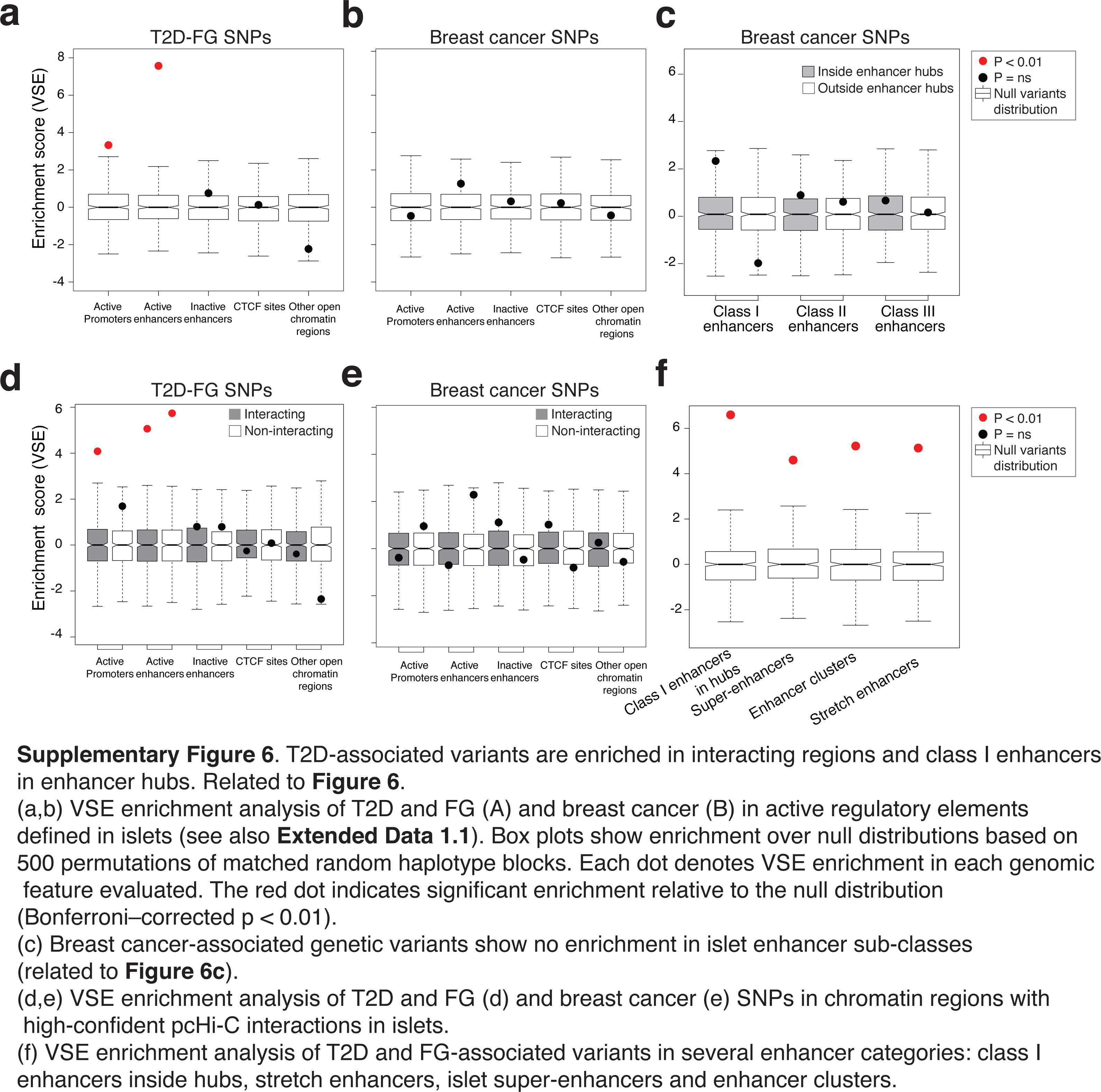
T2D-associated variants are enriched in interacting regions and class I enhancers in enhancer hubs. Related to Figure 6. (a,b) VSE enrichment analysis of T2D and FG (A) and breast cancer (B) in active regulatory elements defined in islets (see also **Extended Data 1.1**). Box plots show enrichment over null distributions based on 500 permutations of matched random haplotype blocks. Each dot denotes VSE enrichment in each genomic feature evaluated. The red dot indicates significant enrichment relative to the null distribution (Bonferroni–corrected p < 0.01). (c) Breast cancer-associated genetic variants show no enrichment in islet enhancer sub-classes (related to Figure 6c). (d,e) VSE enrichment analysis of T2D and FG (d) and breast cancer (e) SNPs in chromatin regions with high-confident pcHi-C interactions in islets. (f) VSE enrichment analysis of T2D and FG-associated variants in several enhancer categories: class I enhancers inside hubs, stretch enhancers, islet super-enhancers and enhancer clusters.

**Supplementary Figure 7.**
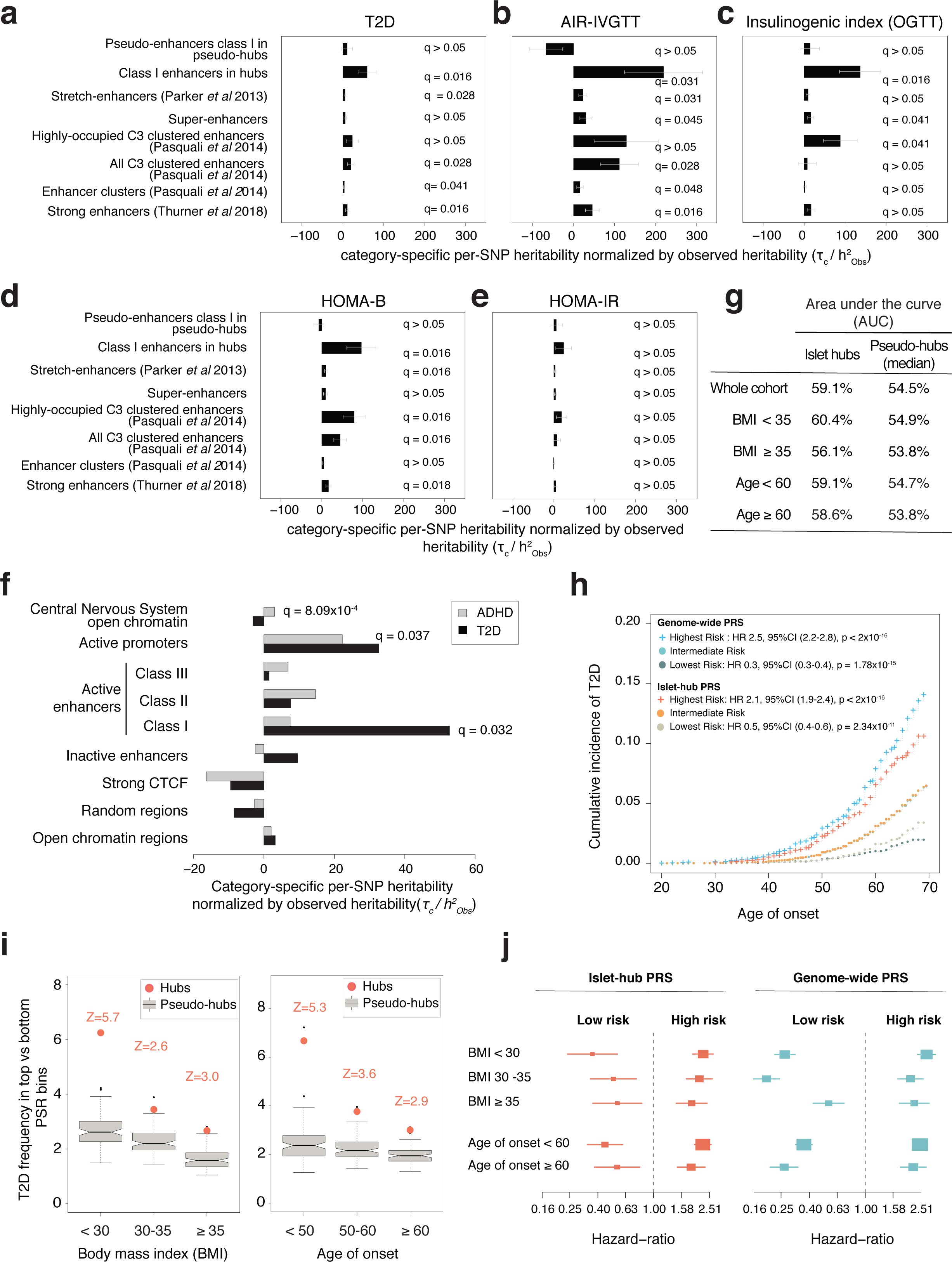
Class I enhancers in hubs contribute to beta-cell related traits heritability. Related to Figure 6. (a-e) Per-SNP heritability estimates of variants in eight islet enhancer domain subtypes for T2D (a), acute insulin release (AIR)-in vivo glucose tolerance test (IVGTT) (b), insulinogenic index (OGTT) (c), HOMA-B (d) and HOMA-IR (e). Bars show category specific per-SNP heritability coefficients (*τ*_*c*_) divided by the LD score heritability (*h^2^*) score observed for each trait. All normalized *τ*_*c*_ coefficients were multiplied by 10^7^ and shown with a SEM. *τ*_*c*_ coefficients were estimated by performing independent stratified LD score regression, controlling for 53 functional annotation categories included in the baseline model. (f) Per-SNP T2D and Attention-Deficit/Hiperactivity Disorder (ADHD) heritability estimates in islet regulatory elements and Central Nervous System (CNS) annotations. Category-specific per-SNP heritability coefficients (*τ*_*c*_) correspond to the expected increase in heritability explained by a single SNP due to the SNP’s being in a given functional category. We normalized *τ*_*c*_ estimates by the observed LD Score heritability (*h^2^*) in each trait respectively, and multiplied by 10^7^. Each *τ*_*c*_ coefficient was calculated by applying independent stratified LD Score regression, controlling for 53 functional annotation categories from the baseline model. Error bars represent the normalized *τ*_*c*_ coefficients ± standard errors. (g) Area-under-the curve (AUC) for islet hub PRS and average AUC across 100 sets of pseudo-hubs PRS. Maximal AUC was also calculated for two ranges of BMI (< 35 and ≥ 35) and age of onset (<60 and ≥ 60). (h) Cumulative incidence of T2D in UK Biobank individuals, based on islet-hub and genome-wide genetic risk groups. Kaplan-Meier survival curves indicate cumulative incidence of T2D across several levels of PRS risk (see legend). We also show hazard ratios per risk group from cox proportional hazard regression analysis on T2D risk with age of onset as underlying time factor. (i) Polygenic risk stratified by BMI (left) and age of onset of T2D (right). The T2D risk ratio was calculated as the frequency of T2D cases in the highest 2.5% PRS bin divided by the frequency of T2D cases in the lowest 2.5% risk bin. T2D cases were stratified using age of onset information and age at recruitment was used in control samples. Boxplots show the risk ratio for PRS from 100 sets of pseudo-hubs, and the orange dot shows that of islet hub PRS. A Z-score of the risk ratio with islet was computed to define the number of standard deviations above the average risk ratio of the pseudo-hub distribution. (j) Forest plot showing hazard ratios (HR) of risk of incident T2D per tails of PRS groups across distinct BMI and age of onset categories. Hazard ratios are denoted by boxes proportional to precision, along with 95% CI error bars

## Data availability

### Sequence reads

Raw sequence reads from pcHi-C, RNA-seq, ChIP-seq, ATAC-seq and 4C-seq reported in this paper will be available from EGA (https://www.ebi.ac.uk/ega) upon publication.

### Processed data files

High-confidence islet pcHi-C interactions, islet regulome annotations, enhancer-promoter assignments, hub coordinates and components and 3D model videos are available upon request and will be made public upon publication.

### Visualization

Data from this study van be visualized in the following browsers:

- Islet regulome browser (http://isletregulome.org/regulomebeta/)^87^, which allows visualization of locus-wide virtual 4C profiles of pcHi-C dataset, alongside with islet regulome annotations including enhancer hubs as well as T2D and FG-associated variants.

- WashU Epigenome browser (http://epigenomegateway.wustl.edu/browser/) allows visualization of ATAC-seq and ChIP-seq signal tracks, regulome annotations, pcHi-C high-confidence interactions and variants associated with T2D and FG can be visualized using the following link:

http://epigenomegateway.wustl.edu/browser/?genome=hg19&session=62hGf7nfcS&statusId=140947077

- CHiCP browser (https://www.chicp.org)^88^, which allows visualization of high-confidence interactions.

**Supplementary Tables:**

**Supplementary Table 1.** T2D and FG-associated genetic variants and predicted target genes.

**Supplementary Table 2.** Functional annotations of islet enhancer hub genes.

**Supplementary Table 3.** Glucose-induced response of hub promoters. **Supplementary Table 4.** List of T2D and FG associated lead variants used for VSE analysis.

**Supplementary Table 5.** Partitioned heritability analysis across functional regulatory elements and islet enhancer types in T2D and FG.

**Supplementary Table 6.** Characteristics of human islet donors and samples. **Supplementary Table 7.** Capture efficiency of Hi-C libraries after HiCUP processing.

**Supplementary Table 8.** ChIP-Seq, ATAC-Seq and RNA-Seq alignment data. Related to Methods.

**Supplementary Table 9.** Oligonucleotide and probe sequences used in this study.

**Supplementary Table 10.** CRISPR-Cas9 sgRNAs used in this study.

**Supplementary Table 11.** Genomic regions used for 3D modeling.

**Supplementary Table 12.** Replication of previously described T2D associations in 15,764 T2D cases and 184,030 control individuals from the UK Biobank population.

### Extended Data 1

**Extended Data 1.1.** Islet regulome: location of islet regulome regions, including enhancers, promoters, and CTCF binding sites.

**Extended Data 1.2.** Enhancer-promoter assignments.

**Extended Data 1.3**. List of T2D-FG associated genetic variants used for functional assessment of chromatin features and gene targets.

**Extended Data 1.4.** List of human islet enhancer hubs.

### Supplementary videos

**Supplementary video 1** – 3D model of *ISL1* locus in human pancreatic islets.

**Supplementary video 2** – 3D model of *ISL1* locus in total B lymphocytes.

### Methods

#### Human islets

Human pancreatic islets from organ donors without a history of glucose intolerance were isolated and purified using established isolation procedures described in ^1^ with local modifications ^2-4^, shipped in culture medium and then re-cultured at 37°C in a humidified chamber with 5% CO2 in RPMI 1640 medium supplemented with 10% fetal calf serum, 100 U/ml penicillin, and 100 U/ml streptomycin for three days before extraction of RNA or fixation for pcHi-C or ChIP-seq, as described below. RNA was extracted from flash-frozen islet pellets using Trizol, following manufacturers’ instructions. For glucose regulation studies, islets were cultured in identical time and medium, except that glucose-free RPMI 1640 medium was supplemented with glucose to achieve final concentrations of either 4 or 11 mM glucose. Donor variables and characteristics of the samples used in this study are provided in **Supplementary Table 6.**

#### Ethics

Islet isolation centers had permission to use islets for scientific research if they were insufficient for clinical islet transplantation following national regulations and ethical requirements and institutional approvals from Leiden University Medical Center, Geneva University Hospitals, University of Lille, and Milano San Raffaele Hospital. Ethical approval for processing chromatin samples from de-identified samples was granted by the Clinical Research Ethics Committee of Hospital Clinic de Barcelona, under registration numbers HCB/2014/0926 and HCB/2014/1151.

#### Hi-C library preparation and capture Hi-C

30-60 million human islet equivalents/donor from four islet donors were cultured as described above for three days prior to fixation in 2% paraformaldehyde (Agar Scientific) at room temperature for 10 minutes with mixing. Fixative was quenched in 125 mM glycine for 5 minutes at room temperature and 15 minutes in ice, and islets were washed twice in PBS. Dry pellets were flash frozen and store at −80C until further usage.

Hi-C libraries were prepared with in-nucleus ligation as described previously ^5,6^. Briefly, chromatin was de-crosslinked and purified by phenol-chloroform extraction and DNA concentration was measured using Quant-iT PicoGreen (Life Technologies). DNA was sheared (Covaris), end-repaired, adenine-tailed and double size-selected using AMPure XP beads. Ligation fragments marked by biotin were immobilized using MyOne Streptavidin C1 DynaBeads (Invitrogen) and ligated to paired-end adaptors (Illumina). The immobilized Hi-C libraries were amplified using PE PCR 1.0 and PE PCR 2.0 primers (Illumina) with 7–8 PCR amplification cycles.

Hi-C libraries were then processed to capture fragments containing annotated promoters, using a previously described design ^6^. Briefly, biotinylated 120-mer RNA baits were designed to the ends of all HindIII restriction fragments that overlap Ensembl-annotated promoters of protein-coding, noncoding, antisense, snRNA, miRNA and snoRNA transcripts, if GC content was 25-65% and <2 consecutive Ns in the sequence within 330 bp of the HindIII restriction fragment terminus. A total of 22,076 HindIII fragments were captured, containing a total of 31,253 annotated promoters for 18,202 protein-coding and 10,929 non-protein genes according to Ensembl v.75 (http://grch37.ensembl.org). Capture was performed with SureSelect target enrichment, using the custom-designed biotinylated RNA bait library and custom paired-end blockers according to the manufacturer’s instructions (Agilent Technologies). After library enrichment, a post-capture PCR amplification step was carried out using PE PCR 1.0 and PE PCR 2.0 primers with 4 PCR amplification cycles. Between 38 and 77 million paired end di-tags were captured from each library (**Supplementary Table 7**).

#### pcHi-C sequence alignment and interaction calling

Raw sequencing reads from the 12 technical sequencing replicates from 4 human islet libraries were processed independently using the pipeline described in^7^, which maps the positions of di-tags against the human genome (GRCh37), filters out experimental artefacts such as circularized reads and re-ligations, and removes duplicate reads. Reads from replicate libraries from each donor were then pooled, and PCR duplicate reads were further removed. Alignment statistics are shown in **Supplementary Table 7.**

Interaction confidence scores were computed with CHiCAGO (Capture Hi-C Analysis of Genomic Organisation)^8^. Briefly, CHiCAGO calls interactions based on a convolution background model reflecting both ‘Brownian’ (real, but expected interactions) and ‘technical’ (assay and sequencing artifacts) components. Interactions with CHiCAGO score >5 were considered high-confidence interactions.

#### ChIP in human pancreatic islets

Related to **Figure 1, Supplementary Figure 1.** The ChIP protocol was adapted from ^9^ with slight modifications as described in ^10^. Between 1,000 and 2,000 human islet equivalents were fixed with 1% paraformaldehyde for 10 minutes at RT. Paraformaldehyde was quenched with 10mM glycine for 5 minutes at RT and cells pelleted at 4°C for 10 minutes at 500g. After washing the sample twice with PBS supplemented with protease inhibitors (Roche), the samples were snap frozen and stored at −80°C until further usage.

Fixed human islets were thawed on ice and subsequently lysed using ice-cold Lysis Buffer (2% Triton X-100, 1% SDS, 100 mM NaCl, 10 mM Tris-HCl pH 8, 1 mM EDTA pH 8.0, 1x protease inhibitor cocktail) for 15 to 20 minutes on ice. Lysed cells were pelleted for 5 minutes at 500 × g at 4C and re-suspended in 130 µL of Sonication Buffer (for Mediator and cohesin ChIPs, 1% Triton X-100, 0.1% SDS, 150 mM NaCl, 20 Mm Tris-HCl pH 8, 2 mM EDTA pH 8.0) or Lysis Buffer (for H3K27ac ChIPs). Chromatin was sonicated using a S220 Focused-ultrasonicator (Covaris) and the following settings: Duty Factor: 2%, Peak Incident Power: 105W, Cycles per Bust: 200, treatment time: 16 minutes. Sheared chromatin was centrifuged at full speed for 15 minutes at 4°C to remove debris and insoluble chromatin. Supernatant was transferred to a fresh low-binding tube to proceed with the ChIP assay and 5% of the lysate was stored to be used as the input sample.

Chromatin was diluted 4 times with ChIP Dilution Buffer (for H3K27ac ChIP 0.75% Triton X-100, 0.1% Na-deoxycholate, 140 mM NaCl, 50 mM HEPES pH8.0,1 mM EDTA, 1x protease inhibitor cocktail) or Sonication Buffer (for Mediator and cohesin ChIPs). Then, 30 µl of pre-blocked magnetic Dynabeads (Thermo Fisher Scientific) were added to pre-clear chromatin by rotation for 1h at 4°C. We added 1 µg of H3K27ac antibody (Abcam ab4729) and 3 µg of rabbit polyclonal antibodies against Mediator (CRSP1/TRAP220 Antibody, A300-793A, Bethyl Laboratories) and cohesin (SMC1 Antibody, A300-055A, Bethyl Laboratories) to the sample and incubated overnight at 4°C while rotating. The next day, 50 µl of magnetic beads were added to the sample and rotated at 4°C for 2h. Beads were subsequently washed at 4C using Low Salt Wash Buffer (1% Triton X-100, 0.1% SDS, 150 mM NaCl, 20mM Tris-HCl, pH 8.0, 2mM EDTA pH 8.0), High Salt Wash Buffer (1% Triton X-100, 0.1 % SDS, 500 mM NaCl,20 mM Tris-HCl pH 8.0, 2 mM EDTA pH 8.0), LiCl Wash Buffer (0.25 M LiCl, 1% NP40, 1% deoxycholate sodium, 10 mM Tris-HCl pH 8.0, 1 mM EDTA pH 8.0), and TE. Beads were eluted in 300 µl of DNA elution buffer at 65°C for 40 minutes with agitation. Beads were placed on a magnet and supernatant was transferred to a fresh tube and de-crosslinked together with input sample following a phenol-chloroform DNA isolation protocol. RT-qPCR was used to evaluate specific enrichment in at least two positive and two negative genomic regions before sequencing.

#### ATAC of human pancreatic islets

Related to **Figure 1.** ATAC-seq library preparations were carried out as described in ^11^ with the following modifications. Fifty human islets were individually selected and washed in ice-cold PBS. Nuclei were isolated by incubating the islets in 300 µl of cold lysis buffer (10 mM Tris-HCl, pH 7.4, 10 mM NaCl, 3 mM MgCl2 and 0.1% IGEPAL CA-630) for 20 minutes on ice with gentle shaking. Nuclei were washed once with 100 µl of ATAC Lysis Buffer and transposition was carried out immediately afterwards using 25 µl 2xTD buffer, 2.5 µl transposase and 22.5 µl nuclease-free water and kept at 37°C for 30 minutes. The reaction was purified using Qiagen’s MinElute Reaction Cleanup Kit.

ATAC libraries were initially amplified for 5 cycles using the following PCR conditions: 72°C 5 min; 98°C 30 s; then cycling at 98°C 10 s, 63°C 30 s and 72°C 1 minute using the following PCR master mix: 25 µl NEBNext High-Fidelity 2X PCR Master Mix, 2.5 µl of 25 µM Forward/Reverse ATAC-seq index primers from Nextera kit (FC-121-1030) and 20 µl of the purified transposed DNA. We used the qPCR plot on ABI7900 (Applied Biosystems) to determine the number of additional cycles of PCR amplification that were required, by using the cycle number that corresponds to 1/3 of the maximum fluorescent intensity. ATAC libraries were purified using Qiagen MinElute PCR Purification Kit and qPCR of open chromatin sites and negative controls were used before sequencing (data not shown).

#### ChIP-seq and ATAC-seq analysis

Related to **Figure 1.**Illumina TruSeq adapters were removed from ChIP-seq reads using cutadapt 1.9.1 (options: −m 20) ^12^ In ATAC-seq reads, low quality bases were trimmed using Trimgalore 0.4.1 (options --quality 15 --nextera), which also removes Nextera transposase adapters (https://github.com/FelixKrueger/TrimGalore). Trimmed reads were aligned to hg19 genome build using bowtie2 2.1.0 (options: --no-unal) allowing no mismatches ^13^. Aligned reads were filtered to retain only uniquely mapped reads (MAPQ>=30) using samtools 1.2 ^14^ and duplicate reads were removed using picard 2.6.0 ^15^. Reads mapping to blacklisted regions ^16^ were also removed using BEDTools 2.13.3 ^17^. For ATAC-seq, reads mapping to mitochondrial genome were also removed. Mapped reads obtained at this stage are referred to as usable reads. The quality of ChIP data was assessed with SPP.R script from phantompeaktools ^18^.

The list of ChIP-seq and ATAC-seq experiments from human islet samples, alignment statistics and accession numbers are shown in **Supplementary Table 8**.

MACS2 was used to determine regions of statistically significant enrichment over corresponding input samples. For ChIP-seq reads from histone modification marks, broad regions of enrichment were called using the options --g hs -- extsize=300 --keep-dup all --nomodel --broad and narrow regions of enrichment were called without using --broad flag. For TF and co-factor ChIP-seq reads, narrow regions of enrichment were called using –g hs –extsize=300 --keep-dup all. For ATAC-seq reads, we used the following options --shift 100 --extsize=200 --keep-dup all --nomodel.

To obtain a robust set of consistent peaks occupied by each TF, co-factor or histone modification mark, we first used MACS2 to call peaks in individual human islet samples with a relaxed stringency threshold (p < 0.01). Then, we pooled all the biological replicates for each mark and identified pooled peaks using a stringent threshold (FDR q < 0.05 for Mediator and cohesin and q < 0.01 for histone modification marks). Finally, we identified a set of consistent peaks when they were present in at least 2 individual human islet samples (out of 3) or at least 3 human islet samples (if we had more than 3 replicates) as well as in the pooled set.

To obtain a robust set of open chromatin sites in human islets, we pooled ATAC consistent peaks from 13 human islet samples. Pooled ATAC peaks that showed multiple sub-peaks in more than 3 human islet samples were manually split, leading to n=241,481 ATAC peaks.

To visualize the data, bigwig files were generated using *bamCoverage* from deeptools (options: −e=300 −-normalizeTo1x 2451960000), related to **Supplementary Figure 1d.**

To assess the tissue-selectivity of islet interactions, we compared high-confidence pcHi-C interactions in human islets with those of 4 human primary blood cell types (erythroblasts, macrophages, naïve CD4^+^ T lymphocytes and total B lymphocytes, from ^6^. We defined “islet-selective interactions” as those that are only present in human islets, “tissue-invariant interactions” as those that are present in islets, as well as in at least 3 out of the 4 human primary blood cell types and “Others” as the rest.

To assess the enrichment of tissue-invariant over islet-selective interactions in specific epigenomic features, we calculated the overlap between extended promoter interacting regions of both categories and epigenomic features of interest, and then normalized by the number of tissue-invariant and islet-selective interactions, respectively.

#### Gene classification based on expression selectivity in human tissues

Related to **Figures S1h, S2m and Figure 4b.** RNA-seq datasets generated in 16 human tissues was obtained from The Human BodyMap 2 Project (www.illumina.com; ArrayExpress ID: E-MTAB-513). RNA-seq datasets from human pancreatic islets and acinar tissue have been previously described ^19,20^

Paired-end reads were aligned using STAR aligner version 2.3.0 ^21^ and a modified version of the hg19 genome in which common SNPs (Global Minor Allele Frequency> 1%) from the dbSNP database 142 were masked ^22^. A maximum mismatch of 10 nucleotides was used, and non-uniquely aligned reads were removed. Quantification of the raw read count was done using HTseq-Count version 0.6.1 with python 2.6.6 and Pysam version 0.8.3 ^23^. Counts were then converted into TPMs ^24^. RNA-Seq alignment statistics and accession numbers are shown in **Supplementary Table 8.**

Islet-selective genes were defined as those showing (a) highest expression selectivity across tissues and (b) highest expression in islets relative to other tissues. To measure the overall tissue selectivity of the expression of genes we calculated a coefficient of variation (C.V.) value among the 16 BodyMap samples, the acinar sample and the average value for human islet samples. To measure gene expression enrichment in pancreatic islets relative to other tissues we computed a Z-score of the average gene expression level in human pancreatic islet samples using the distribution of expression across the 18 samples. We also defined expressed/non-expressed status among all 21,177 baited genes. 12,559 (59.3%) were defined as expressed, and 8,618 (40.7%) as non-expressed if their expression in human pancreatic islets was greater or lower than 1.5 transcripts per million (TPMs). Genes with an expression level 3 times higher in acinar tissue than islets (79, 0.4% of all genes) were considered as likely acinar contaminants and therefore removed. Among the remaining expressed genes, those that fulfilled top quartile values for both the inter-tissue CV and islet-enrichment Z-score were defined as “islet–selective” expressed genes (983, 4.6% of all genes). The remaining expressed genes were classified as expressed, non-islet-specific (11,497, 54.3% of all genes).

For the analysis shown in Supplementary Figure 1h, the Z-score was used as a quantitative measure of islet-specificity of gene expression.

#### Classification of human islet accessible chromatin regions and chromatin states

Related to **Figure 1c.** A set of open chromatin regions (n=249,582) was defined by combining consistent ATAC-seq peaks from 13 human islet samples (n=241,481) and regions that did not show ATAC-seq peaks but showed either Mediator or CTCF binding sites in pooled samples (n=1,319, n= 9,596 respectively) or were bound by at least two of islet transcription factors (n=1,514 ^9^).

We then classified the 249,582 islet open chromatin regions using k-medians clustering of ChIP-seq signal distribution of H3K27ac, H3K4me1, H3K4me3, Mediator, cohesin and CTCF, using human islet samples that were selected based on the greatest signal to noise for these marks. Briefly, −log10 (p-value) signal was calculated for each epigenetic mark using 100 bp bins across a 6 kb window centered on consistent open chromatin regions. K-median clustering was used to classify open chromatin regions into 14 clusters using flexClust ^25^. These 14 clusters were manually merged into 8 major categories based on the enrichment patterns of different chromatin marks as follows: active promoters, active enhancers (class I, II and III), inactive enhancers, regions with strong CTCF binding (I and II), and “inactive” open chromatin regions (open chromatin regions that lacked distinctive enrichments for any of the features that were used for the cluster analysis). Each open chromatin class was ranked by CTCF binding to highlight a small subset of enhancers that are bound by CTCF. Figure 1d shows that enhancer classes I-III show expected H3K27ac and Mediator occupancy profiles in three different human islet samples. Post-hoc analysis showed that human islet transcription start sites were markedly enriched in open chromatin regions classified as active promoters, and to a lesser extent in class I enhancers (Figure 1c). Islet regulome annotations and genomic locations can be found in **Extended Data 1.1**.

#### Aggregation plots for pcHi-C baits and interacting regions

Related to **Figure 1d and Supplementary Figure 1d.** A window of 25 kb around the center point of HindIII fragments containing pcHi-C baits and promoter-interacting regions was selected for histone marks and transcription factor density calculations. The distance density was estimated using Gaussian kernels (python 2.7 function scipy.stats.gaussian_kde) ^26^. The expected distribution was computed using 10 randomizations of the epigenomic feature coordinates across the mappable non-blacklisted genome using *shuffleBed* from BEDTools ^17^, retaining only non-overlapping random regions. Density plots were computed using *computeMatrix* and *plotProfile* tools from Deeptools 2 ^27^.

Aggregation plots of Mediator and H3K27ac ChIP-seq signals from three human islet donor samples were plotted using *computeMatrix* and *plotProfile* tools from Deeptools 2 ^27^. Input DNA was used as a reference.

#### Overlap of interacting regions with epigenomic annotations

Related to **Figure 1e.** The assessment of overlaps between promoter interacting regions and epigenomic annotations was carried out with the CHiCAGO package ^8^ version 1.0.4. To create control interactions, we used the method implemented in the CHiCAGO package to create 100 sets of distance-matched interactions ^8^. Briefly, promoter interacting regions were randomly re-organized across all 22,076 baits such that the distance distribution of the interactions was maintained. The overlaps were also defined through the CHiCAGO package as the number of promoter interacting HindIII fragments that overlap with the genomic regions of interest, using either the set of high-confidence interactions (CHiCAGO score>5), giving us the observed overlap, or the sets of distance-matched control interactions, giving us the expected overlap. For the expected overlaps, a 95% confidence interval of the mean was also obtained.

#### Extension of landing sites

Related to **Figures 1f, 2b and Supplementary Figure 2k-l.** Epigenomic features that showed enrichment in high-confidence interacting regions (e.g. CTCF-, cohesin-, Mediator-bound regions, active promoters and enhancers) also showed increased presence of high-confidence interactions in the immediately adjacent HindIII fragment. To assess this, we considered each epigenomic feature described above, and quantified high-confidence interacting regions in the HindIII fragment that contained the epigenomic feature and adjacent fragments. In parallel, we performed 10 randomizations of the epigenomic feature coordinates across the mappable non-blacklisted genome using *shuffleBed* from BEDTools ^17^, retaining only non-overlapping random regions. This showed that HindIII fragments that were immediately adjacent to the epigenomic features, but do not directly contain the features, show a 2.8 fold enrichment in non-baited promoter interacting regions compared to the number computed using the randomized data. This enrichment was consistent with the expected proximity of adjacent fragments in nuclear proximity ligation assay. We therefore extended the non-baited promoter interacting regions to the immediately adjacent HindIII fragments. Baited promoter interacting regions were not extended, as we did not observe enrichment in adjacent fragments. The interacting sites evaluated in **Figures 1f, 2b and Supplementary Figure 2k-l** were therefore composed of three adjacent HindIII fragments, whilst the primary analysis of overlapping features shown in Figure 1e was carried out with single fragments.

#### Enrichment of tissue-invariant over islet-selective interactions

Related to Figure 1f. The enrichment of tissue-invariant over islet-selective interactions in a specific epigenomic feature 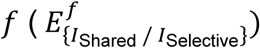 was computed as follows:

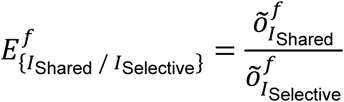

Where *f* is the epigenomic feature of interest, and 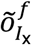 is the normalized overlap of interactions of type x computed as:

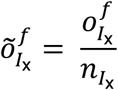

Where 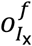 is the number of interactions of type x with extended promoter interacting region overlapping the epigenomic feature *f*, and *n*_*I*x_ is the number of interactions of type x.

### Expression selectivity analysis of islet-selective and tissue-invariant pcHi-C interactions

Related to **Supplementary Figure 1h.** To assess whether islet-selective pcHi-C interactions were preferentially connected with genes showing islet-selective expression, we selected baits that exclusively contained islet-selective or tissue-invariant interactions and computed the gene expression islet specificity Z-score of the genes contained in the baits. Statistical analysis was performed using Wilcoxon’s signed ranked test.

#### 4C-seq Library Preparation

Related to **Supplementary Figure 1j,k.**

4C-seq library preparation of human EndoC-βH1 β cells was performed following the protocol described in ^9^. 4C-seq datasets from native human pancreatic islets in *MAFB* and *ISL1* loci shown in Supplementary Figure 1j,k are taken from ^9^.

Briefly, around 10 million EndoC-βH1 human β cells were crosslinked in paraformaldehyde 2% (Agar Scientific) for 10 minutes at room temperature.

Paraformaldehyde was quenched with 10mM Glycine for 5 minutes at RT and cells pelleted at 4C for 10 minutes at 1,000g. After washing the sample twice with PBS supplemented with protease inhibitors (Roche), the samples were snap frozen and stored at −80C until further usage. Cells were lysed in 50 mM Tris pH=8, 150 mM NaCl, 5 mM EDTA, 0.5% NP-40, 1% TX-100, 1x protease inhibitor cocktail on ice until nuclei were released. Nuclei were then pelleted and digested with 400 units of DpnII (New England Biolabs) overnight. After assessing digestion efficiency, DpnII was inactivated at 65C for 20 minutes and ligation was carried out in 60 units of T4 DNA ligase, 700 µl of ligase buffer (Promega) and 5,7 ml of nuclease-free water overnight at 16°C. Crosslinks were reversed overnight at 65°C using 30 µl of Proteinase K (10 mg/ml) and the next day, after a 30 minutes treatment with RNase A (10mg/ml), DNA was purified using phenol chloroform. Csp6I endonuclease (Fermentas) was used in the second round of digestion using 50 units of the enzyme in a total volume of 500 µl overnight at 37°C. The second ligation was done overnight at 16°C using 100 units of T4 DNA ligase in 15 ml of volume and purified using Amicon Ultra-15 columns (Millipore).

Libraries were prepared by amplifying the concentrated sample with *MAFB* and *ISL1* human promoter-specific primers containing Illumina adaptors described in ^9^. At least 8 independent optimized PCR reactions per viewpoint were pooled for sequencing.

4C-seq libraries were analyzed as described in ^9^. Briefly, 4C-seq reads were sorted, aligned and translated to restriction fragments. A moving average of 30 fragments per window was used to smoothen reads. Next, we calculated for each fragment the Poisson probability of it containing a given number of smoothened reads. To this end, all aligned fragments were randomized 1,000 times in a 2-Mb window centered on the viewpoint and smoothened in the same way. We then defined significant interactions in the 4C-seq experiment as those with a Poisson probability of <1 × 10^−10^.

Virtual interaction profiles (virtual 4C-seq) plots shown depict merged read counts, as computed by CHiCAGO, from islet pcHi-C HindIII fragments that interact with selected bait fragments.

#### **Identification of** islet **TAD-like domains.**

Related to **Supplementary Figure 2a-c.** We defined TAD-like domains by computing genome-wide the Directionality Index (DI) score using the formula proposed by ^28^:

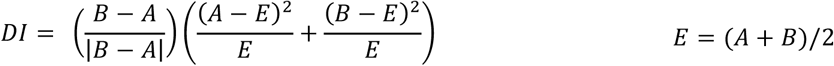

A and B variables from the DI score formula were adapted to pcHi-C as follows:

| Variable | Hi-C (Dixon et al. 2012) | pcHi-C |
| --- | --- | --- |
| A | The number of reads that map from a given 40 kb bin to the upstream 2 Mb. | The number of intra-chromosome interactions going upstream from a sliding window of 5 HindIII fragments. |
| B | The number of reads that map from the same 40kb bin to the downstream 2Mb. | The number of intra-chromosome interactions going downstream from a sliding window of 5 HindIII fragments. |

DI domains were defined as genomic territories flanked by regions with a negative DI score on the 5’ edge and a positive DI score on the 3’ edge.

Interconnectivity between DI domains was computed as a log2 ratio between the number of inter-domain and intra-domain interactions, so that ratios lower than 0 correspond to domains with more intra-domain interaction than inter-domain interactions. Adjacent DI domains with interconnectivity ratios greater than 0 were merged. These merged DI domains defined 3,598 islet TAD-like compartments (see also Supplementary Figure 2a)

#### CTCF occupancy in TAD-like compartments

Related to **Supplementary Figure 2d.** A *de novo* motif analysis was conducted using HOMER ^29^ on a list of consistent CTCF binding sites (see also **ChIP-seq analysis** Methods section), which provided a PWM that matched the known CTCF binding motif. Using annotatePeaks.pl from HOMER, instances of this PWM with a Log score>5 were mapped to islet CTCF binding sites. When multiple motifs were mapped to the same CTCF peak, only the motif with the highest score was kept. We thus determined the position and the orientation of CTCF motifs in consistent CTCF binding sites in human islets. Aggregation plots of CTCF in TAD-like compartments were computed using *computeMatrix scale-regions* and *plotProfile* from DeepTools 2 ^27^.

#### Tissue-specificity of TAD boundary regions

Related to **Supplementary Figure 2e.** TAD boundary tissue-specificity was determined as in ^30^. Islet TAD-like boundaries were defined as 40 kb binned genomic regions overlapping an islet TAD-like edge, similarly to those previously defined by others with Hi-C ^30^. To assess islet-specific TAD boundaries, we examined TAD boundaries from 21 additional tissues. Reference TAD boundaries within 200 kb window were merged into a single TAD boundary “region” using *mergeBed* from BEDTools ^17^. This step was added because TAD boundaries defined in different tissues may be slightly shifted (by a few bins) and borders may be within the same boundary region even if they do not directly overlap (Schmitt et al., 2016). Finally, the number of islet TAD boundary regions overlapping TAD boundaries from the remaining 21 human samples was computed using *intersectBed* (BEDTools).

#### Identification of Promoter-Associated Territories (PATs)

Related to **Supplementary Figure 2f,g.** PATs were created for imputing enhancer-promoter interactions. They were defined as the linear space covered by all the interactions originating from a pcHi-C bait, within the same islet TAD-like compartment. A total number of 16,030 PATs were defined in this manner (see also **Supplementary Figure 3e**).

#### ChromHMM segmentation of regulatory elements

Related to **Supplementary Figure 2h-j.**

To segment the genome into chromatin states based on combinations of chromatin marks in pancreatic islets, we used ChromHMM v1.11 ^31^. Previously published human islet in-house datasets (H3K27me3, H3K36me3, H3K4me1, H3K4me3, H2AZ)^9^, externally generated H3K9ac and H3k9me3 datasets ^32^, and newly generated datasets described in the current study (cohesin, Mediator, CTCF and ATAC) were used. The specific parameters used were as follows: reads were shifted in the 5′to 3′direction by 100 bp. For each of the aligned ATAC-seq and ChIP-seq datasets, read counts were computed in non-overlapping 200-bp bins across the entire genome, which were binarized into 1 (enrichment) or 0 (no enrichment) by comparing the read counts within each bin against a background signal using a Poisson p value threshold of 1 × 10^−4^. Models from 10, 15, 20 and 25 states were trained in parallel. We chose to focus on the 15-state model as it best summarized the pancreatic islet epigenome.

#### ChromHMM enrichment in PATs

Related to **Supplementary Figure 2h,i.** For each PAT we considered the entire linear genomic space, and computed the fraction of genomic space occupied by every ChromHMM state segment. The enrichment of ChromHMM states in PATs was calculated as the fraction of PAT space occupied by a given ChromHMM state divided by the fraction of genomic space occupied by the same state. For 7,085 PATs that were at least 25% smaller than their corresponding TAD-like space, we computed the enrichment of ChromHMM states in the PAT versus the remaining islet TAD-like space.

The analysis of baits was performed separately for bins of baits that differed in their gene expression level. All baits corresponding to an Ensembl promoter in our capture setup were allocated an expression level using the human islet gene expression quantification described above. For baits that contained multiple promoters the average expression level was considered. Baits were then grouped based on their expression levels in 5 bins of equal sizes.

#### Enhancer-promoter assignments

Related to **Supplementary Figure 2k.** We used pcHi-C interactions and PAT features to assign enhancers to promoters contained in interacting baits. We followed a stepwise approach in which each step was only performed on unassigned enhancers from previous steps. For steps 2-4, in which assignments were imputed, we only considered baits that contained at least one active islet promoter, as defined by the islet regulome classification (**Extended Data 1.1**) or ChromHMM analysis. For all assignments, we report candidate target genes with average human islet RNA > 1.5 TPM **(Extended Data 1.2)**. Post-hoc analysis, described in greater detail below, supported imputed assignments based on an increased frequency of moderate-confidence physical interactions (**Supplementary Figure 2l**) and functional correlations (**Figure 2b-d**).

The enhancer-promoter assignment steps were as follows:

1. Enhancers were associated to promoters based on the presence of human islet high-confidence (CHiCAGO score>5) interactions. In cases where the enhancer showed interactions with multiple baits, all genes expressed at > 1.5 TPM were considered. High-confidence interactions that cross TAD boundaries were included for the purpose of enhancer-promoter assignments in this step.

2. If an enhancer did not show high-confidence (CHiCAGO score>5) interactions, we defined the PAT(s) in which it was contained. However, we did not assign the enhancer to all overlapping PATs, because only some active genes are regulated by enhancers; for example, some TADs contain multiple active genes and few or no enhancers. We therefore imputed the assignment of orphan enhancers to overlapping PAT(s) anchored by an active promoter that did show high confidence interactions with other islet enhancers, rather than to other PATs that also contained the enhancer but had no evidence of interacting with any enhancer. Once an enhancer was assigned to a PAT, we selected genes expressed in islets at > 1.5 TPM as candidate gene targets of the enhancer.

3. For remaining enhancers that showed no high-confidence (CHiCAGO score>5) interactions but were contained within 10 kb linear window of a bait containing active promoter(s), we assumed that (a) this linear distance is more likely to provide functional enhancer– promoter communication than promoters located more distally that do not show high-confidence interactions, and (b) random collisions are too frequent to enable the identification of high confidence interactions above background noise with CHiCAGO. We thus imputed these enhancer-promoter assignments, and selected genes expressed in islets at > 1.5 TPM as candidate gene targets of the enhancer

4. In remaining enhancers that showed no high-confidence (CHiCAGO score>5) interactions, but were only contained within the linear genomic space of a single PAT with an active promoter, we imputed the assignment to expressed genes in that PAT bait.

A list of enhancer-promoter assignments can be found in **Extended Data 1.2**.

#### Validation of enhancer assignments

Related to **Figure 2b-d and Supplementary Figure 2l.** Imputed enhancer-promoter assignments were validated by showing (a) increased enhancer-promoter correlations across tissues and human islet samples,

(b) increased islet-specificity of assigned genes, and (c) coordinated changes after exposure to varying glucose concentrations, as described below and (d) higher CHiCAGO scores in the imputed interactions.

#### (a) Enhancer-promoter correlations

Related to **Figure 2b.** We assessed H3K27 acetylation correlation of enhancer and promoter pairs across tissues and human islet samples, based on the assumption that H3K27 acetylation in flanking nucleosomes of genuine enhancer-promoter target pairs should tend to show higher correlation values than unrelated pairs. We found that using only unrelated tissues for this analysis is sometimes not informative because, for example, poor correlation values are expected when an enhancer is exclusively active in one tissue, and the true target gene of that enhancer is expressed in multiple tissues but regulated by different enhancers. This can be complemented by studying correlations across different individual samples from the same tissue, although this is also uninformative in some enhancer-promoter pairs that show limited variation across islet samples. We thus empirically combined tissues and human islet samples to generate a single Spearman’s Rho value for every possible enhancer-promoter pair in each islet TAD, and found that this combined score provided improved discrimination in two functionally characterized loci.

Comparisons of correlation values were made with two enhancer-gene control sets: (i) for every enhancer with an assigned gene promoter, we randomly selected another gene promoter in the same TAD, and (ii) for every enhancer with an assigned gene promoter, we sought a gene promoter from a PAT that also contained that enhancer but was not assigned.

For this analysis, we used 14 human pancreatic islet samples, including 7 samples exposed to 11mM glucose and 4mM glucose, and 51 tissues from Epigenome Roadmap Consortium consolidated epigenomes. Epigenome RoadMap aligned reads from H3K27ac ChIP-seq samples and corresponding inputs were downloaded from egg2.wustl.edu/roadmap/data/byFileType/alignments/consolidated/ and converted to BAM format using *bamToBed* from BEDTools ^17^. To avoid artificial bias in H3K27ac signal strength due to differences in sequencing depth, all datasets selected contained at least 15 million usable reads and were uniformly subsampled to a maximum depth of 30 million usable reads. Active islet enhancers were uniformly extended to +/-750 bp from the center of the peak. Active islet promoters were used without further modification. Promoters were defined as regions that were annotated as such in the islet regulome (**Extended Data 1.1**). The number of reads mapping to active enhancers and promoters from human islets were quantified in all tissues and inputs using *featureCounts* ^33^ and were sequence-depth normalized. Then, sequence-depth normalized input signal was subtracted from the sequence-depth normalized ChIP signal. Spearman’s Rho value is calculated between all pairs of active enhancers and active promoters within TAD-like domains of human pancreatic islets using python’s scipy.stats.spearmanr function.

#### (b) Islet-selectivity of enhancer target genes

Related to **Supplementary Figure 2m.** Islet-selective and expressed non islet-selective genes were defined as described above. We computed the number of islet-selective, non islet-selective but expressed genes, and non-expressed genes among genes that were assigned to an enhancer, and among control genes from PATs that also contain the same enhancer, but were not assigned to the enhancer. The statistical significance was assessed performing a chi-square test (python 2.7 function scipy.stats.chi2_contingency) comparing the frequency of each gene expression class among the two “assigned genes” and “control genes” lists.

#### (c) Concordant glucose regulation of assigned enhancer-promoter pairs

Related to **Figure 2c,d.** H3K27ac ChIP-seq and RNA-seq datasets were analyzed in islets from 7 human donors that were cultured for three days in either 11 or 4 mM glucose, as described above.

To assess glucose-regulation of enhancers, we defined H3K27ac-enriched regions, rather than using annotated enhancers, which typically contain nucleosome-depleted subregions that do not show H3K27ac enrichment. We thus defined consistent H3K27ac-enriched regions for each human donor and each glucose condition treatment using MACS2 with the parameters described above. A total number of 90,814 narrow H3K27ac-enriched regions were interrogated for this analysis. The number of H3K27ac reads mapping to each peak was calculated using FeatureCounts v1.5.0 program (--*ignoreDup* –O --*minOverlap* 10). Then, paired DESeq2 (v1.10.1) analysis was used to assess differential signal strength. Peaks showing differential H3K27ac ChIP-seq signal at adjusted P ≤ 0.05 were then mapped to annotated enhancers.

For RNA-seq of islets exposed to different glucose concentrations, 100 bp paired-end sequencing reads were aligned to masked hg19 genome using STAR aligner v2.3.0 ^21^ (options: --*outFilterMultimapNmax* 1 --*outFilterMismatchNmax* 10). Gene level counts were obtained using FeatureCounts v1.5.0. (-s 2 -p). After removing the genes that did not have at least 5 raw reads mapped in at least 3 replicates, a paired DESeq2 (v1.10.1) analysis was carried out to identify differentially regulated genes. In both cases we calculated the FDR, and chose a value ≤ 0.05 as the significance threshold. This showed that islets exposed to higher glucose concentrations showed a significant increase in RNA levels in 8.4% of all genes, and in in H3K27ac for 7.2% of all annotated enhancers at this significance threshold. Consistent with the notion that some glucose-dependent increase in RNA levels were driven by transcriptional regulatory changes, enhancers and promoters located within 25 kb of glucose-regulated transcripts showed enrichment for significant glucose-induced H3K27ac (q < 10^−3^). A more detailed analysis of glucose-regulated responses across different human islet samples will be presented elsewhere (GA, DR).

To calculate the enrichment of interactions between glucose-induced enhancers and glucose-induced genes (related to Figure 2C), we considered all possible pairs of glucose-induced enhancers (interacting or imputed) and genes with glucose-induced mRNA within an islet TAD-like domain. For each enhancer-promoter pair we created a control pair with a distance-matched baited gene. We excluded experimental pairs when we could not find a distance-matched control. Then, we calculated a Fisher’s exact test p-value to assess if glucose-induced enhancer and genes were enriched in high-confidence or imputed assignments. As an additional control, we assessed if glucose-induced enhancers preferentially contact glucose-repressed genes.

We further examined whether the gene promoters assigned to glucose-induced enhancers also show a significant glucose-dependent increase in H3K27ac levels. As a control, for every glucose-induced enhancer we chose a gene promoter that had the closest distance to the enhancer as the assigned gene promoter. When no control genes were found, only the assigned gene promoter changes without control gene promoters were considered. The median distance for interacting gene promoters and control promoters was 200 kb (IQR 102-356 kb) and 167 kb (IQR 99-351 kb), respectively. The median distance for imputed gene promoters and control promoters was 114 kb (IQR 58-301 kb) and 134 kb (IQR 71-324 kb), respectively.

(d) **Assessment of CHiCAGO scores in imputed assignments**.

Related to **Supplementary Figure 2l.** We considered all imputed promoter-enhancer pairs, as well as the two control sets of promoter-enhancer pairs in the same PAT or TAD described in (a). For each pair, we considered the maximum CHiCAGO score of the interaction between the bait and interacting HindIII fragments.

#### Compilation of T2D-FG associated variants to define putative targets

Related to Figure 3 and **Supplementary Figure 3.** To define the most likely target genes of enhancers that contain putative causal islet regulatory variants we first compiled a comprehensive list of variants associated with T2D and glycemic traits using variants contained in 99% credible sets from (a) the DIAGRAM Metabochip meta-analysis of T2D susceptibility from the DIAGRAM website (diagram-consortium.org/downloads.html)^34^ and (b) the re-analysis of T2D publicly available GWAS datasets and imputation with 1000 Genomes Project (1000G) and UK10K^35^. For loci that were examined in both studies, we considered the union of all variants. Additionally, a list of T2D and Fasting Glycemia-associated lead SNPs was downloaded from the NHGRI-EBI GWAS Catalog on 31/05/2016, from which we selected SNPs if they complied with the following criteria: (a) genome wide significance (p ≤ 5 × 10^−8^) in a study that could be verified in a peer-reviewed publication; (b) the phenotype in the study was reported as Type 2 diabetes, or Type 2 diabetes (and other traits), for T2D-associated SNPs, “Fasting glucose-related traits” or “Fasting plasma glucose”, and (c) they were not identified in or were in high linkage disequilibrium (LD) with SNP in the credible sets from ^34^ and/or ^35^. For this set of lead SNPs, we identified variants in high LD (*r^2^* > 0.8) using PLINK ^36^ and 1000 Genomes Project phase3 data for CEU samples (European loci), CHB and JPT samples (Asian loci) or YRI samples (African loci; Yoruba from Ibadan, Nigeria), in the original populations were the original or replications GWAS studies were carried out. A schematic of the compilation process can be found in **Supplementary Figure 3A** and the full list of genetic variants can be found in **Extended Data 1.3**. Summary statistics from ^35^ and ^34^ can be found at: http://cg.bsc.es/70kfort2d/ and http://diagram-consortium.org/downloads.html respectively.

#### Identification of potential target genes of T2D-FG associated variants

Related to **Figure 3.** We integrated the list of T2D/FG-associated variants with enhancer-promoter assignments to identify putative candidate target genes of T2D-FG-associated variants. In total, we were able to associate 530 enhancer variants from 54 loci to islet-expressed genes using high-confidence interactions and imputations (**Figure 3a**). **Supplementary Table 1** provides a more extensive list of 830 T2D and/or FG-associated variants that overlap an active enhancer or a promoter, which includes information on connections to candidate genes through (a) high-confidence interactions (CHiCAGO score >5), (b) moderate-confidence CHiCAGO interaction scores (2.5-5), (c) imputation, (d) indirect connections through a common hub, and (e) location of actively expressed gene within 10 Kb of disease-associated enhancer variants, regardless of whether they were imputed, given the limited power to discern random collisions from regulatory interactions in this range. This category also included actively transcribed genes from associated variant-containing promoters that overlap pcHi-C baits.

**Supplementary Table 1** additionally lists T2D-FG genetic variants that overlap a promoter interacting region (CHiCAGO score >5) even if it does not overlap a regulatory element, and reports actively transcribed putative target genes.

#### Experimental validation of T2D GWAS variant assignments to target genes

Related to **Figure 3c,d, Supplementary Figures 3 and 4g.i.**

#### Cell lines

Related to **Figure 3c,d and Supplementary Figures 3 and 4g.i.** EndoC βH3 cells ^37^ were maintained in DMEM low glucose (1 g/L) supplemented with GlutaMAX and sodium pyruvate (Thermo Fisher), 2% albumin from bovine serum fraction V (Roche), 50 µM β-mercaptoethanol, 10mM nicotinamide, 5.5 µg/mL human transferrin, 6.7 ng/mL sodium selenite, 100 U/mL penicillin and 100 µg/mL streptomycin. Culture dishes were pre-coated with DMEM (4.5 g/L glucose) 2 µg/ml fibronectin (Sigma) and 1% ECM (Sigma) overnight at 4°C.

HepG2 were obtained from ATCC and maintained in DMEM supplemented with 10% FBS, 2mM L-glutamine, 1mM sodium pyruvate 100 U/mL penicillin and 100 µg/mL streptomycin.

293FT cells were obtained from Thermo Fisher and maintained in DMEM supplemented with 10% FBS, 0.1mM MEM non-essential amino acids, 2mM L-glutamine and 1mM sodium pyruvate, 500 µg/ml geneticin, 100 U/mL penicillin and 100 µg/mL streptomycin.

#### Luciferase reporter assays

For the allele-specific reporter assays presented in Supplementary Figure 3c and Supplementary Figure 4h, the genomic regions containing T2D variants were amplified from human genomic DNA and cloned into the pGL4.23 vector (Promega) at KpnI and HindIII, upstream of a minimal promoter and the Firefly luciferase coding sequence (CDS), using Gibson Assembly (New England Biolabs) ^38^. In the cases where both haplotypes could not be amplified from genomic DNA, site directed mutagenesis was performed using Q5 Site-Directed Mutagenesis Kit (New England Biolabs). Regions of similar size lacking evident regulatory marks in human islets were used as negative controls in addition to the empty pGL4.23 vector. Oligonucleotide sequences used for enhancer cloning and mutagenesis are listed in **Supplementary Table 9**.

Enhancer reporter construct transfections were carried out on 120,000 EndoC βH3 cells in 48-well plates with 0.1pmol of tested construct and 10ng of *Renilla*-expressing pRL vector, by reverse transfection using Lipofectamine 2000 (Thermo Fisher). Luciferase activity was measured 48 hours post transfection with Dual-Luciferase Reporter Assay kit (Promega) on a GloMax-Multi Microplate Multimode Reader. Firefly luciferase measurements were normalized to *Renilla* luciferase. Two experiments with three to four independent transfections were performed per tested enhancer and data are represented as the fold change in relative luciferase signal over the average activity of the negative controls with S.D. Two-sided Student’s *t* test was used to calculate significance.

#### Generation of CRISPR-Cas9 delivery vectors

Related to **Figure 3c,d and Supplementary Figures 3 and 4g.i.** For deletion of genomic regions in the EndoC βH3 cell line, which is resistant to puromycin ^37^ and responds poorly to cell sorting, we replaced the EGFP CDS from pSpCas9(BB)-2A-GFP (pX458 plasmid #48138, Addgene, ^39^) with a hygromycin resistance CDS, generating the pSpCas(BB)-2A-HygR backbone which allows positive selection of EndoC βH3 cells after transfection (plasmid to be deposited, Addgene). The pSpCas(BB)-2A-HygR vector was generated by Gibson assembly using primers listed in **Supplementary Table 9**, and PCR amplification of the hygromycin resistance CDS from lentiMPH v2 (plasmid #89308, Addgene^40^).

#### Guide RNA design and cloning for CRISPR-Cas9-mediated deletions

Related to **Figure 3c,d and Supplementary Figures 3 and 4g.i Figure 3c,d and Supplementary Figures 3 and 4g.i.** We used Cas-Designer (http://www.rgenome.net/cas-designer/) to design 17nt-long guide RNAs (gRNAs).Truncated gRNAs have been shown to be more specific without a significant compromise of the targeting efficiency ^41^. Deletions were achieved by delivery of pairs of gRNAs, with one guide on each side of the target region (Left and Right guides). For each target region, we designed up to 4 different deletions, each with a different “Left+Right” combination of guides. A list of all guide RNAs used in this study can be found in **Supplementary Table 10**.

To increase the efficiency of CRISPR-Cas9-mediated deletions, we used a dual gRNA cloning strategy, which allowed combination of different Left+Right pairs with the same set of oligonucleotides. To this end, a plasmid containing a sgRNA scaffold, a H1 promoter, and a kanamycin resistance CDS (pScaffold-H1) (plasmid to be deposited, Addgene), was generated by TOPO cloning (Thermo Fisher)^42^. The sgRNA scaffold and H1 promoter sequences were amplified from pDECKO-mCherry GFP (plasmid #78535, Addgene, gift from Roderic Guigo & Rory Johnson^43^)(primers listed in **Supplementary Table 9**). PCR primers for gRNA cloning were as follows:

Primer: Sequence
Forward: CGA**GAAGAC**CTcaccgNNNNNNNNNNNNNNNNN*GTTTTAGAGCTAG AAATAGCAA* (**BbsI**/sgRNA1/*sgRNA scaffold*)
Reverse: GGT**GAAGAC**CCaaacNNNNNNNNNNNNNNNNN*GGGAAAGAGTGGT CTCA* (**BbsI**/sgRNA2 reverse complement/*H1 promoter*)

Oligonucleotides corresponding to deletion-yielding pairs of gRNAs were assembled in a reaction mix containing 1X Q5 reaction buffer, 200µM dNTPs, 0.5 µM primer, 0.02U/µl Q5 high-fidelity DNA polymerase and 0.25ng/µl of pScaffold-H1 vector, with T_a_=58°C, 15 sec extension, for 30 amplification cycles. PCR products were digested and ligated to pSpCas(BB)-2A-HygR in a one-step digestion ligation reaction with 1X tango buffer, 1µl FastDigest BbsI (Fermentas), 1µl 0.1M DTT, 1µl 10mM ATP, 0.5µl T7 ligase, 100ng of vector and 1µl of PCR product (diluted 1:20). Digestion-ligation reactions underwent 6 cycles of 5 min at 37°C and 5 min at 23°C, and final 5 min incubation at 37°C, after which 2µl of the final ligation were used to transform 25µl of Stbl3 chemically competent *E. coli*. On the following day, single colonies were picked and analyzed by Sanger sequencing. Plasmids were isolated using a midiprep kit (Promega) and concentrated to 1-2 µg/µl by ethanol precipitation.

#### Enhancer deletions with CRISPR-Cas9

Related to **Figure 3c,d and Supplementary Figures 3 and 4g.i.** EndoC βH3 cells were nucleofected in a Nucleofector B2 (Lonza) following the manufacturer’s protocol, using 2 million cells and 10 µg plasmid DNA per nucleofection with Amaxa cell line nucleofection kit V (Lonza) and program G-017. After nucleofection, 500 µl RPMI media was added to each cuvette. After a 20 min incubation, cells were transferred to a pre-coated 12-well plate containing pre-equilibrated antibiotic-free EndoC βH3 media. Media was changed 18 hours post nucleofection and hygromycin selection was started 6 hours later (200µg/ml). Hygromycin was replenished 72 hours post nucleofection. After a period of 5 days of selection, media was changed daily using 50% conditioned EndoC βH3 media. Cells carrying deletions were passaged every 5-6 days into fresh pre-coated plates similarly to wild type cells. CRISPR-Cas9-mediated deletion efficiency was assessed by PCR, using primers outside the deleted regions (primers listed in **Supplementary Table 9**). Genomic DNA was extracted 5-6 days post-nucleofection.

#### CRISPR inhibition vectors

Related to **Supplementary Figure 3d,e.** To minimize prolonged selection of EndoC βH3 cells with multiple antibiotics, we devised an all-in-one CRISPRi system containing the gRNA expression cassette (U6 promoter, gRNA protospacer and sgRNA scaffold) and the CRISPRi cassette (KRAB domain fused to dCas9, 2A peptide and blasticidin resistance CDS). Two versions of the system were generated: Lenti-(BB)-EF1a-KRAB-dCas9-2A-BlastR (plasmid to be deposited, Addgene) and Lenti-(BB)-hPGK-KRAB-dCas9-2A-BlastR (plasmid to be deposited, Addgene), containing core EF1a or hPGK promoters, respectively. The Lenti-(BB)-EF1a-KRAB-dCas9-2A-BlastR vector was generated by Gibson assembly (NEB, PMID 19363495), using the backbone of lentiCRISPR v2 (plasmid #52961, Addgene, ^44^), and amplifying the KRAB-dCas9 cassette from pHR-SFFV-KRAB-dCas9-P2A-mCherry (plasmid #60954, Addgene, ^45^), and the blasticidin resistance CDS from lentiSAM v2 (plasmid #75112, Addgene, ^40^) using primers listed in **Supplementary Table 9**. The Lenti-(BB)-hPGK-KRAB-dCas9-2A-BlastR was generated by swapping the promoter of Lenti-(BB)-EF1a-KRAB-dCas9-2A-BlastR by hPGK, which was amplified from human genomic DNA with primers listed in **Supplementary Table 9**. hPGK-containing backbone yielded stronger KRAB-dCas9 expression in EndoC βH3 cells and was thus selected for our CRISPRi experiments.

#### CRISPR-mediated inhibition (CRISPRi) and activation (CRISPRa) vectors

Related to **Supplementary Figure 3d,e.** gRNAs targeting transcriptional start site regions were retrieved from the Human Genome-wide CRISPRi-v2 and CRISPRa-v2 top5 libraries, respectively ^46^, and previously validated negative control gRNAs targeting sequences not present in the human genome were retrieved from Addgene (https://www.addgene.org/crispr/reference/grna-sequence/). To induce formation of heterochromatin (CRISPRi) or further activate (CRISPRa) enhancer regions, 20nt-long gRNAs against the core of the enhancer (summit of ATAC-seq/Mediator ChIP-seq signal) were designed using Cas-Designer (http://www.rgenome.net/cas-designer/). guide RNAs used in this study and associated references can be found in **Supplementary Table 10**. gRNA oligonucleotides were cloned into the Lenti-(BB)-hPGK-KRAB-dCas9-2A-BlastR and SAMv2 ^40^ vectors, respectively, following a previously described protocol ^44^. Briefly, oligonucelotides (Thermo Fisher) containing gRNA sequences flanked by BsmBI-compatible overhangs were phosphorylated with T7 polynucleotide kinase (NEB) and annealed. Fragments were ligated into BsmBI-digested destination vector. Ligated constructs were transformed into Stbl3 chemically competent *E. coli* and clones were checked by Sanger sequencing.

#### Lentiviral production

Related to **Supplementary Figure 3d,e.** CRISPRi and CRISPRa lentiviral particles were produced following a previously described lentivirus production protocol ^47^. 293FT cells were seeded at 75,000 cells/cm2 in T75 flasks. 24 hr later when cells reached a confluence of 80%, cells were transfected with gRNA-containing CRISPRi or CRISPRa vectors together with third generation packaging plasmids pMDLg/pRRE, pRSV-Rev and pMD2.G (plasmids #12251, #12253 and #12259, Addgene), using PEI-Pro (Polyplus-transfection) according to manufacturer’s instructions, in antibiotic-free media using a 1:1 ratio of total DNA µg to µl of PEI-Pro. Transfection efficiency was estimated 24 hours later by microscopic inspection of a control EGFP-expressing vector (pLJM1-EGFP) (plasmid #19319, Addgene), which was transfected in parallel. Media was replaced by 9ml of fresh 293FT antibiotic-free media 18 hours post transfection and lentiviral particles were collected 72 hours post transfection. Immediately after collection, supernatants were spun down for 5 min at 1500 rpm, filtered using Steriflip-HV, 0.45µm, PVDF filters (Millipore), supplemented with 1mM MgCl_2_, and treated with 1µg/ml DNaseI (Roche) for 20 minutes at 37°C. Virus particles were then concentrated by overnight incubation with 3ml of Lenti-X Concentrator (Clontech) at 4°C. On the following day, virus particles were collected by cold centrifugation for 45 min at 2500 rpm, resuspension in 90 µl of PBS, aliquoting and storage at −80°C until use.

#### Enhancer modulation with CRISPRi or CRISPRa

Related to **Supplementary Figure 3d,e.** EndoC βH3 and HepG2 cells were transduced during passage with addition of 10µl and 1µl of concentrated lentiviral supernatant, respectively. 24 hr post-transduction media was changed and 48 hr later selection was started with blasticidin (EndoC βH3 8µg/ml, HepG2 cells 3µg/ml). Medium was replenished every 48 hr thereafter, until all negative control cells were dead (usually 8-9 days). After selection, cells were passaged and used for RNA extraction.

#### Quantitative gene expression analysis

Related to **Figure 3c,d and Supplementary Figures 3 and 4g.i.** EndoC βH3 cells carrying deletions were seeded at 0.5 million cells/well in 12-well plates and treated with 1 mM 4-Hydroxytamoxifen for 3 weeks, replenishing 4-Hydroxytamoxifen treatment twice per week, to induce oncogene excision^37^, and passaged every 5-6 days. EndoC βH3 cells transduced with CRISPRi or CRISPRa lentiviral particles were seeded at 0.5 million cells/well in 12-well plates and cultured for 48 hours prior to RNA harvesting. Total RNA was extracted with RNeasy Mini Kit (Qiagen) using in-column RNase-Free DNase Set (Qiagen). 500 ng of total RNA were used for reverse transcription with SuperScript III First-Strand Synthesis SuperMix, using 1 mM dNTPs, 10 µM random primers and 1 unit/µl SUPERase In RNase Inhibitor (Thermo Fisher).

Quantitative PCR was performed with Universal Probe Library assays (UPL, Roche), designed with the Universal ProbeLibrary Assay Design Center. 5 µl reactions were carried out in duplicates, in a QuantStudio 12K Flex (Applied Biosystems), with 1x TaqMan Fast Advanced Master Mix (Thermo Fisher), 1 µM forward and reverse primers, and 250 nM of UPL probe. Quantification was performed using the standard curve method with 5 points of 5-fold serial dilutions. Relative gene expression was calculated by normalizing to the housekeeping gene *RPLP0*. Primers and probes are listed in **Supplementary Table 9.**

#### Logistic regression analysis of PAT features associated with islet-selective RNA expression

Related to **Supplementary Figure 4a.** We used logistic regression to identify PAT features that were the best predictors of islet-selective vs. non-selective expression amongst islet-expressed genes. After removal of highly correlating features (pairwise Pearson’s correlation >0.65) we analyzed the features shown below as independent variables for multiple logistic regression analysis:

### Epigenomic features interrogated in logistic regression analysis

- H3K4me3 signal at TSS in human pancreatic islets

- H3K27me3 signal at TSS in human pancreatic islets

- H3K9me3 signal at TSS in human pancreatic islets

- Number of H3K4me3 peaks at TSS across 139 tissues

- Number of H3K27me3 peaks at TSS across 139 tissues

- TSS length determined by CAGE in human islets (bp)

- CpG island (CGI) length (kb) in promoter

- Number of islet pcHi-C interactions in PAT

- Fraction of PAT interactions that are islet-selective

- Fraction of all PAT interactions that are promoter-enhancer interactions

- Distance of promoter to the closest TAD border (kb)

- Number of islet-assigned class I enhancers

Logistic regression (LR) was implemented using python 2.7 and scikit-learn^48^ with *class_weight* mode set as “balanced” to automatically adjust weights in an inversely proportional manner to the class frequencies in the input data. The machine learning classifier was trained using 70% of the full dataset with 250 iterations and random sampling. The remaining 30% of the dataset was used for validations. In each iteration, the machine learning classifier computed the logistic regression coefficients, {*β*_*i*_}_*i*∈⟦1;n⟧_, for a list of *n* epigenomic features listed above, {χ*_i_*}*_i∈⟦1;n⟧_*, that fit a logit function.

The logistic regression model can be written as follows:

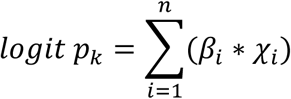

where *p* is the probability that a given gene belongs to a certain gene expression class (*k*).

#### Classification of PATs based on enhancer content

Related to **Supplementary Figure 4B.** Logistic regression analysis showed that the number of class I enhancers assigned to a PAT was independently predictive of islet-selective expression of the PAT genes. Further analyses showed that this effect was apparent with PATs with ≥3 assigned class I enhancers. The only more enriched feature among PATs showing islet-selective expression was the number of non-islet tissues enriched in H3K27me3, however we regarded this as not informative inasmuch as it is a proxy for islet-selective gene activity. We therefore used the number of assigned class I enhancers to classify PATs as follows:

Enhancer-less PATs, without any assigned enhancer (n= 8,303).

Enhancer-poor PATs, with ≥1 assigned class I-III enhancers, but ≤2 assigned class I enhancers (n= 5,104).

Enhancer-rich PATs, with ≥3 assigned class I enhancers (n= 2,623).

#### Definition and analysis of enhancer hubs

Related to **Figure 4.** Enhancer-rich PATs were frequently interconnected through one or more enhancers that interacted with more than one baited promoter (42.4% of all active enhancers had high-confidence interactions with >1 bait). We thus merged enhancer-rich PATs with other PATs that were connected by one or more common enhancers through high-confidence interactions (CHiCAGO score>5) with multiple baits. This resulted in 1,318 enhancer hubs (**Extended Data 1.4**). For 99.5% of hubs all of hub components were restricted to one chromosome.

For the purposes of annotating promoters that form part of hubs, we considered all annotated promoters that were associated with median RNA expression >1.5 TPM in human islets (**Extended Data 1.4**). In few cases (n=426), pcHi-C bait HindIII fragments contained active enhancers, which were shown to establish high-confidence pcHi-C interactions with non-baited fragments that contained active islet promoters. These enhancer-promoter interactions and the active promoters were also considered valid constituents of islet hubs. A list of human islet enhancer hubs is presented in **Extended Data 1.4**.

#### Enrichment in islet-specific expression in hubs

Related to **Figure 4b.** We computed islet-selective, non-islet-selective but expressed genes, and non-expressed genes in hubs as in **Figure 2**, and calculated ratios relative to all genes. Statistical significance was assessed with hypergeometric tests.

#### Enrichment in functional annotations in hub genes

Related to **Figure 4c.** Functional enrichments of hub genes were performed with Enrichr ^49^. Complete results for relevant categories are in **Supplementary Table 2**. As input, we used Ensembl genes from hub baits with average expression in human islets > 1.5 TPM.

#### Promoter-promoter correlations of hub genes

Related to **Figure 4d.** We selected gene pairs whose promoters were in different baits from the same hub. For each hub gene pair, we selected control pairs such that one gene is one of the hub genes and the other is outside the hub but in the same TAD, and another in which both genes outside the hub but in the same TAD. Only genes with islet RNA expression above 1.5 TPMs were considered for this analysis. To avoid potential bias caused by gene pair proximity, gene pairs were binned in 50 Kb windows, and we ensured that there were a similar number of genes present in each bin for each of the three categories. Spearman correlations were computed by combining RNA-seq data from Epigenome RoadMap ^32^ and human islet samples. Significance for comparisons between the three sets of Rho values was computed using kruskal function from the scipy.stats python library.

#### Enhancer-promoter correlations

Related to **Figure 4e.** All enhancer-promoter pairs in islets within a TAD-like domain were classified in three categories: both elements inside the hub, both elements outside the hub, or one element inside and one outside the hub. In all cases we examined sequence-depth normalized H3K27ac in annotated enhancers and promoters across tissues and islet samples as described for **Figure 2b**, and calculated Spearman’s Rho correlation values. Statistical significance of Spearman’s Rho distributions was assessed using the Kruskal function from scipy.stats.

#### Coordinated glucose response in enhancer hubs

Related to **Figure 4f,g.** To test if different enhancers from the same hubs behave coordinately in response to changes in glucose concentrations, we ranked all enhancer hub gene promoters by the p-value (as estimated by DESeq2) with the sign of the fold-change. For each ranked hub promoter, plotted the distribution of H3K27ac fold change values from individual enhancers in that hub, and calculated median and IQR values. Median and IQR values were then represented as a running average with a window size of 50.

#### Quantification of islet-selective interactions in hubs

Related to **Supplementary Figure 4f.** Baits associated to enhancer-hubs show a higher fraction of islet-selective interactions than baits that have ≥5 islet high-confidence interactions but were not classified as hubs. For each hub or control PAT the fraction of islet-selective interactions was computed as the ratio between the number of islet-selective interactions versus the total number of interactions in the same baits. Statistical analysis was performed using Wilcoxon’s signed ranked test.

#### Super-enhancers, stretch enhancers and enhancer clusters

Related to **Supplementary Figures 4j-s, Figure 6** and **Supplementary Figure 6.**

We defined super-enhancers in human islets using ROSE (https://bitbucket.org/young_computation/rose) ^50^ with a transcription start site exclusion zone size of 5 kb (“-t 2500”) and the default stitching size of 12.5 kb. All active islet enhancers from Figure 1c were used as input constituent enhancers and input-subtracted Mediator ChIP-seq signal from a single donor that showed the highest signal to noise was used for ranking the stitched regions. We used the stretch enhancer annotations from ^51^ and enhancer cluster annotations defined by ^9^. We analysed all enhancer clusters or the 1,862 enhancer clusters with high islet TF occupancy (top 50 percentile of islet TF occupancy) as defined in ^9^.

### 3D modeling of enhancer hubs and analysis

Related to **Figures 5** and Supplementary Figure 5.

#### Matrix filtering, normalization and 3D modeling

Related to Figure 5b,c and **Supplementary Figure 5b,c.** Aligned HiCUP pcHi-C files from the 4 human islet samples and total B lymphocytes ^6^ were binned at 5 kb resolution taking into account the size limit of fragments length distribution in the analyzed regions (**Supplementary Table 11**). Next, two filters were applied to the binned matrices that aimed at removing columns of poor coverage. First, all cells in the interaction matrix not covered by pcHi-C baits were removed. Second, bins with poor coverage (with less than 11 counts chromosome-wide) were treated as non-captured.

The filtered interaction matrices were then normalized by distinguishing between *bait-bait* and *bait-non-bait* interactions. In brief, an intersection between sites was normalized by the least interacting captured site of the two intersecting bins:

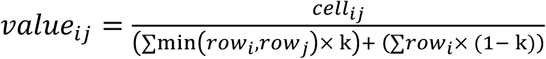

where *k* is set to 1 for bait-bait interactions and to 0 in bait-non-bait interactions.

#### Generation of 3D ensemble structures of enhancer hubs

Related to Figure 5b,c and **Supplementary Figure 5b,c.** Normalized interaction matrices were modelled using TADdyn, a molecular dynamic-based protocol implemented in the TADbit software ^52^. Similarly to TADbit, TADdyn generates models using a restraint-based approach, in which experimental frequencies of interaction were transformed into a set of spatial restraints, as previously described ^53^, but suited for sparse datasets such as pcHi-C. The underlying chromatin fiber was represented by a bead-spring polymer model ^54-58^, where the size (σ) of each bead was defined by the relationship 0.01 nm/bp assuming the canonical 30 nm fiber ^59,60^, as implemented in TADbit. The system scoring function (*H)* was defined as:

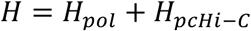

where *H*_pol_ is the contribution of the polymer and H_pcHi-C_ is defined by a set of distance restrains proportional to the experimental frequencies of interaction. *H*_pol_ comprises two potentials that prevent polymer crossings when the beads interact:

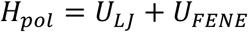

U_LJ_ potential is a purely repulsive Lennard-Jones potential used for the excluded volume interaction defined as:

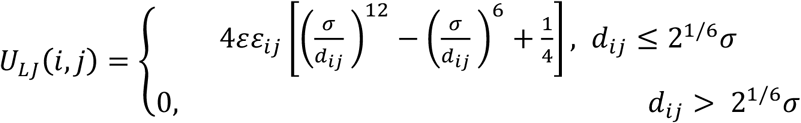

where ε = *k*_*B*_*T* is energy unit of the system, *k*_*B*_ is the Boltzmann constant, *T* = 1.0 (in rescaled internal units) is the temperature, ε*_ij_* is a parameter equal to 10 for connected particles (|*i − j|*= 1), and 1 otherwise, and *d*_*ij*_ is the distance between particles *i* and *j*. The UFENE (Finitely Extensible Nonlinear Elastic) potential used to define the fibre connectivity defined as:

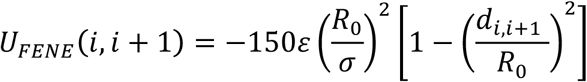

This ensure that consecutive particles on the chain are connected by an elastic energy which allows a maximum bond extension of *R*_0_ = 1.5σ = *75nm*. H_pcHi-C_ is defined by a set of distance restrains (imposed using harmonics) proportional to the experimental frequencies of interaction:

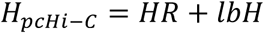

The harmonic restraint (HR) between two particles i and j of the system at spatial distance d*_ij_*, is defined as:

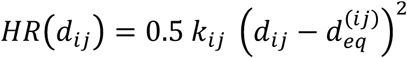

where *k*_*ij*_ is the strength of the restraint, and 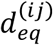 is the equilibrium distance as calculated from the normalized pcHi-C interaction matrix. Here, this type of harmonics are used for (i) imposed proper Harmonic restraint (*HR*), where particles i and j are forced to stay at a distance close to equilibrium; and (ii) impose LowerBoundHarmonic restraint (*lbH*), where particles i and j are forced to stay at a mutual distance larger than equilibrium. The former HR is defined by large strength and short equilibrium distance to bring together in space each pair of particles with high normalized pcHi-C interaction value; conversely, the latter is defined by large strength and large equilibrium distance to keep far apart each pair of particles with low normalized pcHi-C interaction value.

Similarly to TADbit, a grid search approach was used to identify empirically the three optimal parameters to be used: (i) maximal distance between two non-interacting particles (*maxdist)* defining the scale of the obtained models; (ii) a lower-bound cutoff to define particles that do not interact frequently (*lowfreq)*; and (iii) an upper-bound cutoff to define particles that do interact frequently (*upfreq)* ^52,53,61^.

Finally, during the molecular dynamic simulation optimization step the initial polymeric (random walk) configurations were placed in a cubic simulation box of size *10*1σ. The dynamics of the system were described using the underdumped Langevin equation, as implemented in LAMMPs ^62^ with standard parameters ^54^. The *HR* and *lbH* restraints based on the normalized pcHi-C interaction matrix were applied using a modified version of the collective variables Colvar plug-in ^63^.

A total of 500 models were generated for each genomic region and cell type (islets and total B lymphocytes). The contact map generated from the ensemble of models was highly correlated with the input pcHi-C normalized interaction matrices (see **Supplementary Table 11**).Each enhancer hub ensemble was clustered based on structural similarity as implemented in TADbit ^52^, and only the models from the most populated cluster were selected for further analysis.

#### Structural analysis of 3D models

Related to **Supplementary Figure 5.** A set of descriptive measures were calculated to analyze the structural properties of each particle in the most populated cluster of the model ensemble using TADbit ^52^: (i) consistency, which indicates for each particle the percentage of models that superimpose a given particle within a cut-off of 200 nm; (ii) accessibility, which is a measure of how accessible a particle is to an object of radius of 50 nm; and (iii) interactions, which counts the number of particles within a given spatial distance (2 times particle size) from a specified particle. Furthermore, TADbit tools was used to calculate the mean distances particles containing genomic regions of interest in the model. Using the mean distances, a promoter-enhancer interaction network was built as a weighted undirected graph, in which each node represents either a promoter or an enhancer in a specific locus and each edge is weighted by its mean distances values. The promoter-enhancer interaction network was decomposed into communities using MCODE clustering algorithm as implemented in clusterMaker2 ^64^ and visualized with Cytoscape 3.5 ^65^.

#### Spatial distribution of chromatin marks in 3D models

Related to Figure 5d and **Supplementary Figure 5d.** The 3D spatial distribution of ChIP-seq data from human islets was analyzed as follows: for a given central viewpoint, an initial sphere with a radius of 200 nm was constructed from the viewpoint. Then, a set of spherical shells, that occupied a volume equal the initial sphere, were added. Each particle neighboring the viewpoint was assigned to a spherical shell based on their relative distance to the viewpoint. ChIP-Seq data from MACS2 ^66^ was then binned at 5 kb resolution and the mean score value of their coverage score was calculated. The odd ratio was calculated for each shell using a Fisher’s exact test for 2 × 2 contingency tables comparing bins with and without ChIP-Seq signal. A total of 13 different ChIP-seq datasets of human pancreatic islets (CTCF, FOXA2, H3K27me3, H3K27ac, H3K36me3, H3K4me1, H3K4me3, MAFB, NKX2.2, NKX6.1, PDX1, Mediator and cohesin) were analysed and the corresponding circular 2D plot was generated.

#### **SNP enrichment analyses.** Related to Figure 6 and **Supplementary Figure 6.**

The Variant Set Enrichment (VSE) R package ^67^ was used to compute the enrichment of T2D and FG-associated variants in regulatory annotations. VSE creates associated variant sets using lead SNPs and variants in the same haplotype block, and then determines whether overlaps with genomic annotations are enriched relative to null distributions from variant sets that are matched for size and haplotype structure ^68^. As input, we used lead SNPs from the 109 loci listed in **Supplementary Table 4**. Breast cancer lead SNPs were also downloaded from NHGRI-EBI GWAS Catalogue on 27/09/2016, and used as a control. Variants in close linkage disequilibrium (r^2^>0.8) with the lead SNPs were calculated and linkage disequilibrium blocks were built using 1000 Genome Project, Phase III, Oct 2014, Hg19 data and http://raggr.usc.eduwebtool. The sets of lead SNPs and their linked SNPs in high linkage disequilibrium constitute the Associated Variant Set (AVS). VSE ensures that each SNP is only represented once, even when they are in high linkage disequilibrium with more than one lead SNPs. To account for the size and structure of the AVS, a null distribution or Matched Random Variant Set (MRVS) was built from 1000 Genome Project Phase III, based on 500 random permutations of the AVS. The MRVS had an identical total number of loci as the AVS and it was matched in size and structure to the original AVS. The enrichment of the intersections between AVS and the provided genomic features was computed by tallying the number of independent SNPs that overlap with the functional annotation, and comparing it to the overlap of the null distribution (MRVS). VSE enrichment score was defined as the number of standard deviations that the overlapping tally deviates from the null overlapping tally median. The exact p value was calculated by VSE by fitting a density function into the null distribution derived from the MRVS. Significant enrichments or depletions were considered when the Bonferroni adjusted p-value was < 0.01. Human islet regulatory elements used in Supplementary Figure 6 can be obtained from **Extended Data 1.1**. Interacting regions shown in Figure 6 are defined as HindIII fragments in the human genome in either end of high-confidence pcHi-C interactions.

#### **QQ plots.** Related to **Figure 6.**

For genomic inflation estimates of the human islet regulatory annotations we used quantile-quantile (Q-Q) plots, which display the expected –log_10_ p-values under the null hypothesis in the x-axis and observed –log_10_ p-values in the y-axis. We estimated the λ coefficients, a measure of genomic inflation that corresponds to the observed median χ2 test statistic divided by the expected median χ2 test statistic under the null hypothesis. Uniform null distributions were highlighted with a red line. Q-Q plots were generated using summary statistics from^35^, only considering variants with MAF ≥ 5% and p-value ≥ 5×10^−8^. The total number of variants considered was 6,647,137. To assess variants overlapping interacting regions we showed p-value distributions in Q-Q representations and λ measures for variants located in (i) high-confidence pcHi-C interactions (baits and promoter interacting regions), (ii) non-interacting open-chromatin regulatory elements and (iii) the rest of variants that did not overlap either of the previous categories. To assess class I enhancers inside hubs we generated Q-Q plots and λ measures for variants overlapping (i) class I enhancers inside hubs, (ii) open chromatin regulatory elements outside hubs and (iii) the rest of variants that did not overlap either of the previous categories.

#### GWAS meta-analysis of Insulinogenic index from oral glucose tolerance test (OGTT) measures

Related to **Figure 6.** 7,807 individuals from four population studies were included in these analyses: the Inter99 study (ClinicalTrials.gov ID-no: NCT00289237) which is a population-based non-pharmacological intervention study for ischemic heart disease (n=5,305) ^69^, the Health2008 cohort (n=605) ^70^, the 1936 Birth Cohort (n=709) ^71^ and the ADDITION-Pro cohort (n=1,188) ^72^. All study participants gave informed consent and the studies were approved by the appropriate Ethical Committees and were performed in accordance with the scientific principles of the Helsinki Declaration II.

In the four cohorts, glucose-stimulated insulin secretion was evaluated by measurement of plasma glucose and serum insulin at 0, 30 and 120 minutes during a 75 g oral glucose tolerance test (OGTT). Insulinogenic index was calculated based on these as Insulinogenic index = (s-insulin at 30 minutes [pmol/l] − fasting s-insulin [pmol/l]) / p-glucose at 30 minutes (mmol/l). In these analyses, individuals with known diabetes were excluded.

Two sample sets (Inter99 and Health2008) were then genotyped by Illumina OmniExpress array while the two other cohorts were genotyped by Illumina CoreExome array. Genotypes were called by Illumina GenCall algorithm. Genotype data were filtered for variants with call rate <98% and Hardy-Weinberg equilibrium P<10^−5^. Samples were excluded if they were ethnic outliers, had mismatch between genetic and phenotypic sex or had a call rate <95%.

Genotype data from each cohort was imputed to the Haplotype Reference Consortium (HRC) reference panel v1.1 ^73^ at the Michigan Imputation Server (https://imputationserver.sph.umich.edu/index.html#!pages/home) using Minimac3 after phasing genotypes into haplotypes applying Eagle2 ^74^. Posvt-imputation SNP filtering included excluding variants with MAF <0.01 and info score <0.70. In each cohort, association analysis was performed by applying a linear regression model including age and sex as covariates via SNPTEST ^75^. The phenotype was rank-normalized within each cohort before analysis. A fixed-effects meta-analysis implemented in the R package *meta* ^76^ was finally performed.

#### GWAS meta-analysis of homeostasis model assessment β cell function (HOMA-B) and insulin resistance (HOMA-IR)

Related to **Figure 6.** To perform GWAS meta-analysis for HOMA-B and HOMA-IR, GWAS summary statistics were approximated using a recently developed approach, GWIS ^77^. We obtained summary statistics from the latest sex-specific and sex-differentiated GWAS meta-analysis of fasting glycemia (FG) and fasting insulin (FI) performed by the Meta-Analysis of Glucose and Insulin-related traits Consortium (MAGIC) in up to 88,320 and 64,090 individuals (that are comprised in 40 and 33 studies), respectively. To obtain summary statistics data that were appropriate for LD-score regression ^78^ analysis, individuals that were genotyped on the Metabochip array were excluded. FI was measured in pmol/l and natural log transformed, and FG was measured in mmol/l with a cut-off at 7 mmol/l, which were used to generate sex-specific summary statistics for meta-analysis of untransformed FG (up to 67,506 men and 73,089 women) and ln-transformed FI (up to 47,806 men and 50,404 women), without BMI-adjustment. The standard HOMA formulas require untransformed FG/FI measures and use mU/l units for FI. Therefore, HOMA formulae was adapted to compute a GWAS summary statistics for ln-transformed HOMA-B and HOMA-IR given the summary statistics for FG and ln(FI).

#### Heritability estimates

Related to Figure 6 and **Supplementary Figure 7.** To estimate the polygenic contribution of different genomic annotations to GWAS-based heritability of T2D and related traits we applied the stratified LD Score regression method ^78,79^. This method partitions heritability from GWAS summary statistics while accounting for the linkage disequilibrium (LD) structure between markers. The method leverages the relationship between LD structure and association test statistics to estimate the average per-SNP contribution to heritability (τ_c_ coefficient) for a functional category *C*. Briefly, following the relationship in equation:

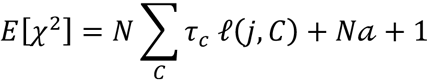

where *N* is the sample size, *C* indexes categories, ℓ(*j, C*) is the stratified LD Score of *SNP*_*j*_ with respect to functional category *C*, and *a* measures the contribution of confounding biases. The per-SNP heritability contribution to category *C* (τ_c_) for either a quantitative or case-control trait is estimated with the previous equation via multiple regression of association test statistics (χ2) for *SNP*_*j*_ against ℓ*(j, C*). Finally, statistical significance is calculated by testing whether per-SNP heritability is greater in the category *C* than out of the category. The Z-score of the τ_c_ coefficient measured the statistical significance of the contribution of category *C* to SNP heritability after controlling for the effects of a full baseline model of 53 genomic annotations defined by ^79^. Further details are explained in^79^.

We obtained GWAS summary statistics for T2D ^35^, Insulinogenic index (see **GWAS meta-analysis of Insulinogenic index from oral glucose tolerance test (OGTT) measures**), HOMA-B and HOMA-IR (see **GWAS meta-analysis of homeostasis model assessment β cell function (HOMA-B) and insulin resistance (HOMA-IR)**), and Acute Insulin Response (AIR) based on intravenous glucose tolerance tests (IVGTT) ^80^. As a control trait, we used summary statistics from a well-powered meta-analysis for Attention-Deficit/Hiperactivity Disorder (ADHD)^81^. To partition SNP-heritability based on GWAS summary statistics across islet regulatory annotations, we followed precisely the procedure described by ^79^. We used the stratified LD Scores calculated from the European-ancestry samples from the 1000 Genome Project, Phase III ^82^ and in the regression analysis we only considered the association statistics of ∼1M HapMap3 SNPs (excluding the MHC region and including SNPs with MAF ≥ 5%).

We tested whether any of the following human islet regulatory annotations contributes significantly to the SNP heritability of islet related traits: (a) open chromatin regions, (b) regions with strong CTCF binding (I and II), (c) inactive enhancers, (d) active promoters, (e) class I enhancers, (f) class II enhancers, (g) class III enhancers, (h) class I enhancers in enhancer hubs, (i) highly-occupied C3 clustered enhancers ^9^, (j) all C3 clustered enhancers ^9^(k) enhancer clusters (with intervening space) ^9^, (j) stretch enhancers ^51^, strong enhancer states^83^ and (k) super-enhancers as defined in this study.

We also included three control annotation sets. Our first control was Central Nervous System (CNS) functional category – a collection of cell-type specific annotations defined by ^79^. A second control annotation was generated by creating a set of random non-open chromatin regions matching enhancer length (average length = ∼800 bp). We excluded ENCODE blacklisted regions (wgEncodeDacMapabilityConsensusExcludable.bed.gz and wgEncodeDukeMapabilityRegionsExcludable.bed.gz) and regions with average ENCODE mappability scores (wgEncodeCrgMapabilityAlign100mer.bigWig) below the average score across all enhancers (∼0.997). Finally, we built control pseudo-hubs as the third control annotation. Pseudo-hubs were anchored in non-hub baits that otherwise showed high-confidence interactions in islet pcHi-C maps. For every real hub, we selected non-hub baits and associated genomic regions that were intended to match the enhancers assigned to the real hubs (referred to in Figure 6 as pseudo-enhancer). To associate control genomic regions with non-hub baits we selected 800 bp windows within the same islet TAD that had the closest possible distance to the anchor as enhancer-bait distance from matching real enhancer hubs. Only TAD-like domains that contained at least one active annotation in the islet regulome were selected. We ensured that the 800 bp windows chosen to match interacting hub enhancers (i) did not overlap active enhancers, active promoters or pc-HIC baits, (ii) did not overlap blacklisted regions and (iii) achieved a regional mappability score above the average score for active enhancers. To obtain a control annotation for hub class I enhancers, we only considered the fraction of 800 bp-windows that were selected based on class I enhancer-bait distances from the matching real hubs.

We added these islet and control annotations to the full baseline model of 53 functional annotations ^79^, one at a time, and we calculated the magnitude and statistical significance of the per-SNP contribution to heritability of each annotation. We provided the per-SNP heritability τ_c_ coefficient for each particular regulatory annotation and phenotype. To facilitate comparability across traits and annotations, we normalized the τ_c_ estimates by dividing them by the observed scale LD Score heritability for each phenotype, and we multiplied by 10^7^. To correct for multiple testing, we generated τ_c_ *q*-values (FDR-adjusted p-values calculated from the Z-scores of the τ_c_ coefficients) with the *qvalue* R package (Alan Dabney and John Storey, Department of Biostatistics, University of Washington) over 17 functional categories and 6 traits. FDR significance threshold was set at 0.05.

#### Polygenic risk scores (PRSs)

Related to **Figure 6.** We created Polygenic risk scores (PRSs) based on T2D GWAS summary statistics from ^35^ (*base dataset*). UK Biobank individuals ^84^ were used as the *target datasets*, which comprised testing and validation datasets.

To select markers used for PRS we first considered all genetic markers that were used as input for phasing and genotype imputation by UK Biobank investigators, and filtered for variants with MAF ≥ 5% and imputation quality score > 0.8. We then reconciled the base and target datasets by looking at the variant overlap between summary statistics and the imputed UK Biobank data, discarding variants showing allele inconsistency between both datasets. We also removed those located in the MHC region, resulting in a final collection of 5,352,387 variants.

Concerning UK Biobank samples, we first applied the following exclusions to the entire cohort: (i) individuals with excess of relatives (> 10 third-degree relatives according to the UK Biobank dataset), (ii) individuals without gender information, (iii) individuals with ICD10 codes E10 (insulin-dependent diabetes mellitus), E13 (other specified diabetes mellitus) and E14 (unspecified diabetes mellitus), (iv) individuals without body mass index (BMI) information, and (v) in pairs of individuals with third-degree relatedness (as provided by UK Biobank investigators) we excluded subjects with the highest missing rate for a set of high-quality markers. T2D cases were defined as those with E11 ICD10 diagnosis code. The sample size of the final set of qualifying UK Biobank individuals was 377,981 controls and 15,764 cases.

To validate the T2D case-control definition based on ICD10 codes, we estimated the association statistics for well-established T2D loci. We used 15,764 T2D cases (accounting for information about age of diagnosis) and 184,030 controls that were filtered to avoid any contamination of pre-diabetic individuals (no family history of diabetes mellitus and age at recruitment ≥ 55, consistent with the average age at onset of T2D reported in Caucasian populations ^85^). The association statistics under an additive model for 74 known T2D associated variants were estimated by applying logistic regression in PLINK v1.9 ^36^, and adjusting for 7 principal components (estimated by UK Biobank investigators), BMI, age at recruitment, batch information and sex. From these, only five signals, from which four were discovered in East Asian populations, do not show consistent direction of effect with the reference study (**Supplementary Table 12**).

The entire dataset of 377,981 controls and 15,764 cases was divided in test and validation datasets. For the test dataset, we included only control subjects with age at recruitment ≥ 55 years and no family history of diabetes mellitus. The final sample size of the UK Biobank test dataset was 6,305 T2D cases and 73,922 controls. The remaining UK Biobank samples were used as a validation dataset, comprising 9,459 T2D cases and 226,777 controls. Control individuals in this validation dataset were included regardless of their age or family history.

PRS models were calculated using the PRsice software ^86^ with default settings and the following clumping parameters *(--clump-r2 0.6 --clump-p 0.01*). We included 11 covariates in the analysis: the 7 principal components provided by UK Biobank investigators as well as BMI, age at recruitment, batch information, and sex. PRS models were built using the T2D GWAS summary statistics from ^35^ as the base dataset and the abovementioned subset of the UK Biobank as the test dataset (6,305 T2D cases and 73,922 controls).

To examine islet enhancer hubs, we calculated PRS using variants overlapping all hub pcHi-C baits and enhancers. This generated a PRS model of 179 variants. In parallel, we generated a genome-wide PRS model based on the entire set of common genetic variants shared by the base and the validation dataset (∼5M variants) that finally comprised 1,152 variants. As an additional control, we used genetic variants overlapping pseudo-enhancer hubs. We generated pseudo-enhancer hubs by shuffling all hub pcHi-C baits and their assigned enhancer fragments across the genome. We excluded pseudo-enhancer hubs that mapped to TADs containing real hubs, or that crossed TAD boundaries. Pseudo-interacting regions that mapped to blacklisted regions or had low mappability were excluded. We then built a PRS using variants overlapping pseudo-baits and pseudo-enhancers. We generated 100 sets of pseudo-baits and pseudo-enhancers and the corresponding PRS models (median of 280 variants).

We applied the resulting PRS models to the UK Biobank validation dataset (9,459 T2D cases and 226,777 controls) following the equation below provided by the PRSice authors ^86^:

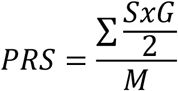

where *S* corresponds to the summary statistic of the effective allele, *G* is the number of the effective allele and *M* is the number of SNPs.

To assess the predictive power of genome-wide, islet hub and pseudo-hub PRS, we examined the T2D frequency in the UK Biobank validation dataset across 40 bins, each one containing 2.5% of individuals ranked by the PRS score. For analyses that considered PRS as a function of BMI and age of onset, we focused on all 226,777 control individuals and a subset of 6,127 T2D cases with age of diagnosis > 20. For each of the three PRS models (genome-wide, islet hub and 100 sets of pseudo-hubs), we calculated the ratio of T2D frequency in the highest and lowest PRS bins, each one containing 2.5% of the entire population. We also used cox proportional hazard models, with age of onset (age at recruitment for controls) as the underlying time-scale, to analyse the effect of highest and lowest PRS values on the risk of T2D for the genome-wide, the islet-hub and 100 sets of pseudo-hubs PRS models. The effect of the top 2.5% PRS bin was calculated using as reference all the other PRS bins (1-39), and the same approach was used to analyze the protective effect on T2D for the bottom 2.5% PRS bin. In this analysis, the risk was presented as hazard ratios (HR) of extreme PRS group with the 95% confidence interval (CI). We adjusted the analysis for sex, BMI, 7 principal components provided by UK Biobank investigators and batch information. Kaplan-Meier graphs of cumulative incidence of T2D are shown for the subset of individuals with highest (bin 40), intermediate (bin 2-39) and lowest (bin 1) PRS scores for the genome-wide and the islet-hub PRS model.

For the islet hub PRS model, and the PRS of 100 sets of pseudo-hubs, we stratified this analysis in three categories of BMI (< 30, 30 − <35 and ≥ 35), and in three categories of age of diagnosis for T2D (<50, 50 − < 60 and ≥ 60). The age of controls was censored at the time of recruitment. We also used cox proportional hazard models to show the HR for the top and bottom PRS bins according to the genome-wide and islet hub PRS models across three BMI categories (< 30, 30 − <35 and ≥ 35) and two categories of age of diagnosis (<60 and ≥ 60). Results are illustrated in forest plots with effect sizes and 95% CI. We further examined the predictive power of the PRS models by calculating the area-under-the curve (AUC) statistic. We addressed the performance of the PRS models in two ranges of BMI (< 35 and ≥ 35) and in two ranges of age of diagnosis for T2D (<60 and ≥ 60). Finally, we calculated the odds ratio for T2D in individuals with highest PRS scores (top 2.5% bin) compared to the remainder UK Biobank individuals of the validation dataset in different age and BMI categories. We assessed this with a logistic regression model adjusted for the first seven principal components of ancestry, sex, age, BMI and batch information. We used z-scores to test for statistically significant differences in either the ratio of T2D frequencies or the ORs between the islet hub and the genome-wide PRS, and the distribution of 100 PRS sets of pseudo-hubs.

## References

1. Andersson, R. et al. An atlas of active enhancers across human cell types and tissues. Nature 507, 455–461 (2014).

2. Dunham, I. et al. An integrated encyclopedia of DNA elements in the human genome. Nature 489, 57–74 (2012).

3. Roadmap Epigenomics, C. et al. Integrative analysis of 111 reference human epigenomes. Nature 518, 317–30 (2015).

4. Chatterjee, S., Khunti, K. & Davies, M.J. Type 2 diabetes. Lancet 389, 2239–2251 (2017).

5. Flannick, J. & Florez, J.C. Type 2 diabetes: genetic data sharing to advance complex disease research. Nat Rev Genet 17, 535–49 (2016).

6. Fuchsberger, C. et al. The genetic architecture of type 2 diabetes. Nature 536, 41–47 (2016).

7. Parker, S.C. et al. Chromatin stretch enhancer states drive cell-specific gene regulation and harbor human disease risk variants. Proc Natl Acad Sci U S A 110, 17921–6 (2013).

8. Pasquali, L. et al. Pancreatic islet enhancer clusters enriched in type 2 diabetes risk-associated variants. Nat Genet 46, 136–143 (2014).

9. Thurner, M. et al. Integration of human pancreatic islet genomic data refines regulatory mechanisms at Type 2 Diabetes susceptibility loci. Elife 7(2018).

10. Fadista, J. et al. Global genomic and transcriptomic analysis of human pancreatic islets reveals novel genes influencing glucose metabolism. Proc Natl Acad Sci U S A 111, 13924–9 (2014).

11. van de Bunt, M. et al. Transcript Expression Data from Human Islets Links Regulatory Signals from Genome-Wide Association Studies for Type 2 Diabetes and Glycemic Traits to Their Downstream Effectors. PLoS Genet 11, e1005694 (2015).

12. Whyte, W.A. et al. Master transcription factors and mediator establish superenhancers at key cell identity genes. Cell 153, 307–19 (2013).

13. Gaulton, K.J. et al. A map of open chromatin in human pancreatic islets. Nat Genet 42, 255–9 (2010).

14. Farh, K.K. et al. Genetic and epigenetic fine mapping of causal autoimmune disease variants. Nature 518, 337–43 (2015).

15. Vahedi, G. et al. Super-enhancers delineate disease-associated regulatory nodes in T cells. Nature 520, 558–62 (2015).

16. Hnisz, D. et al. Super-enhancers in the control of cell identity and disease. Cell 155, 934–47 (2013).

17. Cohen, A.J. et al. Hotspots of aberrant enhancer activity punctuate the colorectal cancer epigenome. Nat Commun 8, 14400 (2017).

18. Lee, S. et al. Optimal unified approach for rare-variant association testing with application to small-sample case-control whole-exome sequencing studies. Am J Hum Genet 91, 224–37 (2012).

19. Mansour, M.R. et al. Oncogene regulation. An oncogenic super-enhancer formed through somatic mutation of a noncoding intergenic element. Science 346, 1373–7 (2014).

20. Javierre, B.M. et al. Lineage-Specific Genome Architecture Links Enhancers and Non-coding Disease Variants to Target Gene Promoters. Cell 167, 1369–1384 e19 (2016).

21. Cairns, J. et al. CHiCAGO: robust detection of DNA looping interactions in Capture Hi-C data. Genome Biol 17, 127 (2016).

22. Scharfmann, R. et al. Development of a conditionally immortalized human pancreatic beta cell line. J Clin Invest 124, 2087–98 (2014).

23. Schofield, E.C. et al. CHiCP: a web-based tool for the integrative and interactive visualization of promoter capture Hi-C datasets. Bioinformatics 32, 2511–3 (2016).

24. Mularoni, L., Ramos-Rodriguez, M. & Pasquali, L. The Pancreatic Islet Regulome Browser. Front Genet 8, 13 (2017).

25. Dixon, J.R. et al. Topological domains in mammalian genomes identified by analysis of chromatin interactions. Nature 485, 376–80 (2012).

26. Nora, E.P. et al. Spatial partitioning of the regulatory landscape of the X-inactivation centre. Nature 485, 381–5 (2012).

27. Benazra, M. et al. A human beta cell line with drug inducible excision of immortalizing transgenes. Mol Metab 4, 916–25 (2015).

28. Scott, R.A. et al. An Expanded Genome-Wide Association Study of Type 2 Diabetes in Europeans. Diabetes 66, 2888–2902 (2017).

29. Wood, A.R. et al. A Genome-Wide Association Study of IVGTT-Based Measures of First-Phase Insulin Secretion Refines the Underlying Physiology of Type 2 Diabetes Variants. Diabetes 66, 2296–2309 (2017).

30. Gaulton, K.J. et al. Genetic fine mapping and genomic annotation defines causal mechanisms at type 2 diabetes susceptibility loci. Nat Genet 47, 1415–25 (2015).

31. Xia, Q. et al. The type 2 diabetes presumed causal variant within TCF7L2 resides in an element that controls the expression of ACSL5. Diabetologia 59, 2360–2368 (2016).

32. Nobrega, M.A. TCF7L2 and glucose metabolism: time to look beyond the pancreas. Diabetes 62, 706–8 (2013).

33. Barrett, J.C. et al. Genome-wide association study and meta-analysis find that over 40 loci affect risk of type 1 diabetes. Nat Genet 41, 703–7 (2009).

34. Morris, A.P. et al. Large-scale association analysis provides insights into the genetic architecture and pathophysiology of type 2 diabetes. Nat Genet 44, 981–90 (2012).

35. Senee, V. et al. Mutations in GLIS3 are responsible for a rare syndrome with neonatal diabetes mellitus and congenital hypothyroidism. Nat Genet 38, 682–7 (2006).

36. Ait-Lounis, A. et al. The transcription factor Rfx3 regulates beta-cell differentiation, function, and glucokinase expression. Diabetes 59, 1674–85 (2010).

37. Bau, D. et al. The three-dimensional folding of the alpha-globin gene domain reveals formation of chromatin globules. Nat Struct Mol Biol 18, 107–14 (2011).

38. Serra, F. et al. Automatic analysis and 3D-modelling of Hi-C data using TADbit reveals structural features of the fly chromatin colors. PLoS Comput Biol 13, e1005665 (2017).

39. Varshney, A. et al. Genetic regulatory signatures underlying islet gene expression and type 2 diabetes. Proc Natl Acad Sci U S A 114, 2301–2306 (2017).

40. Wood, A.R. et al. Defining the role of common variation in the genomic and biological architecture of adult human height. Nat Genet 46, 1173–86 (2014).

41. Boyle, E.A., Li, Y.I. & Pritchard, J.K. An Expanded View of Complex Traits: From Polygenic to Omnigenic. Cell 169, 1177–1186 (2017).

42. Khera, M.C., K.; Aragam, C.A.; Emdin, D.; Klarin, M.; Haas, C.; Roselli, P.; Natarajan, S.; Kathiresan. Genome-wide polygenic score to identify a monogenic risk-equivalent for coronary disease. BioRxiv (2017).

43. Finucane, H.K. et al. Partitioning heritability by functional annotation using genome-wide association summary statistics. Nat Genet 47, 1228–35 (2015).

44. Gjesing, A.P. et al. Genetic and phenotypic correlations between surrogate measures of insulin release obtained from OGTT data. Diabetologia 58, 1006–12 (2015).

45. Anubha Mahajan, D.T., Matthias Thurner, Neil R Robertson, Jason M Torres, N William Rayner, Valgerdur Steinthorsdottir, Robert A Scott, Niels Grarup, James P Cook, Ellen M Schmidt, Matthias Wuttke, Chloé Sarnowski, Reedik Mägi, Jana Nano, Christian Gieger, Stella Trompet, Cécile Lecoeur, Michael Preuss, Bram Peter Prins, Xiuqing Guo, Lawrence F Bielak, DIAMANTE Consortium, Amanda J Bennett, Jette Bork-Jensen, Chad M Brummett, Mickaël Canouil, Kai-Uwe Eckardt, Krista Fischer, Sharon LR Kardia, Florian Kronenberg, Kristi Läll, Ching-Ti Liu, Adam E Locke, Jian’an Luan, Ioanna Ntalla, Vibe Nylander, Sebastian Schönherr, Claudia Schurmann, Loïc Yengo, Erwin P Bottinger, Ivan Brandslund, Cramer Christensen, George Dedoussis, Jose C Florez, Ian Ford, Oscar H Franco, Timothy M Frayling, Vilmantas Giedraitis, Sophie Hackinger, Andrew T Hattersley, Christian Herder, M Arfan Ikram, Martin Ingelsson, Marit E Jørgensen, Torben Jørgensen, Jennifer Kriebel, Johanna Kuusisto, Symen Ligthart, Cecilia M Lindgren, Allan Linneberg, Valeriya Lyssenko, Vasiliki Mamakou, Thomas Meitinger, Karen L Mohlke, Andrew D Morris, Girish Nadkarni, James S Pankow, Annette Peters, Naveed Sattar, Alena Stančáková, Konstantin Strauch, Kent D Taylor, Barbara Thorand, Gudmar Thorleifsson, Unnur Thorsteinsdottir, Jaakko Tuomilehto, Daniel R Witte, Josée Dupuis, Patricia A Peyser, Eleftheria Zeggini, Ruth J F Loos, Philippe Froguel, Erik Ingelsson, Lars Lind, Leif Groop, Markku Laakso, Francis S Collins, J Wouter Jukema, Colin N A Palmer, Harald Grallert, Andres Metspalu, Abbas Dehghan, Anna Köttgen, Goncalo Abecasis, James B Meigs, Jerome I Rotter, Jonathan Marchini, Oluf Pedersen, Torben Hansen, Claudia Langenberg, Nicholas J Wareham, Kari Stefansson, Anna L Gloyn, Andrew P Morris, Michael Boehnke, View ORCID ProfileMark I McCarthy. Fine-mapping of an expanded set of type 2 diabetes loci to single-variant resolution using high-density imputation and islet-specific epigenome maps. BioRxiv (2018).

46. Bonas-Guarch, S. et al. Re-analysis of public genetic data reveals a rare Xchromosomal variant associated with type 2 diabetes. Nat Commun 9, 321 (2018).

47. Sudlow, C. et al. UK biobank: an open access resource for identifying the causes of a wide range of complex diseases of middle and old age. PLoS Med 12, e1001779 (2015).

48. Bycroft, C.F., D.; Petkova, G.; Band, L.T.; Elliott, K.; Sharp, A.; Motyer, D.; Vukcevic, O.; Delaneau, J.; O’Connell, A.; Cortes, S.; Welsh, G.; McVean, S.; Leslie, P.; Donnelly, J.; Marchini. Genome-Wide Genetic Data on ∼ 500,000 UK Biobank Participants. BioRxiv (2017).

49. Osterwalder, M. et al. Enhancer redundancy provides phenotypic robustness in mammalian development. Nature 554, 239–243 (2018).

50. Montavon, T. et al. A regulatory archipelago controls Hox genes transcription in digits. Cell 147, 1132–45 (2011).

51. Raich, N. et al. Autonomous developmental control of human embryonic globin gene switching in transgenic mice. Science 250, 1147–9 (1990).

52. Patrinos, G.P. et al. Multiple interactions between regulatory regions are required to stabilize an active chromatin hub. Genes Dev 18, 1495–509 (2004).

53. Akalin, A. et al. Transcriptional features of genomic regulatory blocks. Genome Biol 10, R38 (2009).

54. Harmston, N. et al. Topologically associating domains are ancient features that coincide with Metazoan clusters of extreme noncoding conservation. Nat Commun 8, 441 (2017).

55. Schmitt, A.D. et al. A Compendium of Chromatin Contact Maps Reveals Spatially Active Regions in the Human Genome. Cell Rep 17, 2042–2059 (2016).

56. Ahlqvist, E. et al. Novel subgroups of adult-onset diabetes and their association with outcomes: a data-driven cluster analysis of six variables. Lancet Diabetes Endocrinol (2018).

57. DeFronzo, R.A. et al. Type 2 diabetes mellitus. Nat Rev Dis Primers 1, 15019 (2015).

58. Inouye, M.A.G.; Nelson, C. P.; Wood, A. M.; Sweeting, M. J.; Dudbridge, F.; Lai, F. Y.; Kaptoge, S.; Brozynska, M.; Wang, T.; Ye, S.; Webb, T. R.; Rutter, M. K.; Tzoulaki, I.; Patel, R. S.; Loos, R. JL.; Keavney, B.; Hemingway, H.; Thompson, J.; Watkins, H.; Deloukas, P.; Di Angelantonio, E.; Butterworth, A. S.; Danesh, J.; Samani, N. J. Genomic risk prediction of coronary artery disease in nearly 500,000 adults: implications for early screening and primary prevention. BioRxiv (2018).

## References for Supplementary Material

1. Ricordi, C., Lacy, P.E., Finke, E.H., Olack, B.J. & Scharp, D.W. Automated method for isolation of human pancreatic islets. Diabetes 37, 413–20 (1988).

2. Melzi, R. et al. Role of CCL2/MCP-1 in islet transplantation. Cell Transplant 19, 1031–46 (2010).

3. Kerr-Conte, J. et al. Upgrading pretransplant human islet culture technology requires human serum combined with media renewal. Transplantation 89, 1154–60 (2010).

4. Bucher, P. et al. Assessment of a novel two-component enzyme preparation for human islet isolation and transplantation. Transplantation 79, 91–7 (2005).

5. Nagano, T. et al. Comparison of Hi-C results using in-solution versus in-nucleus ligation. Genome Biol 16, 175 (2015).

6. Javierre, B.M. et al. Lineage-Specific Genome Architecture Links Enhancers and Non-coding Disease Variants to Target Gene Promoters. Cell 167, 1369–1384 e19 (2016).

7. Wingett, S. et al. HiCUP: pipeline for mapping and processing Hi-C data. F1000Res 4, 1310 (2015).

8. Cairns, J. et al. CHiCAGO: robust detection of DNA looping interactions in Capture Hi-C data. Genome Biol 17, 127 (2016).

9. Pasquali, L. et al. Pancreatic islet enhancer clusters enriched in type 2 diabetes risk-associated variants. Nat Genet 46, 136–143 (2014).

10. Kagey, M.H. et al. Mediator and cohesin connect gene expression and chromatin architecture. Nature 467, 430–5 (2010).

11. Buenrostro, J.D., Giresi, P.G., Zaba, L.C., Chang, H.Y. & Greenleaf, W.J. Transposition of native chromatin for fast and sensitive epigenomic profiling of open chromatin, DNA-binding proteins and nucleosome position. Nat Methods 10, 1213–8 (2013).

12. Martin, M. Cutadapt removes adapter sequences from high-throughput sequencing reads. 2011 17(2011).

13. Langmead, B. & Salzberg, S.L. Fast gapped-read alignment with Bowtie 2. Nat Methods 9, 357–9 (2012).

14. Li, H. et al. The Sequence Alignment/Map format and SAMtools. Bioinformatics 25, 2078–9 (2009).

15. McKenna, A. et al. The Genome Analysis Toolkit: a MapReduce framework for analyzing next-generation DNA sequencing data. Genome Res 20, 1297–303 (2010).

16. Dunham, I. et al. An integrated encyclopedia of DNA elements in the human genome. Nature 489, 57–74 (2012).

17. Quinlan, A.R. & Hall, I.M. BEDTools: a flexible suite of utilities for comparing genomic features. Bioinformatics 26, 841–2 (2010).

18. Kharchenko, P.V., Tolstorukov, M.Y. & Park, P.J. Design and analysis of ChIP-seq experiments for DNA-binding proteins. Nat Biotechnol 26, 1351–9 (2008).

19. Moran, I. et al. Human beta cell transcriptome analysis uncovers lncRNAs that are tissue-specific, dynamically regulated, and abnormally expressed in type 2 diabetes. Cell Metab 16, 435–48 (2012).

20. Akerman, I. et al. Human Pancreatic beta Cell lncRNAs Control Cell-Specific Regulatory Networks. Cell Metab 25, 400–411 (2017).

21. Dobin, A. et al. STAR: ultrafast universal RNA-seq aligner. Bioinformatics 29, 15–21 (2013).

22. Sherry, S.T. et al. dbSNP: the NCBI database of genetic variation. Nucleic Acids Res 29, 308–11 (2001).

23. Anders, S., Pyl, P.T. & Huber, W. HTSeq--a Python framework to work with high-throughput sequencing data. Bioinformatics 31, 166–9 (2015).

24. Wagner, G.P., Kin, K. & Lynch, V.J. Measurement of mRNA abundance using RNA-seq data: RPKM measure is inconsistent among samples. Theory Biosci 131, 281–5 (2012).

25. Leisch, F. A toolbox for K-centroids cluster analysis. Comput. Stat. Data Anal. 51, 526–544 (2006).

26. Jones E., Oliphant T. & P. P. SciPy: Open Source Scientific Tools for Python. (2001).

27. Ramirez, F., Dundar, F., Diehl, S., Gruning, B.A. & Manke, T. deepTools: a flexible platform for exploring deep-sequencing data. Nucleic Acids Res 42, W187–91 (2014).

28. Dixon, J.R. et al. Topological domains in mammalian genomes identified by analysis of chromatin interactions. Nature 485, 376–80 (2012).

29. Heinz, S. et al. Simple combinations of lineage-determining transcription factors prime cis-regulatory elements required for macrophage and B cell identities. Mol Cell 38, 576–89 (2010).

30. Schmitt, A.D. et al. A Compendium of Chromatin Contact Maps Reveals Spatially Active Regions in the Human Genome. Cell Rep 17, 2042–2059 (2016).

31. Ernst, J. & Kellis, M. ChromHMM: automating chromatin-state discovery and characterization. Nat Methods 9, 215–6 (2012).

32. Roadmap Epigenomics, C. et al. Integrative analysis of 111 reference human epigenomes. Nature 518, 317–30 (2015).

33. Liao, Y., Smyth, G.K. & Shi, W. featureCounts: an efficient general purpose program for assigning sequence reads to genomic features. Bioinformatics 30, 923–30 (2014).

34. Gaulton, K.J. et al. Genetic fine mapping and genomic annotation defines causal mechanisms at type 2 diabetes susceptibility loci. Nat Genet 47, 1415–25 (2015).

35. Bonas-Guarch, S. et al. Re-analysis of public genetic data reveals a rare X-chromosomal variant associated with type 2 diabetes. Nat Commun 9, 321 (2018).

36. Chang, C.C. et al. Second-generation PLINK: rising to the challenge of larger and richer datasets. Gigascience 4, 7 (2015).

37. Benazra, M. et al. A human beta cell line with drug inducible excision of immortalizing transgenes. Mol Metab 4, 916–25 (2015).

38. Gibson, D.G. et al. Enzymatic assembly of DNA molecules up to several hundred kilobases. Nat Methods 6, 343–5 (2009).

39. Ran, F.A. et al. Genome engineering using the CRISPR-Cas9 system. Nat Protoc 8, 2281–2308 (2013).

40. Joung, J. et al. Genome-scale CRISPR-Cas9 knockout and transcriptional activation screening. Nat Protoc 12, 828–863 (2017).

41. Fu, Y., Sander, J.D., Reyon, D., Cascio, V.M. & Joung, J.K. Improving CRISPR-Cas nuclease specificity using truncated guide RNAs. Nat Biotechnol 32, 279–284 (2014).

42. Shuman, S. Novel approach to molecular cloning and polynucleotide synthesis using vaccinia DNA topoisomerase. J Biol Chem 269, 32678–84 (1994).

43. Pulido-Quetglas, C. et al. Scalable Design of Paired CRISPR Guide RNAs for Genomic Deletion. PLoS Comput Biol 13, e1005341 (2017).

44. Sanjana, N.E., Shalem, O. & Zhang, F. Improved vectors and genome-wide libraries for CRISPR screening. Nat Methods 11, 783–784 (2014).

45. Gilbert, L.A. et al. Genome-Scale CRISPR-Mediated Control of Gene Repression and Activation. Cell 159, 647–61 (2014).

46. Horlbeck, M.A. et al. Compact and highly active next-generation libraries for CRISPR-mediated gene repression and activation. Elife 5(2016).

47. Klages, N., Zufferey, R. & Trono, D. A stable system for the high-titer production of multiply attenuated lentiviral vectors. Mol Ther 2, 170–6 (2000).

48. Abraham, A. et al. Machine learning for neuroimaging with scikit-learn. Front Neuroinform 8, 14 (2014).

49. Kuleshov, M.V. et al. Enrichr: a comprehensive gene set enrichment analysis web server 2016 update. Nucleic Acids Res 44, W90–7 (2016).

50. Whyte, W.A. et al. Master transcription factors and mediator establish super-enhancers at key cell identity genes. Cell 153, 307–19 (2013).

51. Parker, S.C. et al. Chromatin stretch enhancer states drive cell-specific gene regulation and harbor human disease risk variants. Proc Natl Acad Sci U S A 110, 17921–6 (2013).

52. Serra, F. et al. Automatic analysis and 3D-modelling of Hi-C data using TADbit reveals structural features of the fly chromatin colors. PLoS Comput Biol 13, e1005665 (2017).

53. Bau, D. & Marti-Renom, M.A. Genome structure determination via 3C-based data integration by the Integrative Modeling Platform. Methods 58, 300–6 (2012).

54. Kremer, K., and Grest, G.S. Dynamics of entangled linear polymer melts: A molecular-­--dynamics simulation. J. Chem. Phys., 5057–12718 (1990).

55. Rosa, A. & Everaers, R. Structure and dynamics of interphase chromosomes. PLoS Comput Biol 4, e1000153 (2008).

56. Rosa, A., Becker, N.B. & Everaers, R. Looping probabilities in model interphase chromosomes. Biophys J 98, 2410–9 (2010).

57. Di Stefano, M., Rosa, A., Belcastro, V., di Bernardo, D. & Micheletti, C. Colocalization of coregulated genes: a steered molecular dynamics study of human chromosome 19. PLoS Comput Biol 9, e1003019 (2013).

58. Di Stefano, M., Paulsen, J., Lien, T.G., Hovig, E. & Micheletti, C. Hi-C-constrained physical models of human chromosomes recover functionally-related properties of genome organization. Sci Rep 6, 35985 (2016).

59. Finch, J.T. & Klug, A. Solenoidal model for superstructure in chromatin. Proc Natl Acad Sci U S A 73, 1897–901 (1976).

60. Harp, J.M., Hanson, B.L., Timm, D.E. & Bunick, G.J. Asymmetries in the nucleosome core particle at 2.5 A resolution. Acta Crystallogr D Biol Crystallogr 56, 1513–34 (2000).

61. Bau, D. et al. The three-dimensional folding of the alpha-globin gene domain reveals formation of chromatin globules. Nat Struct Mol Biol 18, 107–14 (2011).

62. Plimpton, S. Fast Parallel Algorithms for Short-Range Molecular Dynamics. J. Comput. Phys 117, 1–19 (1995).

63. Fiorin, G., Klein, M.L. & Hénin, J. Using collective variables to drive molecular dynamics simulations. Molecular Physics 111, 3345–3362 (2013).

64. Morris, J.H., Meng, E.C. & Ferrin, T.E. Computational tools for the interactive exploration of proteomic and structural data. Mol Cell Proteomics 9, 1703–15 (2010).

65. Shannon, P. et al. Cytoscape: a software environment for integrated models of biomolecular interaction networks. Genome Res 13, 2498–504 (2003).

66. Zhang, Y. et al. Model-based analysis of ChIP-Seq (MACS). Genome Biol 9, R137 (2008).

67. Ahmed, M. et al. Variant Set Enrichment: an R package to identify disease-associated functional genomic regions. BioData Min 10, 9 (2017).

68. Cowper-Sal lari, R. et al. Breast cancer risk-associated SNPs modulate the affinity of chromatin for FOXA1 and alter gene expression. Nat Genet 44, 1191–8 (2012).

69. Gjesing, A.P. et al. Genetic and phenotypic correlations between surrogate measures of insulin release obtained from OGTT data. Diabetologia 58, 1006–12 (2015).

70. Thuesen, B.H. et al. Cohort Profile: the Health2006 cohort, research centre for prevention and health. Int J Epidemiol 43, 568–75 (2014).

71. Drivsholm, T., Ibsen, H., Schroll, M., Davidsen, M. & Borch-Johnsen, K. Increasing prevalence of diabetes mellitus and impaired glucose tolerance among 60-year-old Danes. Diabet Med 18, 126–32 (2001).

72. Johansen, N.B. et al. Protocol for ADDITION-PRO: a longitudinal cohort study of the cardiovascular experience of individuals at high risk for diabetes recruited from Danish primary care. BMC Public Health 12, 1078 (2012).

73. McCarthy, S. et al. A reference panel of 64,976 haplotypes for genotype imputation. Nat Genet 48, 1279–83 (2016).

74. Loh, P.R., Palamara, P.F. & Price, A.L. Fast and accurate long-range phasing in a UK Biobank cohort. Nat Genet 48, 811–6 (2016).

75. Marchini, J. & Howie, B. Genotype imputation for genome-wide association studies. Nat Rev Genet 11, 499–511 (2010).

76. Schwarzer, G. meta: An R package for meta-analysis. R News 7, 40–45 (2007).

77. Nieuwboer, H.A., Pool, R., Dolan, C.V., Boomsma, D.I. & Nivard, M.G. GWIS: Genome-Wide Inferred Statistics for Functions of Multiple Phenotypes. Am J Hum Genet 99, 917–927 (2016).

78. Bulik-Sullivan, B.K. et al. LD Score regression distinguishes confounding from polygenicity in genome-wide association studies. Nat Genet 47, 291–5 (2015).

79. Finucane, H.K. et al. Partitioning heritability by functional annotation using genome-wide association summary statistics. Nat Genet 47, 1228–35 (2015).

80. Wood, A.R. et al. A Genome-Wide Association Study of IVGTT-Based Measures of First-Phase Insulin Secretion Refines the Underlying Physiology of Type 2 Diabetes Variants. Diabetes 66, 2296–2309 (2017).

81. Demontis, D. Discovery of the first genome-wide significant risk loci for ADHD. bioRxiv (2017).

82. Genomes Project, C. et al. A global reference for human genetic variation. Nature 526, 68–74 (2015).

83. Thurner, M. et al. Integration of human pancreatic islet genomic data refines regulatory mechanisms at Type 2 Diabetes susceptibility loci. Elife 7(2018).

84. Sudlow, C. et al. UK biobank: an open access resource for identifying the causes of a wide range of complex diseases of middle and old age. PLoS Med 12, e1001779 (2015).

85. Becerra, M.B. & Becerra, B.J. Disparities in Age at Diabetes Diagnosis Among Asian Americans: Implications for Early Preventive Measures. Prev Chronic Dis 12, E146 (2015).

86. Euesden, J., Lewis, C.M. & O’Reilly, P.F. PRSice: Polygenic Risk Score software. Bioinformatics 31, 1466–8 (2015).

87. Mularoni, L., Ramos-Rodriguez, M. & Pasquali, L. The Pancreatic Islet Regulome Browser. Front Genet 8, 13 (2017).

88. Schofield, E.C. et al. CHiCP: a web-based tool for the integrative and interactive visualization of promoter capture Hi-C datasets. Bioinformatics 32, 2511–3 (2016).

